# Genetic control of the dynamic transcriptional response to immune stimuli and glucocorticoids at single cell resolution

**DOI:** 10.1101/2021.09.30.462672

**Authors:** Justyna A Resztak, Julong Wei, Samuele Zilioli, Edward Sendler, Adnan Alazizi, Henriette E Mair-Meijers, Peijun Wu, Xiaoquan Wen, Richard B Slatcher, Xiang Zhou, Francesca Luca, Roger Pique-Regi

## Abstract

Synthetic glucocorticoids, such as dexamethasone, have been used as treatment for many immune conditions, such as asthma and more recently severe COVID-19. Single cell data can capture more fine-grained details on transcriptional variability and dynamics to gain a better understanding of the molecular underpinnings of inter-individual variation in drug response. Here, we used single cell RNA-seq to study the dynamics of the transcriptional response to glucocorticoids in activated Peripheral Blood Mononuclear Cells from 96 African American children. We employed novel statistical approaches to calculate a mean-independent measure of gene expression variability and a measure of transcriptional response pseudotime. Using these approaches, we demonstrated that glucocorticoids reverse the effects of immune stimulation on both gene expression mean and variability. Our novel measure of gene expression response dynamics, based on the diagonal linear discriminant analysis, separated individual cells by response status on the basis of their transcriptional profiles and allowed us to identify different dynamic patterns of gene expression along the response pseudotime. We identified genetic variants regulating gene expression mean and variability, including treatment-specific effects, and demonstrated widespread genetic regulation of the transcriptional dynamics of the gene expression response.

## Introduction

The immune system exerts its function through a delicate and timely balance of pro- and anti-inflammatory processes. An effective response of the immune cells ensures that pathogens are neutralized; yet, it is equally crucial that inflammatory processes are timely modulated to avoid pathological states, such as systemic inflammation [120], cytokine storm [26], and sepsis [102]. Specificity in recognizing potential pathogens and mounting a proportionate response is important to prevent or resolve infectious diseases; importantly, dysregulation of these processes leads to chronic conditions, including autoimmune, allergic diseases, and asthma. The hypothalamic-pituitary-adrenal axis plays a central role in the balance between pro- and anti-inflammatory processes through regulation of glucocorticoids in the bloodstream. Because of their potent anti-inflammatory effects, synthetic glucocorticoids have widespread pharmacological applications. Steroid anti-inflammatory drugs are used to treat asthma (budesonide) [41], autoimmune diseases (budesonide, dexamethasone, prednisone, methylprednisolone) [38, 72, 73, 107, 109, 123], and several other inflammatory conditions. Dexamethasone is a potent synthetic glucocorticoid that is used to treat leukemia because of its ability to inhibit lymphocyte proliferation [87]. Most recently, dexamethasone has been recognized as the first effective drug to prevent severe complications from COVID-19 [36]. Severe COVID-19 is thought to be a consequence of an exaggerated response of the immune system to SARS-CoV-2 infection [5]. Elevated production of pro-inflammatory cytokines results in tissue damage in severe COVID-19 patients and can ultimately be fatal. By activating anti-inflammatory processes, dexamethasone can prevent this “cytokine storm” and dramatically improve disease outcome [36].

Despite an obviously large contribution of environmental exposures (e.g. allergens, pathogens), genetic and gene-environment interaction effects have a major role in immunological phenotypes [66]. GWAS have successfully identified hundreds of variants associated with autoimmune and allergic diseases risk. For example, genetic associations from GWAS explain up to 33% of the heritability of childhood asthma [86]. Common genetic variants also explain a large fraction of inter-individual variation in the response to pathogens [99]. In addition to large-scale GWAS, *in vitro* studies of the immune transcriptional response to pathogens have identified thousands of variants that contribute to inter-individual variation [2, 7, 12, 39, 79, 89, 97]. These studies have mostly focused on molecular phenotypes, including gene expression, RNA processing, and chromatin accessibility. Interestingly, the same molecular quantitative trait locus (QTL) mapping approaches, have also identified genetic variants that modify an individual’s response to glucocorticoids [63, 65], thus highlighting how genetic variation can regulate the important balance between pro- and anti-inflammatory processes.

A key component of the immune system function is the molecular signaling between the different cell types that compose it. This communication is crucial to enable an accurate response to pathogens. However, it is only with the advent of single cell technology that we can deeply characterize transcriptional responses of individual cell types, while preserving their cross-talk through stimulation of the entire PBMC fraction. Single cell technology also enables the analysis of cell-to-cell transcriptional variability. Gene expression variability plays an important role in biological systems. It has been suggested that in the immune system gene expression variability increases the ability to respond to immune stimuli [31]. *In vitro* studies demonstrate that immune stimuli can modify expression variability in CD4+ T cells [21] and variation of gene expression variability is also observed across individuals in human populations [76, 100].

With the great heterogeneity of function between the different cell types comprising the immune system, it is expected that the genetic effects on gene expression will vary across cell types as well. Indeed, multiple studies have leveraged scRNA-seq technologies to map cell-type-specific eQTLs across the different subpopulations comprising complex tissues. Studies in peripheral blood mononuclear cells [119], fibroblasts [78], tumor samples [59], and pluripotent and differentiating cells [15, 23, 42, 78] found several eQTLs that would have been missed in bulk sequencing approaches due to being active in only one or a few cell types. While in many single-cell studies most of the detected eQTLs were only found in a single cell type [44, 56, 78], this likely overestimates the overall cell-type specificity of genetic effects on gene expression due to incomplete power of eQTL discovery in these datasets [81]. Previous approaches have successfully mapped the genetic determinants of responses to infection, drugs, and other stimuli [2, 7, 12, 25, 37, 39, 47, 48, 49, 51, 61, 62, 64, 77, 79, 89, 97, 104, 125] using bulk RNA-seq. However, these genotype-by-environment (GxE) effects are also likely to be cell-type-specific [28], especially in the context of highly-specialized immune responses, as investigated in recent studies of response eQTL mapping using single cell technology in immune cells exposed to pathogens, bacteria and yeast [81], and influenza A [92]. While different immune stimuli have been studied at single cell resolution; the response to glucocorticoids has not been previously examined.

Here, we used single cell RNA-seq to study the dynamics of the transcriptional response to glucocorticoids in activated PBMCs from 96 African American youths with asthma and the genetic basis of variation in gene expression mean and variability across individuals.

## Results

### Identification of major cell types and gene expression patterns

To systematically characterize genetic determinants of transcriptional response to glucocorticoids at single cell resolution, we studied peripheral blood mononuclear cells (PBMCs) of 96 African American children with asthma. We stimulated the PBMCs with phytohemagglutinin (PHA, a T cell mitogen) or lipopolysaccharide (LPS, a component of the bacterial membrane), and treated with the glucocorticoid dexamethasone for a total of 5 conditions (including the unstimulated control, Figure 1A). We generated a total of 292,394 high quality cells and detected 46,384 expressed genes from 96 individuals across 5 conditions (Table S1 and Figure S1)

**Figure 1:**
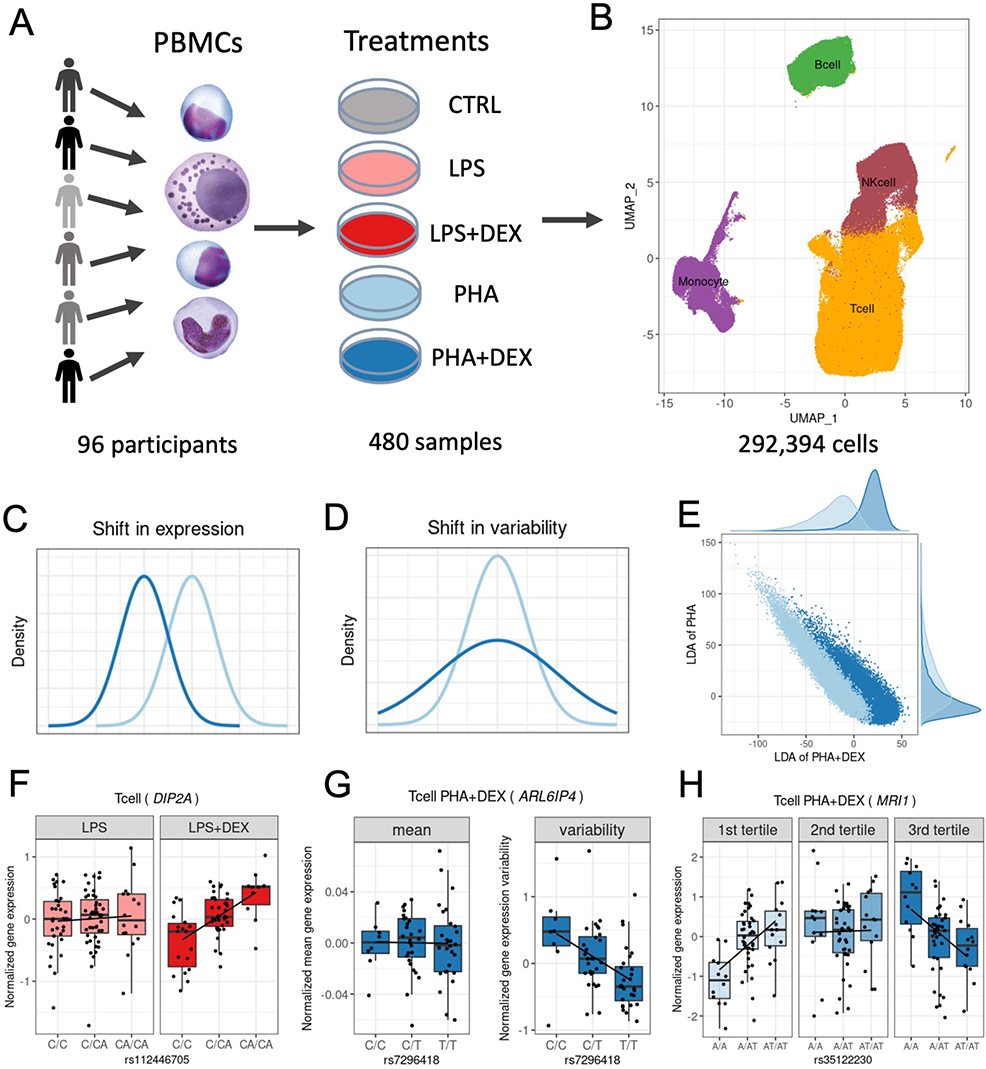
Study overview. **(A)** PBMCs were collected from 96 African American donors with asthma. Cells were stimulated with phytohemagglutinin (PHA), lipopolysaccharide (LPS), and treated with glucocorticoid dexamethasone (DEX) for a total of 5 conditions (including a control), for a total of 480 samples. **(B)** UMAP visualization of the sc-RNA data for a total of 293,394 high quality cells, colored by four major cell types: B cells (green), Monocytes (purple), NK cells (maroon) and T cells (orange). **(C)** Density plot exemplifying changes in mean gene expression between conditions (PHA+DEX vs PHA). **(D)** Density plot exemplifying changes in gene variability between conditions (PHA+DEX vs PHA). **(E)** Scatter plot representing the low dimensional manifolds obtained by diagonal linear discriminant analysis (DLDA) in the T cells treated with PHA and PHA+DEX: x axis denoting the PHA+DEX response pseudotime and y-axis denoting the PHA response pseudotime. **(F)** Example of a LPS+DEX response eQTL for *DIP2A* gene in the T cells. The boxplots depict normalized gene expression levels for the three genotype classes in the two treatment conditions contrasted to identify the response eQTL: LPS+DEX (right panel) and LPS (left panel). **(G)** Example of a variability QTL without an effect on the mean gene expression for the *ARL6IP4* gene in T cells treated with PHA+DEX. The boxplots depict normalized variability (right) and mean (left) gene expression for the three genotype classes. **(H)** Example of a dexamethasone response dynamic eQTL for the *MRI1* gene in the T cells treated with PHA+DEX.

We clustered the cells and identified four major clusters (Figure 1B): B cells, Monocytes, NK cells and T cells. T cells formed the largest cluster with 183,289 cells (about 63%), followed by NK cells (47,824, about 16%), B cells (30,888, about 11%) and Monocytes (30,393, 10%). We confirmed the cell identity using gene expression for cell type-specific genes (Figure S2) While cells are clustered primarily by cell types in the UMAP plot, we did observe partial separation by treatment when analyzing each major cluster separately (Figure S3).

Using this dataset we will explore the more fine-grained details on transcriptional response to immune treatments, including gene expression(Figure 1C), variability (Figure 1D) and dynamics in a short-term span within the treatment (Figure 1E). Then we identify the genetic variants regulating gene expression mean and variability (Figure 1G), including treatment-specific effects (Figure 1F) and also examine genetic regulation of the transcriptional dynamics of the gene expression response (Figure 1H).

### Cell-type specific gene expression response

To identify genes differentially expressed in response to the treatments in each cell type, we first aggregated the count data for each cell type-treatment-individual combination. We considered 4 contrasts in each cell type (Figure S4, Table S10), (1) LPS versus CTRL; (2) LPS+DEX versus LPS; (3) PHA versus CTRL; (4) PHA+DEX versus PHA. We detected a total of 6,571 differentially expressed genes (DEGs) (*q <* 0.1 and fold change *>*1.41 (Figure 2A, Figure S5)). Monocytes showed the strongest gene expression response compared to the other cell types across all treatments (4,460 DEGs). LPS induced the weakest gene expression response in all cell types except monocytes. This was expected because LPS stimulates myeloid cells preferentially [54, 117]. Glucocorticoids induced stronger responses than LPS and PHA across most cell types. In general, we observed that the transcriptional response to glucocorticoids and immune stimuli was highly cell type-specific, such that the majority of DEGs were identified in only one cell type across the four conditions (86% in LPS, 63.7% in LPS+DEX, 66.7% in PHA and 59.6% in PHA+DEX, Figure S6).

When we considered the global patterns of treatment effects on gene expression, we observed that effect sizes of dexamethasone and immune stimuli on gene expression were negatively correlated (Figure 2B), which indicated, as expected, that glucocorticoids reversed the gene expression effects induced by immune stimuli (Figure 2A and Figure S7). The antagonistic effect of glucocorticoids compared to the immune stimuli was observed also at the pathway level using gene ontology enrichment analysis (Figure S8, Figure S9) [e.g., the type I interferon signaling pathway, the response to LPS, the cytokine-mediated signaling pathways, and innate immune response (Figure 2C, Figure S10A-C)]. These immune-related pathways shared across cell types appeared to be the molecular basis of how immune stimuli and glucocorticoids affect the immune system transcriptome in opposite directions. To better characterize these gene expression shifts at the pathway level, we derived a pathway-specific score that combines the gene expression changes of all the genes that belong to that pathway term (see methods for more details). Here, we considered type I interferon signaling pathway as an example (Figure 2C). This pathway was only enriched in genes upregulated for immune stimuli (LPS or PHA) and downregulated for the dexamethasone conditions. We also observed that the score of IFN increased after LPS or PHA stimulation, while the score in the DEX condition was similar to those of unstimulated cells. Type I interferons (IFNs) play important roles in the innate and adaptive immune responses not only to viruses but also to bacterial pathogens[9, 40, 68]. Genes enriched in the type I interferon signaling pathway include *JAK1*, *STAT1*, *STAT2*, *IRF1*, *IRF7*, *IRF8* and *OAS1*. Treatment with LPS or PHA significantly increased expression of genes in the IFN I pathway, while dexamethasone counter-acted this effect. We also observed that genes upregulated in response to immune stimuli or downregulated in response to dexamethasone were enriched in the COVID-19 pathway, and that this enrichment was shared across cell types except monocytes (Figure 2D).

**Figure 2:**
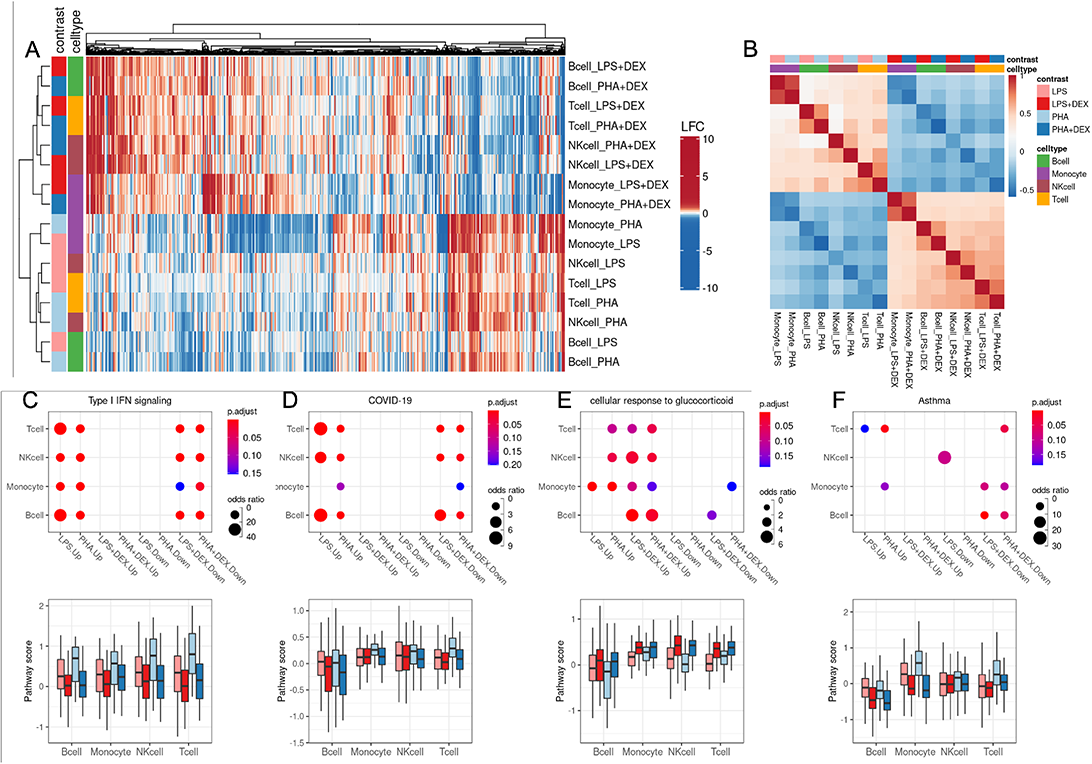
Identification of differentially expressed genes (DEGs). **(A)** Heatmap of log_2_ fold change (LFC) of 6,571 DEGs (column) across 16 conditions (4 contrast *×* 4 cell types, row). **(B)** Heatmap of Spearman correlation of log_2_ fold change (LFC) across 16 conditions (4 cell types *×* 4 contrasts). **(C-F)** Pathway analysis across cell-types and treatment conditions for enrichment in DEG (top) and boxplot for average pathway score (bottom) for four pathways:**(C)** Type I interferon signaling,**(D)** Coronavirus disease-COVID-19,**(E)** glucocorticoid stimulus, and **(F)** asthma.

Moreover, we revealed some dexamethasone-specific pathways, such as the cellular response to glucocorticoid stimulus pathways (Figure 2E, FDR*<* 0.1) and the stress response to metal ion pathways (Figure S10D, FDR*<* 0.1), which are both enriched in genes upregulated in response to treatment with dexamethasone and are consistent with previous findings that glucocorticoids can modulate ion channel activity [34, 115]. In addition, we found that dexamethasone repressed gene expression of the asthma pathway with high cell type-specificity, mainly in B cells (Figure 2F, FDR*<* 0.1). B cells play an important role in the induction of asthma by releasing IgE molecules [96].

### Gene expression variability is modified by glucocorticoids and immune stimuli

Beyond looking at the transcriptional response to immune stimuli for each cell-type separately, single cell data provide an unprecedented opportunity to study gene expression dynamics (mean and variability) in response to treatments. Here, we aimed to investigate changes in gene expression variability induced by treatments. We implemented a method based on the negative binomial distribution to calculate two gene dynamics-related parameters: gene expression mean (*µ*) and dispersion (*ϕ*), based on the work of Sarkar et al. [100]. To avoid the mean-variability dependency and detect treatment effects that affect solely variability, we regressed out any residual mean effects on dispersion and used this mean-corrected dispersion as the measure of gene expression variability in the following analyses (Figure S11). Similar to the above differential expression analysis, we compared gene expression variability in four paired contrasts (i.e., LPS, LPS+DEX, PHA and PHA+DEX) (Figure S12, Table S11). We detected a total of 1,409 differentially variable genes (DVGs, *q <*0.1 and fold change *>*1.41) across cell types and conditions (Figure 3A, Figure S13). We identified the most DVGs in monocytes (910) followed by T cells, NK cells, and B cells. Dexamethasone induced changes in gene expression variability for 50 to 351 genes, with the most DVGs in Monocytes treated with PHA and dexamethasone. When considering gene expression variability, we also observed negative correlations between differential effects of immune stimuli (LPS or PHA) and those of dexamethasone, similar to the findings from differential gene expression analysis (Figure 3B and C, and Figure S14). These findings indicate that dexamethasone dampens the immune response not only by changing gene expression mean but also by altering gene expression variability.

**Figure 3:**
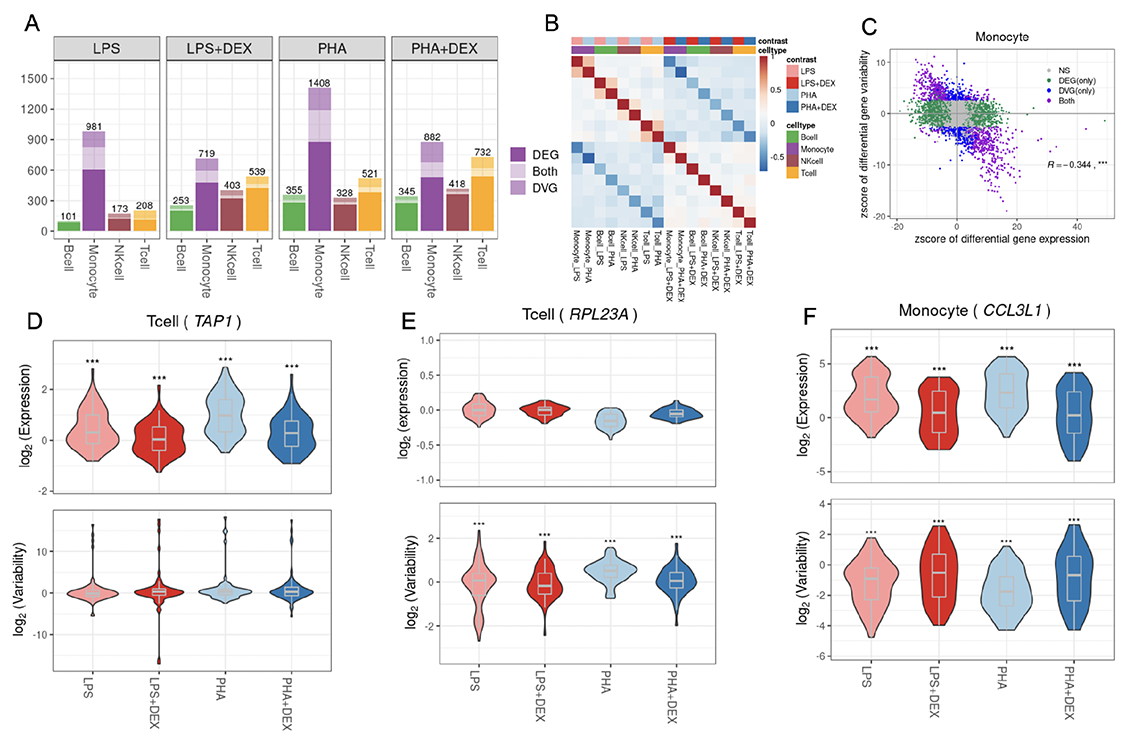
Identification of Differentially variable genes (DVGs). **(A)** Number of the genes with differential mean or variability across cell-types and contrasts: genes with significant changes on gene mean only (dark shading), genes with significant changes on gene variability only (medium shading) and genes with significant changes on both gene mean and gene variability (light shading). **(B)** Heatmap of Spearman correlation of log_2_ fold change (LFC) of gene variability across 16 conditions (4 cell types *×* 4 contrasts). **(C)** Scatterplot of the z score of differential gene expression (x axis) and the z score of differential gene variability (y axis) in monocytes across 4 contrasts. Each dot represents a gene with colors indicating significant (FDR*<*10%) changes on gene mean expression only (green), gene variability only (blue), both (purple) and neither (grey). **(D)** Boxplot of gene expression variability (bottom) and mean expression (top) for the gene (*TAP1*), an example of a gene that undergoes significant changes only to gene expression in response to immune treatments in T cells. **(E)** Boxplot of gene expression variability (bottom) and mean expression (top) of the gene (*RPL23A*), which does not show significant changes in gene expression mean but with significant changes in gene expression variability after immune treatment in T cells. **(F)** Boxplot of gene expression variability (bottom) and mean expression (top) of the gene (*CCL3L1*), which shows changes in both gene expression mean and gene expression variability (bottom panel) after immune treatment in monocytes.

We then considered the effects of the treatments on mean and variability estimated from the same model. We first confirmed that the fold changes identified with this parametric method are highly correlated with those obtained with DESEQ2 (correlation 0.76–0.97, Figure S15). Among the genes only undergoing changes of mean expression levels, *TAP1* was overexpressed in T cells after LPS and PHA stimulation while its expression levels in the DEX conditions were similar to those of unstimulated cells (Figure 3D). In contrast, we did not observe any changes in gene expression variability of *TAP1* across the immune treatments. The *TAP1* and *TAP2* proteins form a protein complex, called transporter associated with antigen processing (TAP), helping transfer peptides from pathogens into the endoplasmic reticulum (ER) [19]. One example of a gene where the treatment modifies expression variability without changes in mean expression was *RPL23A* in T cells following immune stimulation (Figure 3E). This gene encodes a ribosomal protein, which may be one of the target molecules involved in mediating growth inhibition by interferon [43].

When comparing the effect size of treatments on gene expression mean and variability, we observed genes following two major patterns: independent effect on the mean and the variability, or a negative correlation between the effects on the mean and variability. This last pattern was more pronounced in monocytes (Figure 3A, Figure 3C, and Figure S16), for 16%–23% genes across contrasts. One such gene is *CCL3L1* which encodes a cytokine involved in many inflammatory and immunoregulatory processes, such as inhibiting HIV entry by binding to a protein of *CCR5* [17] (Figure 3F). Nevertheless, DEGs and DVGs were largely enriched for similar biological pathways, especially in monocytes and in response to glucocorticoids in all cell types (Figure S17). However, in B cells, NK cells and T cells, DEGs in response to LPS are enriched mostly in different pathways compared to DVGs in response to the same treatment (Figure S17). Shared enriched pathways included cytokine-mediated signaling, type I interferon signaling and innate immune response (Figure S18). We also identified several pathways that only appeared to be enriched in DVGs, including ribosome, translation, mRNA catabolic process, protein localization to the ER, protein targeting to the membrane and the ER (Figure S19).

### Characterizing dynamic responses

Single-cell transcriptomics can also be used to study heterogeneous cell states during important biological processes, such as cell-type differentiation during development [15, 75]. Most algorithms developed to study cell-type differentiation rely on finding a path (i.e. lower dimensional manifold) with cells connecting the initial and final states that can be summarized with a trajectory. The gene expression changes of each cell over the trajectory can be mapped to time, or pseudo-time. Methods to study gene expression dynamics have focused on cell cycle, differentiation trajectory and long duration treatments [13, 15] and are generally unsupervised. To investigate treatment effects in a short time span, we have employed a novel approach based on the diagonal linear discriminant analysis (DLDA) to construct a robust low dimensional representation of the transcriptional response dynamics for each cell-type and treatment. This method is supervised as we know the treatment labels and the two end points in the high dimensional space where most of the cells (i.e., their centroids) will be. Because this method uses supervised learning and has a well known solution, the resulting trajectory (i.e., the line connecting the two centroids) should be more stable and reproducible.

We performed resampling analysis to demonstrate the robustness and stability of the DLDA compared to two existing methods: Monocle 3 [88] and scanpy [127]. We focused on the T-cells treated with PHA+DEX or PHA, as this is the most abundant cell type and the condition with the largest number of DEG. To validate the effectiveness of the novel approach (DLDA), we randomly re-sampled half the dataset for 50 replications. In each replication, we applied our novel approach DLDA and the two other methods to compute response pseudotime.

We do not have the underlying ground truth on how the cells should be exactly ordered according to response pseudotime; yet, we can still assess how expression for each of the responding genes correlates with the estimated pseudotime in each iteration across all cells. Ideally, the cells would be sorted similarly in each iteration, and the correlation of the gene expression with the pseudotime would stay constant. If large variation is observed, it indicates that the pseudotime inference is not robust. We computed the variance of correlation coefficient for each gene across 50 replicates and observed that our method shows the lowest variance (Figure S20A).

For each replicate we can also pick the top 20 most highly correlated genes with the pseudotime variable, which ideally would always be the same. However, across 50 iterations, we identified the following number of most highly correlated genes: 103, 154 and 24 for Monocle, scanpy and DLDA, respectively. This result shows that only for DLDA, the most highly correlated genes remain mostly constant across iterations (24 vs 20 ideally). Not only these genes kept changing for the other methods, but the correlation of these genes with the pseudotime variable was very highly variable (Figure S20B-D). The median value of the Jaccard index for those 20 top genes across all resampling pairs are 0.29, 0.11 and 0.90 for Monocle, scanpy and DLDA, respectively (Figure S20E).

In summary, the resampling analysis demonstrated our approach outperforms previous approaches in stability and accuracy of pseudo-time and the consistency of the pseudo time-determined genes.

Then we applied our novel approach to calculate 4 DLDA axes corresponding to 4 contrasts for each cell-type as follows: LPS (compared to unstimulated), PHA (compared to unstimulated), LPS+DEX (compared to LPS) and PHA+DEX (compared to PHA). As expected, we observed that the first two DLDA axes were able to distinguish cells treated with immune stimuli (LPS or PHA) from the unstimulated group across all cell types (Figure 4A,B and Figure S21). We observed the clearest separation in monocytes, in line with the strongest gene expression response to stimuli found in monocytes. The latter two DLDA axes clearly divided the cells into two groups (Figure 4A,B and Figure S21): one treated with DEX and the other treated with immune stimuli. Based on these observations, we defined the first two DLDA axes as immune-stimuli pseudotime and the third and fourth axes as the dexamethasone response pseudotime.

**Figure 4:**
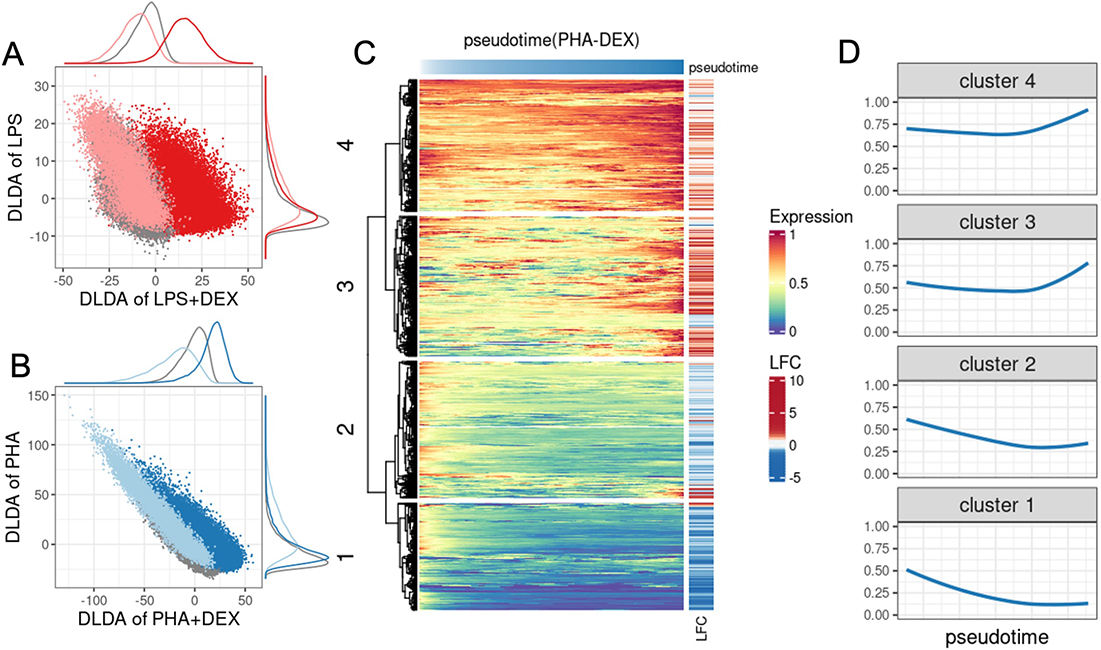
Characterization of dynamic changes along immune response pseudotime. **(A)** Scatterplot of the DLDA pseudotime for LPS+DEX (x axis) and the DLDA pseudotime for LPS (y axis) in T cells. Each dot represents a cell, with the color indicating the treatment condition: grey for unstimulated, light red for LPS and dark red for LPS+DEX. **(B)** Scatterplot of the DLDA pseudotime for PHA+DEX (x axis) and the DLDA pseudotime for PHA in T cells. Each dot represents a cell, with the color indicating the treatment condition: grey for unstimulated, light blue for PHA and dark blue for PHA+DEX. **(C)** Heatmap of relative gene expression for 1,617 DEGs (rows) averaged over windows containing 10% of T-cells (columns) sliding at a step of 0.1% of T cells along the PHA+DEX response pseudotime. For each gene, gene expression is expressed relative to the highest expression across pseudotime. Red color denotes high expression value, yellow denotes medium expression value and blue denotes low expression value. The LFC column on the right indicates the log_2_ fold changes for each DEG between PHA and PHA+DEX. **(D)** Dynamic patterns of relative gene expression averaged across all genes within each cluster using the same sliding window approach as in C using the same sliding window approach as in C.

We next sought to analyze gene expression dynamic changes along the response pseudotime within each treatment for each cell type separately and identified different dynamic patterns of gene expression. For example, in T cells treated with PHA+DEX, four distinctive gene expression dynamic patterns were identified among 1,617 DEGs that were differentially expressed in PHA+DEX (Figure 4C). Expression of genes within clusters 1 and 2 decreased along dexamethasone pseudotime while genes in cluster 1 are relatively lower expressed than those of cluster 2. Most of them were downregulated in the DEX condition compared to PHA (Figure 4D). By contrast, the genes within clusters 3 and 4 tended to first slightly decrease in expression and then burst at a later point. Most of the genes constituting these clusters were upregulated. In line with above enrichment analysis for DEGs, the genes within cluster 1 and 2 were mostly enriched in immune-related pathways, such as, response to LPS (Figure S22), while genes from cluster 4 were preferentially enriched in metal ion-related pathways, such as stress response to copper ion, cellular response to zinc ion, and cell response to cadmium ion (Figure S22). Cluster 4 thus captures the dynamic response that is specific to dexamethasone. Collectively, this novel supervised approach (DLDA) provides a more fine-grained resolution of the dynamics of gene expression response to immune stimuli.

### Genetic effects on gene expression across cell types and treatments

To dissect the genetic contributions to the diversity of immune response dynamics, we mapped the *cis* regulatory variants affecting gene expression response to immune stimuli across cell types and treatment conditions. First, we performed *cis*-eQTL mapping using FastQTL [83] in each cell type-condition combination separately, and discovered 5,190 unique genes with genetic effects on gene expression (eGenes, 10% FDR, Table S3, S12, Figure S23). Genetic effect sizes discovered across cell types and conditions were significantly correlated with those discovered in bulk leukocyte gene expression data on a larger sample that included the same individuals analyzed here [93] (Figure S24A). Expectedly, genetic effects on gene expression best correlated with those from bulk data from T cells as these are the most abundant PBMC type, thus contributing most to the eQTL signal discovered in bulk data.

We next sought to investigate if the genetic effects were shared or context-specific in PBMCs. For this analysis, we used multivariate adaptive shrinkage (mash) method [118] on eQTL summary statistics. Mash uses an empirical Bayes method to estimate patterns of similarity in effect sizes across conditions and then uses these patterns to improve accuracy of effect size estimates while also increasing power. Here, Mash takes advantage of parallel measures of genetic effects across the different cell types and conditions, thus improving the power of eQTL discovery. Indeed, we found approximately an order of magnitude more eGenes in each condition at 10% LFSR (Figure 5A, Figure S25, Table S3, S13, S14) compared to condition-by-condition analysis at 10% FDR using FastQTL. The increase in power is due to a high degree of sharing of genetic effects across conditions. Borrowing information across conditions also allows us to increase the precision of genetic effect size estimation. Correlations between genetic effects estimated in each condition and in bulk leukocyte gene expression data are all close to 0.6 (Spearman correlation, p-value*<* 0.01, Figure S24B), which is a marked improvement over correlations between FastQTL estimates in single-cell and bulk data (Figure S24A). Additionally, 41% of the eGenes we discovered were novel compared to eGenes discovered in the bulk leukocyte dataset (Figure S26).

**Figure 5:**
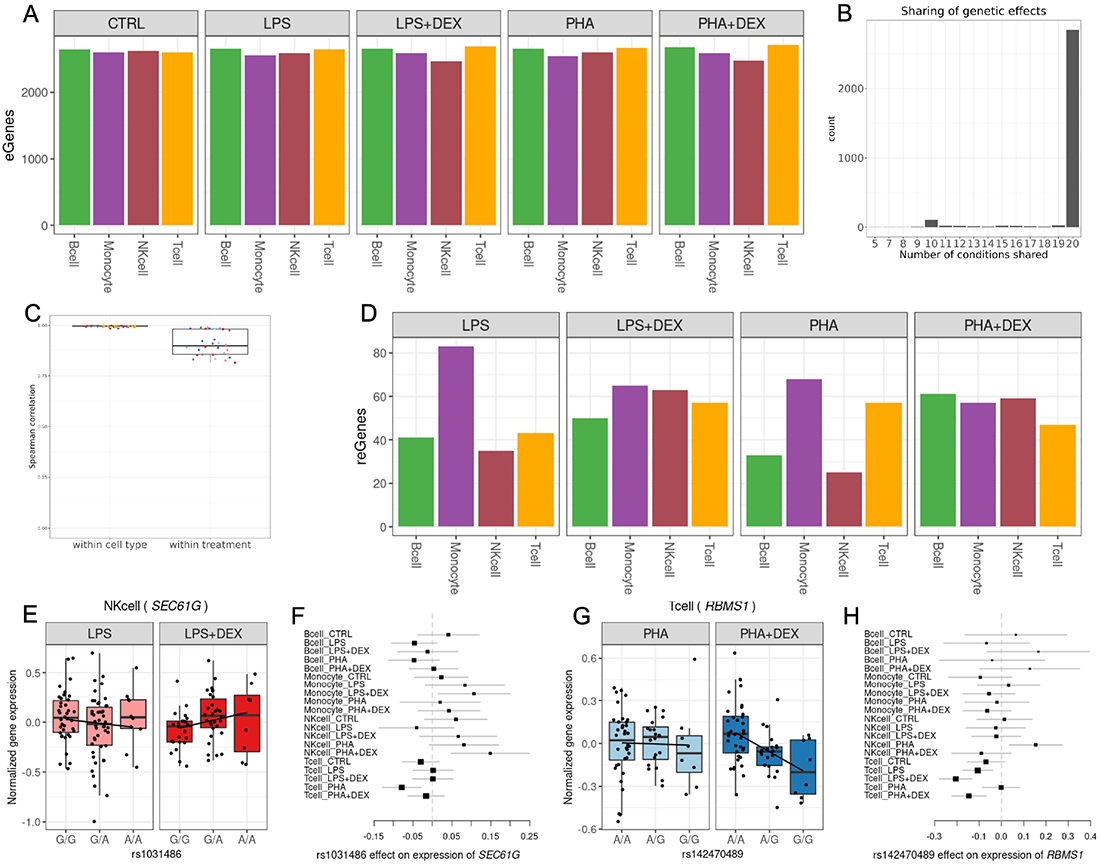
Genetic effects on gene expression. **(A)** Barplot of the number of genes with eQTLs (eGenes) in each condition significant in multivariate adaptive shrinkage analysis (10% LFSR). **(B)** Barplot of the number of eGenes (y axis) with genetic effects in the same direction across the number of conditions shared (x axis). **(C)** Boxplot of Spearman correlations between significant genetic effects on gene expression for each pair-wise comparison of two treatment conditions within each cell type, and for each pair-wise comparison of two cell types within each treatment condition, color reflects cell types (as in Figure 5A) or treatment: control (grey), LPS (pink), LPS+DEX (red), PHA (light blue), and PHA+DEX (dark blue). **(D)** Barplot of the number of genes with response eQTLs (reGenes). **(E)** Boxplot of normalized gene expression for *SEC61G* gene in NK cells treated with LPS(left panel) and LPS+DEX (right panel) across the three genotype classes of the reQTL rs1031486 (x axis). **(F)** Forest plot of the estimated genetic effect for rs1031486 minor allele on the expression of the *SEC61G* gene across all conditions (values reflect slopes as in 5E). Each line represents the 95% confidence interval around the estimate. **(G)** Boxplot of normalized gene expression for *RBMS1* gene in T cells treated with PHA (left) and PHA+DEX (right), across the three genotype classes of the reQTL rs142470489 (x axis). **(H)** Forest plot of the estimated genetic effect of rs142470489 minor allele on the expression of the *IRF3* gene across all conditions (values reflect slopes as in 5G). Each line represents the 95% confidence interval around the estimate.

Genetic effects on gene expression were in the same direction across all 20 conditions for the majority of eGenes (2,832) (Figure 5B, Figure S27, Figure S28, Figure S29). A subset of eGenes (84) shared genetic effects in the same direction across ten conditions (Figure 5B), representing a subdivision between NK cells and T cells in one group and monocytes and B cells in the other group. We observed higher correlations of genetic effect sizes across treatment conditions within the same cell type than across cell types for the same treatment condition (Figure 5C, Figure S30, Figure S31), with higher within cell-type correlations for monocytes and B cells (Spearman *ρ >* 0.99) compared to NK cells (Spearman *ρ >* 0.95) and T cells (Spearman *ρ >* 0.97). A similar result was observed when considering eGenes that were shared across pairs of conditions, with the cell-type specific eGenes (6–51%) representing a larger fraction than the treatment-specific eGenes (1–5%) in the pairwise comparisons (Figure S29). This result highlights the stronger contribution of cell type effects compared to environmental perturbations on genetic regulation of gene expression, which we have observed previously in other cell types [28].

We analyzed pairs of treatment and control conditions within each cell type to discover response eQTLs (reQTLs). Overall, we found 328 genes with reQTLs (reGenes, Figure 5D). For the treatments including DEX the number of reQTLs was roughly similar across cell-types. In contrast, among the different cell types stimulated with LPS or PHA, the largest number of reGenes were found in monocytes (LPS and PHA) and T-cells (PHA). eGenes were enriched for differentially expressed genes across all contrasts and for differentially variable genes in 14/16 contrasts (Fisher’s test p*<* 0.05, Table S4). reGenes were enriched for differentially expressed genes only in B cells treated with LPS+DEX and for differentially variable genes in three conditions (B cell LPS+DEX, Monocyte PHA+DEX and NK cell PHA+DEX, Fisher’s test p*<* 0.05, Table S5).

Dexamethasone reGenes varied between 47 in T cells treated with PHA+DEX and 65 in Monocytes treated with LPS+DEX. For example, we found a reQTL for the *SEC61G* gene in NKcells, where the A allele was associated with higher expression after dexamethasone treatment.(Figure 5E,F, Figure S32). SEC61G is a subunit of the SEC61 translocon responsible for translocation of newly-synthesized proteins into the ER. This complex is required for replication of flaviviruses in human cells [101]. While *SEC61G* is primarily known as a proto-oncogene in several types of cancers [29, 55, 58, 71], it has previously been found to be upregulated in NK cells upon IL2 stimulation [18]. To understand the phenotypic relevance of the reGenes, we considered genes associated with diseases in TWAS and found that 61 reGenes have previously been implicated in immune diseases (Table S15). For example, rs142470489 is a reQTL for the *RBMS1* gene which encodes a member of a small family of proteins that bind single stranded DNA/RNA, involved in the process of DNA replication, gene transcription, cell cycle progression and apoptosis. TWAS implicated *RBMS1* in inflammatory bowel disease [133]. The G allele at rs142470489 increases the dexamethasone immunomodulatory effect on expression of *RBMS1* in T cells treated with PHA+DEX (Figure 5G,H, Figure S33).

### Genetic effects on gene expression variability

Single cell data combined with our new modeling approach allowed us to quantify changes across contexts and individuals in gene expression variability independently of the changes in the mean gene expression. To identify genetic effects on gene expression variability that account for inter-individual variation, we used a QTL mapping approach to discover variability QTLs (vQTLs). We discovered 123 genes with variability QTLs (vGenes), with highest numbers of vQTLs detected in T cells across all conditions (2,989-3,230, corresponding to 98-107 genes) (10% LFSR, Figure 6A, Figure S34A, Table S16, Table S6, Table S17, and Table S18). vGenes were highly enriched for ribosomal proteins (GO enrichment for “cytosolic ribosome” p-value= 1.6 *×* 10*^−^*^43^). However, overall genetic effects across contexts for vQTLs were less correlated than for eQTLs (Figure S35). We observed the highest correlations of genetic effects across treatment conditions within monocytes compared to the other cell types (Figure S36A), and an overall decrease in pairwise correlations across cell types in treatment conditions compared to the control (Figure S36B). Similarly to eGenes, genetic effects on gene expression variability were shared in the same direction for most of the vGenes (98) across all cell types and treatment conditions (Figure 6B, Figure S37). In contrast to eGenes, for vGenes we observed similar correlations of genetic effect sizes across treatment conditions within the same cell type than across cell types for the same treatment condition (Figure 6C). This is confirmed also when considering sharing of genetic effects across conditions (0.05-0.32 for cell-type specific vGenes and 0.06-0.34 for treatment-specific vGenes) (Figure S38).

**Figure 6:**
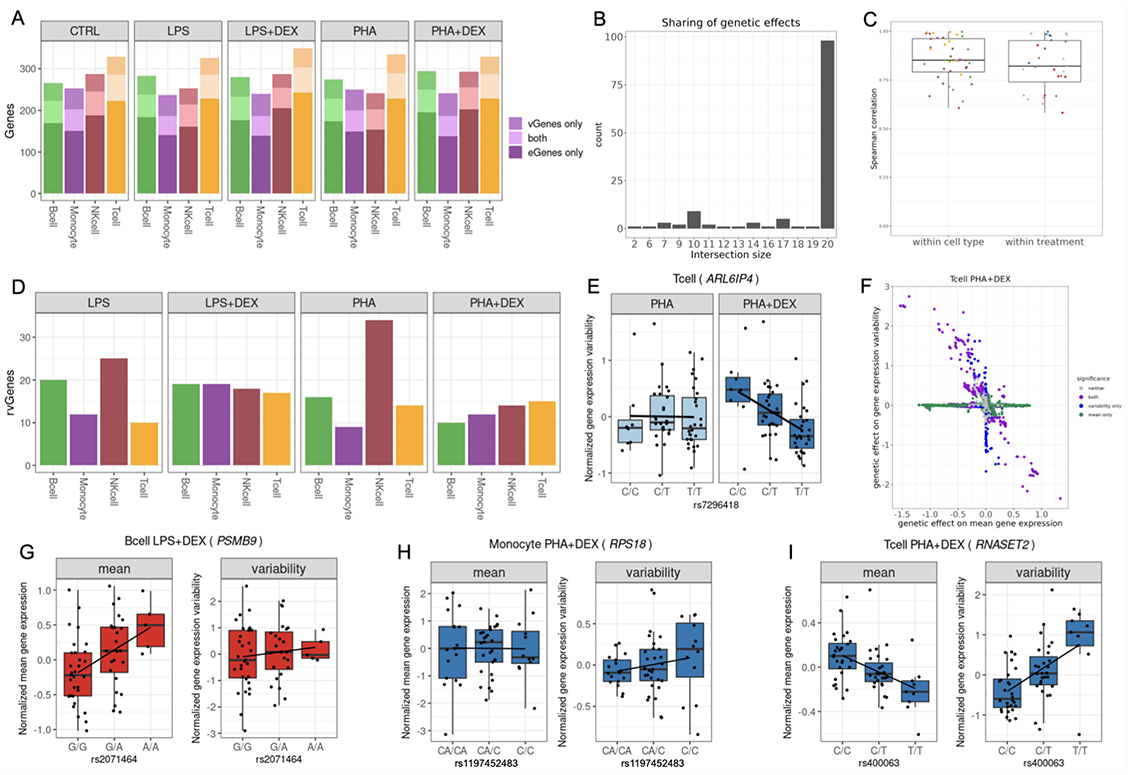
Genetic effects on gene expression variability. **(A)** Barplot of the number of genes with eQTLs (eGenes) for which we did not detect genetic effects on variability (dark shading), number of genes with vQTLs (vGenes) for which we did not detect any genetic effects on gene expression (medium shading), and genes with both eQTLs and vQTLs (light shading). **(B)** Boxplot of the Spearman correlations between significant genetic effects on gene expression variability for each pair-wise comparison of two treatment conditions within each cell type, and for each pair-wise comparison of two cell types within each treatment condition, color reflects cell types (as in Figure 6A) or treatment: control (grey), LPS (pink), LPS+DEX (red), PHA (light blue), and PHA+DEX (dark blue). **(C)** Barplot of the number of vGenes (y axis) with genetic effects in the same direction across the number of conditions shared (x axis). **(D)** Barplot of the number of genes with response vQTLs (rvGenes). **(E)** Boxplot of normalized gene expression variability for *ARL6IP4* gene in T cells treated with PHA(left panel) or PHA+DEX (right panel) across the three genotype classes of the vQTL rs7296418 (x axis). **(F)** Scatterplot of genetic effects on mean gene expression (x axis) versus genetic effects on gene expression variability (y axis) for T cells treated with PHA+DEX; color represents genetic variants with significant effects on mean gene expression only (green), gene expression variability only (blue), both (purple), and neither (grey). **(G)** Boxplot of normalized mean gene expression (left panel) and normalized gene expression variability (right panel) for *PSMB9* gene in B cells treated with LPS+DEX with significant genetic effect of rs2071464 on mean gene expression, but no effect on gene expression variability. **(H)** Boxplot of normalized mean gene expression (left panel) and normalized gene expression variability (right panel) for *RPS18* gene in Monocytes treated with PHA+DEX with significant genetic effect of rs1197452483 on gene expression variability, but no effect on mean gene expression. **(I)** Boxplot of normalized mean gene expression (left panel) and normalized gene expression variability (right panel) for *RNASET2* gene in T cells treated with PHA+DEX with significant genetic effect of rs400063 on both mean gene expression, and on gene expression variability.

To identify response vQTLs, we performed a direct comparison of genetic effect sizes across conditions in each cell type. We discovered 61 unique variability response genes (rvGenes, Figure 6D), including a total of 47 for dexamethasone response. We identified 16 rvGenes that were previously implicated in immune diseases via TWAS. For example, *ARL6IP4* is associated with multiple sclerosis and allergic rhinitis, and we found the T allele at rs7296418 to decrease gene expression variability of this gene in T cells treated with PHA+DEX, but not in PHA condition (Figure 6E, Figure S39). ARL6IP4 may have a role in splicing regulation [84], and was reported to inhibit splicing of the pre-mRNA of Herpes singlex virus (HSVI) [53].

### Comparison between genetic effects on gene expression mean and variability

To compare eQTLs and vQTLs, we considered the genetic effects on mean and variability estimated from the same model (Figure S34B, Table S19, Table S7, Table S20, and Table S21). We identified three distinct patterns (Figure 6F, Figure S40): genetic variants affecting gene expression levels only (7,155-11,314 gene-SNP pairs), genetic variants affecting gene expression variability only (1,709-2,128 gene-SNP pairs), and genetic variants affecting both gene expression levels and variability (669-1,207 gene-SNP pairs). Genetic effects in this last category were negatively correlated with opposite signs for eQTLs and vQTLs (Pearson correlation *−*0.25 - *−*0.8, p-value*<* 0.05).

For example, rs2071464 is an eQTL for the *PSMB9* gene in B cells stimulated with LPS+DEX, but we found no genetic effect of this variant on gene expression variability (Figure 6G, Figure S41). The *PSMB9* located in the MHC class II region encodes a member of the proteasome B-type family, which is a 20S core beta subunit of the proteasome. Several studies indicated the proteasome plays a critical role in cardiovascular diseases [98, 124], inflammatory response and autoimmune diseases [45]. The subunit of proteasome encoded by *PSMB9* was found to be involved in the process of infectious diseases [105], autoimmune diseases [46] and oncology [122]. Previous TWAS study revealed that expression of *PSMB9* has also been implicated in a number of autoimmune disorders including asthma [133]. We found the C allele at the genomic position rs1197452483 to increase the gene expression variability of *RPS18* gene in monocytes treated with PHA+DEX, without any effect on mean gene expression levels (Figure 6H, Figure S42). This gene encodes a ribosomal protein that is a component of the 40S subunit, and its expression has been implicated in rheumatoid arthritis, multiple sclerosis, and psoriasis [133]. *In vitro* silencing of this gene leads to decreased rate of viral infection [106, 112]. Genetic polymorphism at rs400063 confers effects on both gene expression mean and variability of the *RNASET2* gene in the T cells treated with PHA+DEX, but in opposite directions (Figure 6I, Figure S43). RNASET2 plays a crucial role in innate immune response by recognizing and degrading RNAs from microbial pathogens that are sensed by *TLR8* [30]. Expression of *RNASET2* has been implicated in Crohn’s disease, inflammatory bowel disease, and rheumatoid arthritis [133] via TWAS. eQTLs were likely to be also vQTLs (OR=10.45, p-value*<* 2.2 *×* 10*^−^*^16^); however, for the majority of vQTLs, we did not detect significant genetic effects on gene expression levels. vGenes were significantly enriched for genes with differential mean expression for six of the 16 conditions considered (LPS, LPS+DEX and PHA in B cells and T cells, Fisher’s test p-value*<* 0.05, Table S8). vGenes were also enriched for genes with differential variability for seven contrasts (LPS and PHA in B cells and T cells; LPS and PHA+DEX in NK cells; and PHA+DEX in Monocytes; Fisher’s p-value *<* 0.05).

### Mapping of dynamic eQTL using response pseudotime

While reQTLs represent genetic variants that modulate the average response to treatments across cells, our dynamic analysis uncovered more complex cellular response trajectories which we have quantitatively analyzed through DLDA. Through the characterization of the dynamic response patterns above, we observed that cluster 4, representing late response genes, in T cells treated with PHA+DEX was enriched for genes with genetic effects on gene expression (OR= 2.53, Fisher’s exact test p-value=3.5 *×* 10*^−^*^3^). This indicated the potential role of genetic effects in the regulation of dynamic gene expression changes. We therefore systematically investigated genetic variants that modulate these response trajectories (dynamic eQTLs) (Figure S44) and discovered a total of 3,899 dynamic eQTLs corresponding to 588 eGenes across all the conditions (Table S9, Table S22). The largest number of dynamic eGenes were identified in T cells, with dynamic eGenes in the immune-stimuli pseudotime (244 for LPS and 225 PHA) and for dexamethasone response pseudotime (18 LPS+DEX and 54 for PHA+DEX). We also observed that dynamic eQTLs are enriched among eGenes and response eGenes (Figure S45, Figure S46); yet a large number of them are novel. This pattern is particularly noticeable in T cells (Figure S47A, B, Fisher’s exact test p-value*<* 0.01). To investigate whether these dynamic eGenes are potentially causal genes for immune diseases, we tested their overlap with immune-related genes identified from TWAS ([133]) and revealed a total of 92 dynamic eGenes (Table S9) in T cells which have previously been implicated in 22 immune-related diseases in TWAS [133].

To illustrate genetic effects on dynamic response patterns along pseudotime, we focused on T cells treated with PHA+DEX. We showed a dynamic eQTL for *ABO*, which was previously reported to be associated with allergic rhinitis in TWAS [133]. This gene encodes an enzyme responsible for glycosyltransferase activity to determine the blood group in an individual. This gene has been linked to the strongest signal for severe coronavirus disease 2019 (COVID-19) by genome-wide association studies [80, 103]. We discovered a dynamic eQTL modulating *ABO* gene expression along dexamethasone response pseudotime such that *ABO* expression rapidly increased in heterozygous individuals along DEX pseudotime but remained relatively stable in individuals homozygous for the alternate allele (Figure 7A). As a result, we observed a negative genetic effect of the alternate allele on *ABO* gene expression in the mid and late pseudotime. Another example was a dynamic eQTL for the *HLA-DRB5* gene which has been implicated in 14 immune-related diseases, including asthma. The protein product of *HLA-DRB5* forms the beta chain of MHC II. MHC II molecules are used by antigen-presenting cells to display foreign peptides to the immune system to trigger the body’s immune response, but have also been found on activated T cells where they function as signal-transducing receptors [35]. We observed divergent genetic regulation effects on this gene across the three pseudotime tertiles: negative genetic effects on this gene in early pseudo-time, no genetic effects in mid and positive genetic effects in late pseudotime (Figure 7B). We revealed very distinct patterns of dynamic changes in *HLA-DRB5* gene expression along pseudotime for the three genotype classes such that the gene expression sharply decreased in individuals homozygous for A allele at rs116611418, displayed the least gene expression changes in heterozygous individuals throughout the pseudotime and rapidly increased in the late pseudotime in individuals homozygous for the alternate allele (Figure 7B). Additionally, we discovered a dynamic eQTL modulating *GADD45G* expression along dexamethasone response pseudotime such that *GADD45G* expression appeared to be extinguished most rapidly in individuals homozygous for the G allele at rs11265809, with heterozygous individuals following an intermediate slope, and carriers of two A alleles displaying the least changes to *GADD45G* expression along the pseudotime (Figure 7C). As a result, we observed a genetic effect on *GADD45G* expression only in the first tertile of the dexamethasone response pseudotime. The protein encoded by this gene responds to environmental stresses [113, 130]. This gene was reported to be associated with asthma and systemic lupus erythematosus in the TWAS studies [133].

**Figure 7:**
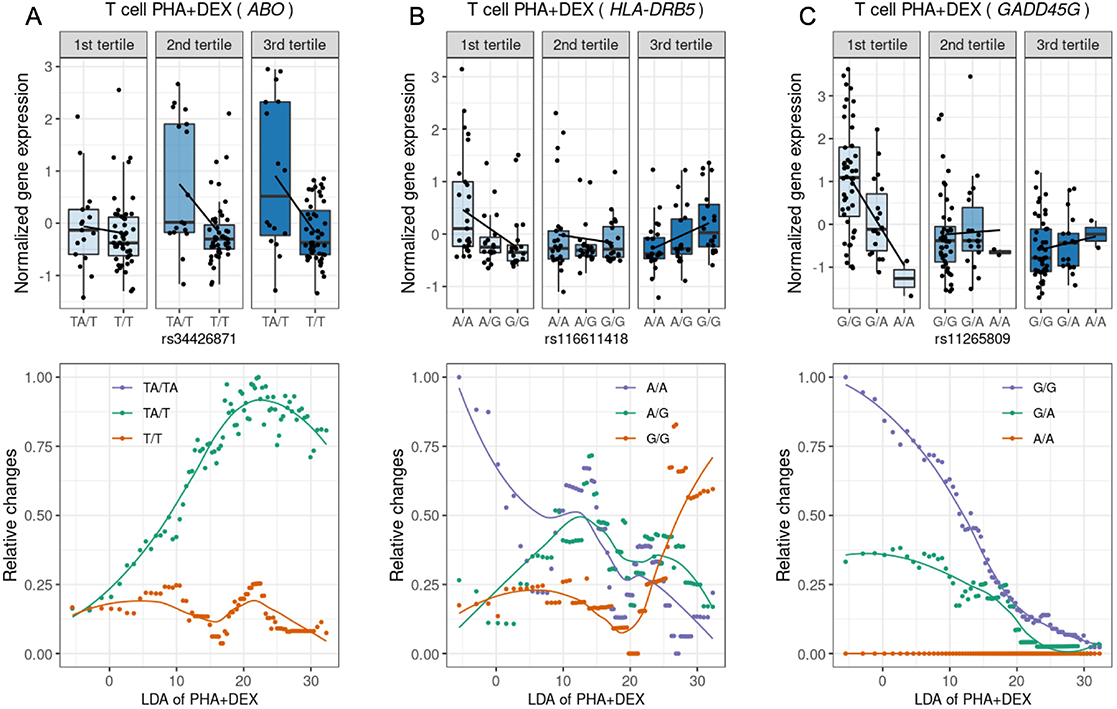
Identification of dynamic eQTLs along immune response pseudotime. For the following genes: **(A)** *ABO*, **(B)** *HLA-DRB5*, and **(C)** *GADD45G*; the boxplots on the top represent normalized gene expression in T cells treated with PHA+DEX for each DLDA pseudotime tertile and genotype; and the bottom plots represent the smoothed dynamic gene expression changes along response trajectory pseudotime for each genotype separately (trendlines are fit using locally estimated scatterplot smoothing).

### Biological insight into the mechanism underlying asthma

Glucocorticoids are the common drugs used to treat asthma and other autoimmune diseases by suppressing immune response. We wanted to explore how genes associated with asthma respond to these immune treatments. We considered 78 asthma-associated genes identified by a Probabilistic Transcriptome Wide Association Study (PTWAS) by integration of eQTLs from whole-blood tissue in GTEx with asthma GWAS cohorts [133]. This type of analysis connects disease risk with gene expression levels. We identified a total of 30 genes that were differentially expressed in at least one condition in our data and were also associated with asthma (Figure 8A).

**Figure 8:**
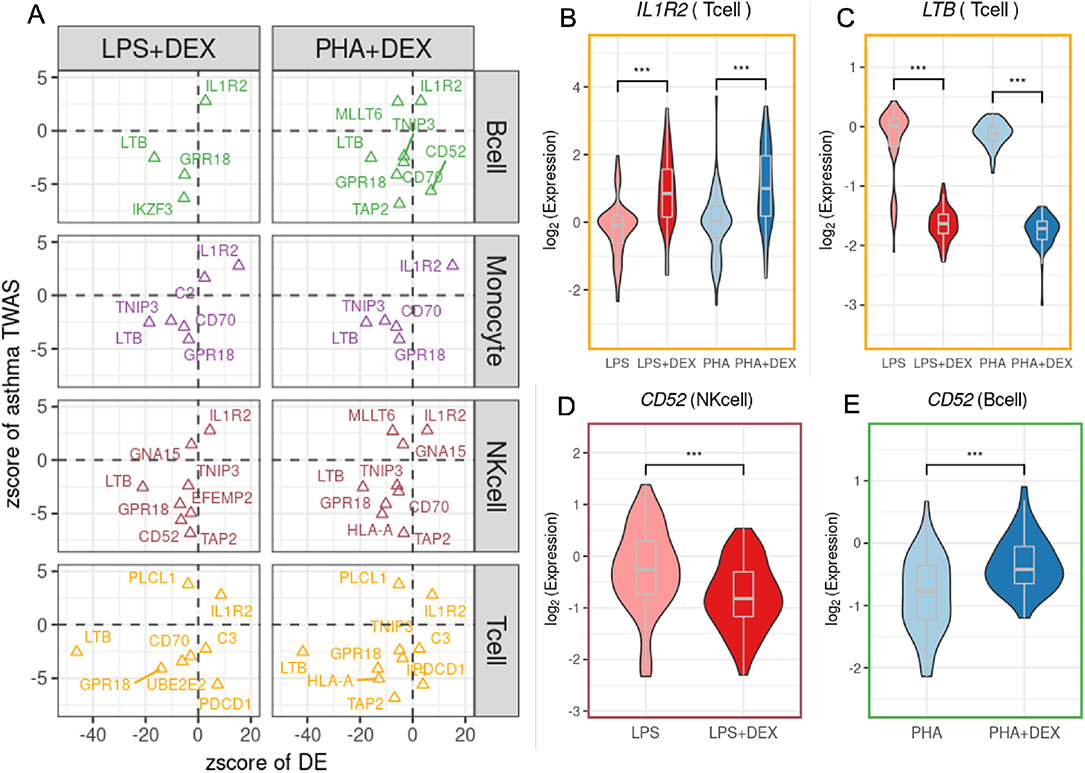
The roles of treatments in the asthma-determined genes: **(A)** Scatter plots of the z-score of dexamethasone effects (x-axis) on 18 asthma-associated genes against the z-score from PTWAS (y-axis). Each dot color represents cell-type. **(B-E)** Example genes: **(B)** *IL1R2* in T-cells, **(C)***LTB* in T-cells and **(D and E)***CD52* in NK-cells and B-cells. Each dot is an individual.

Examples include *IL1R2* (Interleukin 1 receptor type 2), encoding a cytokine receptor that belongs to the interleukin 1 receptor family [4, 85]. This protein binds interleukin alpha (IL1A), interleukin beta (IL1B), and interleukin 1 receptor, type I(IL1R1/IL1RA), and acts as a decoy receptor that inhibits the activity of its ligands. This gene is upregulated in response to dexamethasone in all cell types.(Figure 8B).

The expression of *LTB* is significantly repressed by glucocorticoids via inhibiting the activity of the upstream regulatory factor *NF-κB* across cell-types (Figure 8C) [121]. *LTB* is downregulated in response to dexamethasone in all cell types and is also an eGene in T cells treated with Dexamethasone (PHA-Dex). The protein encoded by Lymphotoxin-beta (*LTB*) forms a heterotrimer with *LTA* on the surface of lymphoid cells [11], involved in a variety of inflammatory responses [131]. This gene is also known as Tumor Necrosis Factor C (TNFC) which was found to decrease gene expression in children with bronchopneumonia following treatment with methylprednisolone in combination with azithromycin [129].

*CD52* was functionally verified to be a potential candidate causal gene linked to asthma in a previous study [33]. In vivo experiment validated that anti-CD52 antibody reduced the inflammatory response in lymphoid cells and airway epithelium [33]. *CD52* displays a cell-type specific transcriptional immune response to dexamethasone, with increased levels in B cells (Figure 8D) but decreased levels in NK cells (Figure 8E) after DEX treatment.

## Discussion

Single-cell technologies allow the study of all aspects of the gene expression response to stimuli — mean, variability and dynamics — across different cell types. Here, we used single cell RNA-seq to study the dynamics of the transcriptional response to glucocorticoids in activated PBMCs. Using this system, we demonstrated that both gene expression mean and variability can be modified by glucocorticoids and immune stimuli. Further, we identified the genetic variants regulating gene expression mean and variability across treatments. We introduced a novel computational approach which allowed us to track the short-term transcriptional response dynamics to treatments at single-cell resolution. Using this method, we identified different dynamic patterns of gene expression along the response pseudotime and demonstrated widespread genetic regulation of these dynamics.

Studies of glucocorticoid response in tightly-controlled experimental settings of individual cell types found varied effects of glucocorticoids on different cells [52, 60, 90, 91, 114, 116, 128]. However, glucocorticoid response could not be studied in the context of the cellular complexity of the immune system prior to the advent of single-cell technologies. Here, we studied the gene expression response of four cell types to glucocorticoids within activated PBMCs. We revealed an opposite pattern of effect on gene expression between immune stimuli and dexamethasone in all cell types analyzed, which was evident both at the gene level and pathway level. These pathways with antagonistic patterns mainly contained immune-related pathways, such as IFN, response to lipopolysaccharide, cytokine-mediated signaling, and innate immune response, mostly shared across cell types, in line with previous findings [82]. Our study also highlighted the specialization in immune response of the different cell types as the majority of DEGs were identified within only one cell type. We observed these cell-type-specific patterns on the pathway level, such as asthma pathway mainly enriched in B cells and T cells, and granulocyte activation mostly enriched in monocytes. Our results are obtained in a childhood asthma cohort and may not be representative of gene expression levels in healthy children or adult populations; yet, the fundamental genetic and biological findings can be most likely extended to healthy individuals. Most importantly, our results provide a unique resource to investigate the molecular underpinnings of immune diseases such as childhood asthma. Many of the differentially expressed genes overlap with genes associated with asthma in TWAS and also well characterized biomarkers. Interestingly, for CD52 we can identify glucocorticoids acting with opposite effects in two different cell-types.

Previous studies implied the important role of gene expression variability in many aspects, such as developmental states [14, 94], T cell differentiation [21], and aging [67]. Our systematic study of gene expression variability in response to immune treatments revealed major effects of immuno-modulators on gene expression variability, with a total of 1,409 DVGs across cell types and contrasts. We observed the same pattern of negative correlation of effects of immune stimuli and dexamethasone on gene expression variability as for gene expression mean levels. Similarly, these DVGs tended to be enriched in immune-related pathways in an antagonistic way similar to the expression analysis. These findings demonstrated that immune treatments affected not only gene expression but also altered the transcriptional variability of genes involved in the immune system. Although we regressed out the effect of mean on gene expression variability, for a subset of genes, we observed a negative correlation between the effects of each treatment on gene expression variability and mean, respectively. While this negative correlation could be a consequence of correcting for the mean-variance dependency, our results suggest that for a subset of genes the treatment affects differently the mean and the variance, thus resulting in different response dynamics. Together, these findings can enhance our understanding of the molecular basis of how individuals respond to immune stimuli and suppress the activated immune system in response to drugs at the cell-type level. Genetic effects or environmental perturbations can affect the average gene expression, but disrupting the cis regulatory code can also reduce the tightness in the gene trascriptional response resulting in higher variability among individual cells. The mean gene expression response may not be affected but the dysregulation in gene expression variability may ultimately result in a modified drug-response phenotype.

Previous single cell studies in human induced pluripotent stem cells (iPSCs) differentiating into neurons and in response to oxidative stress identified a large fraction (46%) of eQTLs that had not previously been found in the Genotype-Tissue Expression (GTEx) catalog [42]. Similarly, we found that 44% of the eGenes identified in this study were not found in bulk leukocyte data from the same population which also includes the same individuals and had a larger sample size [94]. These additional discoveries were likely due to cell-type-specific genetic effects. However, when we consider genes expressed in all conditions (treatments and cell types) and use a rigorous statistical framework to determine to what extent genetic effects are shared across conditions we found a high degree of sharing as indicated by the fact that 65% of eGenes were significant in all 20 conditions. This is similar to the findings of the GTEx consortium that the majority of eQTLs are shared across all 49 surveyed tissues [1]. However, we found relatively fewer celltype-specific eQTLs compared to GTEx data across tissues [83], which indicates higher sharing of genetic effects across cell types within the same tissues than across different tissues. This finding is in line with the higher sharing of genetic effects among related brain tissues in GTEx data [83]. We previously explored the relative contribution of cell type and treatments in modulating genetic effects on gene expression using allele-specific expression. We found greater variability of genetic effects across three cell types than across 12 treatments [28]. Similarly, here we find more variation of genetic effects on gene expression across the 4 PBMC cell types when compared to the 5 treatment conditions. Importantly, we observed a similar pattern in gene expression variability and mean, although it was weaker for variability.

After filtering all eGenes with shared genetic effects across treatments, we discovered that 25% of eGenes have reQTLs. While eGenes were more likely to be differentially expressed and differentially variable in the majority of conditions, reGenes were not enriched for differentially expressed genes and only enriched for differentially variable genes in two conditions, PHA-activated monocytes and PHA+DEX treated T cells. This means that genes responsive to environmental conditions are also more likely to harbor genetic variants that affect gene expression levels, while genes with GxE effects are less likely to be detected as differentially expressed or differentially variable.

Identification of the genetic variants underlying inter-individual differences in cell-to-cell gene expression variability is technically more challenging than detecting genetic effects on mean gene expression. Previously, researchers have estimated that more than four thousand individuals would be needed to achieve 80% power to detect the strongest genetic effects on gene expression dispersion in induced pluripotent stem cells [100]. This calculation was based on a different technology, where each cell is processed as an independent library and the number of cells/individual that can be analyzed is very limited. However, with the technology available when we started our study, we were able to interrogate a large number of cells and individuals in the same library, thus reducing technical noise. This likely resulted in greater power to detect genetic effects on gene expression variability. Other factors that may contribute to our ability to discover these vQTLs are the cell types used and our study design. We observed that genetic effects on gene expression variability were most stable across different treatment conditions for monocytes, despite the strongest gene expression response to treatment in that cell type. We also demonstrated that treatments decrease the correlation of genetic effects on gene expression variability across the different cell types, demonstrating that GxE effects on gene expression variability were cell-type-specific. We found high enrichment for ribosomal protein genes within vGenes, which may be due to higher expression levels of this class of genes leading to an increased power to detect effects on gene expression variability.

Although we used a measure of gene expression variability that does not have a mean-variance dependency, we found genetic effects affecting both gene expression mean and variability but with opposite direction of effect for 13 genes. Even though eQTLs were enriched in vQTLs, we did not detect significant genetic effects on mean gene expression for most vQTLs. This demonstrates that genetic variants may increase variability of gene expression across cells, without affecting mean gene expression levels. While eGenes were enriched for differentially expressed genes, vGenes were depleted for them. This could be because genes responding to treatments are required to be under tighter genetic control of expression variability. Prior work on variance of gene expression and of its genetic control was performed on bulk RNA-seq datasets, therefore investigating variance in the population. These studies demonstrated that genes with a TATA box have increased noise both at the gene and allelic expression level (Mogno et al., 2010; Sigalova et al., 2020; Findley et al, 2021). TATA box promoters, as opposed to CpG island-associated promoters, have been associated with tissue-specific genes and high conservation across species (Carninci et al., 2006). Findley et al showed that genes which are more tolerant to loss of function mutations showed greater allelic expression residual variance, indicating that redundancy in gene function allows for less stringent genetic control. This could allow for the evolution of new regulatory elements, resulting in new patterns of gene expression. Conversely, genes which are under negative selection (i.e. low dN/dS) have low residual variance, underscoring the importance of preserving stable expression of these genes. Single cell studies allow analysis of variability within individuals, as a new molecular phenotype with potentially relevant biological function. Our results suggest that a tighter genetic control of gene expression variability is preserved in specific cell types and conditions. Specifically, vGenes were depleted of DVGs within all contrasts in monocytes but enriched for DVGs in the other cell types following immuno-stimulation and in T cells treated with dexamethasone. Additionally, we have demonstrated that effects of genetic variation on gene expression variability could vary across environmental conditions. Interestingly, these GxE effects on gene expression variability and the GxE effects on gene expression mean affected genes associated with the same immunological diseases.

Lastly, we introduced a novel computational approach designed to track the short-term transcriptional response dynamics to treatments at single-cell resolution. Our new method is supervised, easy to implement and can be directly applied to studying the dynamics of transcriptional responses to other treatments at single-cell level, to facilitate understanding the diversity in response to environmental stimuli across heterogeneous tissues. Previous studies investigated transcriptional dynamics in experimental designs where individual cells were sampled at a range of various times [13] or collected from various cell type differentiation stages [15]. In these studies, pseudotime was inferred from computational methods [8] or directly measured by experimental design [13]; but here we show that the pseudotime trajectory for these unsupervised methods is not very stable when resampling the data. The pseudotime inferred from our novel approach reflects the transcriptional dynamics of the immune response and is robust across resampling. Although our data were sampled at one time point, we still revealed transcriptional response dynamics using this projection, suggesting initial heterogeneity of cells, probably arising from diversity in initial cell states. Alternatively, this heterogeneity of response across cells may be a reflection of the dynamics of the immune system. We purposely used a study design where all the cell types were cultured together to study immune response, to preserve the natural interactions between cells that is key to orchestrate the immune response. This is reflected in the observed gene expression response to stimuli across all the cell types, even though we used immune activators which primarily affect one cell type. Thus, it is likely that in our study the observed heterogeneity of response across pseudotime reflects the dynamics of secondary signal propagation (cytokine or direct cell-to-cell interaction) across a population of cells. Using this method, we identified different dynamic patterns of gene expression along the response pseudotime and demonstrated widespread genetic regulation of these dynamics. Importantly, we demonstrated genetic effects on the dynamics of *ABO* expression response to glucocorticoids, which strongly correlated with susceptibility to SARS-CoV-2 infection [80, 103].

In summary, our study demonstrates that a full understanding of transcriptional response dynamics requires investigations beyond changes in mean gene expression, which are enabled by single-cell studies and robust computational methods. Elucidating gene expression variability and pseudotemporal response dynamics will contribute to a fuller picture of the heterogeneity within and across cell populations. Studies of transcriptional response dynamics and genetic contributions to their heterogeneity in future experiments of treatment response, tumor microenvironment dynamics and cell type differentiation across healthy and disease states will uncover new disease mechanisms based on altered control of gene expression heterogeneity.

## Methods

### Biological samples

Peripheral blood mononuclear cells (PBMCs) used in this study were collected from children aged between 10 and 16 years old as part of the longitudinal study Asthma in the Life of Families Today (ALOFT, recruited from November 2010 to July 2018, Wayne State University Institutional Review Board approval #0412110B3F) [93], which was established to explore the effects of family environments on childhood asthma. The 96 samples included in the current study were randomly selected from a pool of 136 samples passing the following criteria: donor’s age 10-16 years old, donor’s race self-reported as Black, availability of at least 3.5 million cryopreserved PBMCs, and to ensure a 50:50 ratio between female and male participants. For each participant multiple samples were collected longitudinally, but we included in this study only the earliest sample passing the filtering criteria. PBMCs collected into two BD Vacutainer™Glass Mononuclear Cell Preparation Tubes (Becton Dickinson and Co., East Rutherford, NJ) were extracted using a previously-published Ficoll centrifugation protocol [126], cryopreserved in freezing media (70% RPMI, 20% CS-FBS, 10% DMSO; 3 *−* 8 *×* 10^6^ cells/ml) and stored in liquid nitrogen until the day of the experiment.

### Cell culture and single-cell preparation

Cells were processed in batches of 16. For each batch, PBMCs were removed from liquid nitrogen storage and quickly thawed in 37*^◦^*C water bath before diluting with 6 ml of warm starvation media (90% RPMI 1640, 10% CS-FBS, 0.1% Gentamycin) and counting using Trypan blue staining on Countess II (Life Technologies Corporation, Bothell, WA). Cells were subsequently centrifuged at 400*×*g for 10 minutes and resuspended in culture medium at 2 *×* 10^6^ cells/ml. 500 *µ*l of cell suspension was plated in each of 5 wells of a 96-well round-bottom cell culture plate for each sample. Cells were incubated in starvation media overnight (approx. 16 hours) at 37*^◦^*C and 5% CO_2_. The following morning each of the five wells for each individuals were treated with either: 1 *µ*g/ml LPS [6] + 1 *µ*M dexamethasone [77], 1 *µ*g/ml LPS + vehicle control alone, 2.5 *µ*g/ml PHA [77] + 1 *µ*M dexamethasone, 2.5 *µ*g/ml PHA + vehicle control alone, or vehicle control alone (control). Note that the vehicle control used was 1*µ*l of ethanol in 10 ml of media, so that the effect of the vehicle control was negligible. After six hours, cells were pooled across individuals for a total of five treatment-specific pools. The pools were centrifuged at 300 rcf for 5 min at 4*^◦^*C, washed with 5 ml ice-cold PBS + 1% BSA and centrifuged again. Each pool was resuspended in 2 ml ice-cold PBS + 1% BSA and filtered through a 40 *µ*m Flow-Mi™strainer (SP Scienceware, Warminster, PA). Cell concentration was determined using Trypan blue staining on Countess II, and adjusted to 0.7*×*10^6^ cells/ml to 1.2 *×*10^6^ cells/ml. Each pool was loaded onto a separate channel of 10XGenomics®Chromium machine (10XGenomics, Pleasanton, CA), according to the manufacturer’s protocol, with batches 4, 5 and 6 loaded on two separate Chromium Chips for a total of 2 wells per treatment pool. Batches 1-4 and one chip of batch 5 were processed using v2 chemistry, and the other chip of batch 5 and batch 6 were processed using v3 chemistry. Library preparation was done according to the manufacturer’s protocol.

### Sequencing

Sequencing of the single-cell libraries was performed in the Luca/Pique-Regi lab using the Illumina NextSeq 500 and 75 cycles High Output Kit with 58 cycles for R2, 26 for R1, and 8 for I1.

### Genotype data

All individuals in this study were genotyped from low-coverage (*∼*0.4X) whole-genome sequencing and imputed to 37.5 M variants using the 1000 Genomes database by Gencove (New York, NY). These data were used for all genetic analyses and to calculate PCs of genotypes to be used as covariates in downstream statistical analyses. Genotype PCA was run on all biallelic autosomal SNPs with cohort MAF*<*= 0.1 using library SNPRelate [134] in R 4.0.

### single-cell RNA-seq raw data processing (Alignment and demultiplexing)

The raw FASTQ files were mapped to the GRCh38.p12 human reference genome using the kb tool (a wrapper of kallisto and bustools) with the argument of workflow setting to be lamanno [69]. Two procedures were performed in the kb tool. First, we used kallisto to pseudo align reads to the reference genome and quantify abundances of transcripts [10]. Second, we transformed the kallisto outputs into BUS (Barcode, UMI, Set) single cell format using bustools [70]. A total of 45 pooled libraries were generated in our experiment. Among 45 libraries and 6 batches, 3 experiments (LPS+DEX, PHA and PHA+DEX) in batch 2 and 3 experiments (LPS+DEX, PHA and PHA+DEX) in batch 3, were excluded because of failure of the 10x Genomics instrument resulting in broken emulsions, for a total of 39 libraries remaining for all subsequent analyses. We removed debris-contaminated droplets using the DIEM R package [3]. With this procedure, we obtained a count matrix of 301,637 cells with 116,734 genes features (including spliced and unspliced) across 39 library pools. The aligned counts matrix were transformed into a Seurat object for the subsequent functional analysis. To demultiplex the 16 individuals pooled together for each batch, we used the popscle pipeline (dsc-pileup followed by demuxlet) with the default parameters [44]. Two input files are required to provide for dsc-pileup, a BAM file and VCF file. The BAM files were generated by running cellranger (v2.1.1) count function aligned to the GRCh37 human reference genome (downloaded from 10X genomics, 3.0.0). The VCF file obtained from DNA genotype data (see above) was filtered to remove any SNP that was not covered by scRNA-seq reads. The resulting VCF contained 935,634 SNPs after removing SNPs with MAF less than 0.05. After assigning the identity to each cell, we removed mismatching barcodes between individual identity and batch. A total of 292,394 cells with 116,734 genes (including spliced and unspliced) were entered into the downstream analysis. For 96 Individuals, we had both control and LPS conditions, while the other three conditions (LPS+DEX, PHA, PHA+DEX) were assayed for 64 individuals because of instrument channel failure in batches 2 and 3 for those experimental conditions. This clean dataset had a median of 7,994 cells (Figure S48), a median of 4,251 UMI counts, and 1,810 genes measured on average in each cell across 39 experiments (Figure S49). In terms of spliced reads, we detected a median of 2,705 UMIs and 892 genes for each cell across all 39 experiments (Figure S50).

### Clustering, UMAP and cell type annotation

`Seurat`(V3) was employed for preprocessing, clustering, and visualizing the scRNA-seq data [110]. We performed log-normalization on all the data using `NormalizeData` with default parameters and selected 2,000 highly variable genes using `FindVariableFeatures` with variance-stabilizing transformation (vst) followed by standardization of these highly variable genes for downstream analysis using `ScaleData`. Linear dimensionality reduction was carried out by `RunPCA` on scaled data with 100 principal components (PCs). We ran `RunHarmony` with chemistry as covariates to correct chemistry effects (V2 and V3), which is scalable to a large number of cells and robust [50]. The Harmony-adjusted PCs were then used to construct a Shared Nearest Neighbor (SNN) graph using `FindNeighbors` (dims=1:50) and cell clustering was subsequently implemented by running `FindClusters` with the resolution set to 0.15. Finally, Uniform Manifold Approximation and Projection (UMAP) was applied to visualize the clustering results using the top 50 Harmony-adjusted PCs. Cells from different chemistries, batches, and treatments shared similar clustering patterns (Figure S51, Figure S52, Figure S53).

We identified 13 clusters then annotated cell clusters based on the following canonical immune cell type marker genes (Figure S2), B cells (*MS4A1* or *CD79A*), Monocytes (*CD14* or *MS4A7*), natural killer (NK) cells (*GNLY* or *NKG7*) and T cells (*CD3D* or *CD8A*). Clusters 0, 4, 5, and 7-12 were annotated as T cells with 183,289 cells, clusters 3 and 6 were annotated as monocytes (30,393), cluster 1 was defined as NK cells (47,824) and cluster 2 as B cells (30,888).

### Aggregation of single-cell-level data for differential gene expression and genetic analyses

We generated pseudo-bulk RNA-seq data by summing the spliced counts from `kallisto` for each gene and each sample across all cells belonging to each of the four cell types (B cell, Monocyte, NK cell and T cell), separately. This generated a data matrix of 42,554 genes from autosomes (rows) and 1,536 combinations of cell type+treatment+individual (columns). The number of cells in each combination displayed a similar distribution between treatments for most cell-types (Figure S54 and Table S2, S23). B cells have the smallest number of cells for each combination while T cells have the most number of cells for each combination (Figure S54), which is consistent with the cell-type compositions in PBMCs. Next, we focused on protein coding genes and filtered out genes with less than 20 reads across cells and removed combinations with less than 20 cells. This last filtering step resulted in a data matrix of 15,770 protein coding genes (rows) and 1,419 combinations (columns), which were used for all subsequent differential expression and genetic analyses.

### Differential gene expression analysis

We carried out differential gene expression analysis for each of the five batches separately using R DESeq2 package [57] and then combined the results across five batches using meta-analysis. For each batch, we estimated gene expression effects of the treatments using four contrasts: (1) LPS, LPS vs CTRL; (2) LPS+DEX, LPS+DEX vs LPS; (3) PHA, PHA vs CTRL; (4) PHA+DEX, PHA+DEX vs PHA. We found that compared to adding a batch covariate in DESeq2, our approach of dividing the analysis by batch produced more robust results in our data, likely because the overdispersion parameter of DESeq2 was estimated for each batch separately. In the meta-analysis procedure, we first extracted the summary statistics, including estimated effects (*β_i_*) and standard error (*s_i_*). Based on the fixed-effects model of meta-analysis, the weighted average effects are calculated by 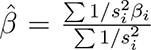 and its estimated variance can be expressed by 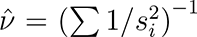. We then constructed the test statistics 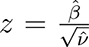 which follows a normal distribution under the null hypothesis. To correct for multiple hypothesis testing, we used the qvalue function in R 4.0 to estimate false discovery rate (FDR) from a list of *p*-values from the above *z* test statistics for each condition, separately. Differentially expressed genes (DEG) were defined as those with FDR less than 0.1 and absolute estimated log_2_ fold change larger than 0.5. We carried out gene ontology (GO) enrichment analysis [i.e., Biological process (BP), molecular function (MF), and cellular component (CC)], for these DEGs identified across conditions using `R ClusterProfiler` package (V3.16.1) [132].

### Estimation of gene expression mean and dispersion

Using the single-cell data generated from Fluidigm experiment Sarkar et.al proposed a zero-inflated negative binomial (ZINB) distribution to estimate gene expression variance [100]. Some literature demonstrated that it is not necessary to consider the zero inflation term in the model for droplet-based single-cell RNA data [111]. Based on the computational framework that was established by Sarkar et.al, we adapted a negative binomial (NB) distribution to model the count data for each gene *j*, using two main parameters for the mean(*µ_j_*) and dispersion(*ϕ_j_*) in our scRNA data. Then, we derived the variance of gene expression by 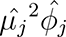. The details on NB model are described as follows,

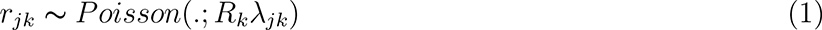

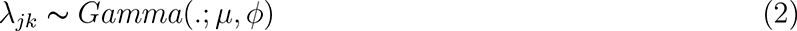

Where *r_jk_* is the number of molecules for *k* cell, *j* gene;*R_k_* is a size factor of each cell, equal to total reads in each cell divided by median reads across cells, to account for difference in cellular sequencing depth; *λ_jk_* is a latent variable, representing gene abundance and further is assumed to be Gamma distribution with mean value *µ*, variance *µ*^2^*ϕ*. By integrating the latent variable *λ_jk_*, we derived density function for each observation as follows,

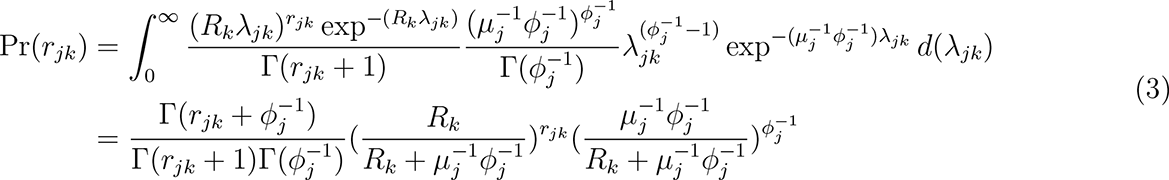

Then, we estimated parameters of *µ_j_* and *ϕ_j_* by minimizing the negative log likelihood using the optim function in R 4.0 using the “L-BFGS-B” algorithm. Two filters were applied prior to estimation of gene expression parameters (mean and dispersion): (1) removing genes with less than 20 reads across cells (with a total of 30,972 autosomal genes remaining for the analysis); (2) removing the combinations (cell type+treatment+individual) with less than 20 cells (with a total of 1,419 combinations across all our data). To guarantee robust parameter estimation, we focused on genes with more than 15 reads that are expressed across at least 15 cells for each cell type/treatment combination.

Similar to what reported in previous studies [20, 22, 24, 27], we observed that both gene variance or dispersion linearly depended on mean parameters (Figure S11A,B). To further correct the dependence between dispersion and mean, we defined a residual dispersion, capturing the departure from the global trend. The residual dispersion was calculated by removing the part of dispersion that could be predicted by the overall trend of gene mean across genes in each cell type separately. This adjusted dispersion was uncorrelated with the gene mean value (Figure S11C) across genes.

The well established method (regression BASiCs model) were proposed to perform differential test for gene expression variability, which can correct the mean confounder by fitting a global linear trend of mean and dispersion across genes in a bayesian hierarchical framework [21]. In analogy to the BASiCs model, we quantified gene expression variability that was not confounded by mean expression by the two discrete stages (1) Use a negative binomial model to obtain the estimation of the parameters mean and dispersion similar to Sakar et al. approach; (2) Fit a linear model between mean and dispersion across genes to remove the part of dispersion that can be predicted by mean expression. We always used the adjusted values unless otherwise stated.

### Calculation of pathway specific score

We calculated the score of a specific GO term or pathway using the transcriptional matrices (mean and dispersion) patterns of genes involved in that term/pathway across all the conditions as follows: (1) extracted the subset of differentially expressed genes in at least one condition belonging to a specific GO term or pathway; (2) scaled mean-centered gene matrices values (mean and dispersion) for each gene across individuals for each cell type and batch separately; (3) subtracted gene expression values of individuals in the CTRL condition to calculate relative expression changes; and, (4) calculated the average values as a specific pathway score across genes for each individual.

### Differential gene variability analysis

We focused on 15,770 protein-coding genes for differential gene variability analysis. To account for the noise from the batch, we implemented the same strategy for differential gene expression analysis for differential gene variability and differential gene mean analyses via fitting models by batch separately and we meta-analyzed the summary statistics. We first took the log 2 transformation of residual dispersion and mean, then fitted models using linear regression where treatments were considered for each batch, separately. To obtain more robust estimation of the gene expression parameters for statistical inference, we only focused on genes that had at least 3 individuals in the treatment and 3 individuals in the control condition that was being contrasted. The differentially variable genes (DVG) and differential mean genes (DEG) were defined as those with FDR less than 0.1 and absolute value of log 2 fold change greater than 0.5. We also used ClusterProfiler in R to carry out GO enrichment analysis for these identified DVGs.

### DLDA response pseudotime method

We developed a new pseudotime method based on diagonal linear discriminant analysis (DLDA) to characterize the degree to which single cells respond to immune treatments. A key difference compared to previous methods [32, 88] is that DLDA is supervised and estimates a straight line trajectory between two endpoints which should be particularly suitable for a short time treatment. Basically, the trajectory is a one dimensional line connecting the two centroids of a treatment and control conditions for each cell-type. In DLDA only a subset of genes that are highly differentially express between the two conditions are used based on the following formula:

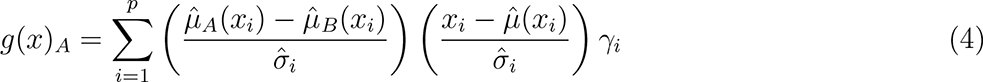

Where the subscripts of A and B represent the two conditions considered in each contrast, respectively; *µ̂_i_* denoting mean gene expression of the *i_th_* gene for A, B or across groups and *σ̂_i_* meaning standard deviation of the *i_th_* gene across conditions. In this application of DLDA we only used the DEGs that were differentially expressed in the corresponding condition (10%FDR by DESeq2) by introducing an indicator variable (*γ_i_*, assigning to 1 if the *i_th_* gene is a significant DEG in the contrast of A versus B or 0 otherwise. This is equivalent to denoising by hard thresholding instead of the soft thresholding that a method like nearest shrunken centroids would use. Note that the left term of the summation is a weight that is exactly a standardized effect size, and the right term evaluates for a given gene how far it is from the midpoint between all the cells. By ignoring the off-diagonal terms and only considering genes that are differentially expressed, DLDA achieves a more robust classification performance when the number of samples available for training is relatively limited. While this approach assumes the shortest Euclidean distance alignment between the centroids of the treated and untreated cells, it should be a reasonable assumption for short time-periods (here 6 hours), compared to learning a complex curved one-dimensional manifold in a high dimensional space. This assumption provides a more robust and stable trajectory followed by the cells from the control to the treated state when we iteratively repeat the procedure resampling the data, compared to more complex nonlinear trajectory methods.

Four kinds of DLDA axes were calculated by the following contrasts for each cell type separately, (1) DLDA axis of LPS was estimated in the CTRL and LPS conditions to infer the response pseudotime to LPS; (2) DLDA axis of LPS+DEX was computed in the LPS and LPS+DEX conditions to calibrate the response pseudotime to DEX; (3) DLDA axis of PHA was estimated in the CTRL and PHA conditions to characterize the response pseudotime to PHA stimuli; and, (4) DLDA axis of PHA+DEX was estimated in the PHA and PHA+DEX conditions to characterize the response pseudotime to DEX. To achieve this, we excluded the data from batches 2 and 3, resulting in 264,545 cells. Next, to correct chemistry effects, we performed standard log-normalization for each subset (split by chemistry) followed by integrating datasets together using seurat (V3) for each cell type, separately. After the DLDA representing the trajectory pseudotime is calculated for each cell-type and treatment, we use a sliding window approach (10% cells in each window, sliding along the defined pseudotime with a step of 0.1% cells), we analyzed gene expression dynamic changes along the response pseudotime within each treatment for each cell type separately and identified different dynamic patterns of gene expression using the k-means algorithm

### Validation of the accuracy of DLDA in response pseudotime

We performed resampling analysis to demonstrate the robustness and stability of the DLDA compared to two existing methods: Monocle 3 [88] and scanpy [127]. For the simplification of the analysis, we focused on the T-cells treated with PHA+DEX or PHA that were measured using 10x Genomics chemistry V3, including 29,157 cells and 6,571 DEGs. To validate the effectiveness of the novel approach (DLDA), we randomly re-sampled half the dataset for 50 replications. In each replication, we applied our novel approach DLDA and the two other methods to compute response pseudotime. We do not have the underlying ground truth on how the cells should be exactly ordered according to response pseudotime; yet, we can still assess how expression for each of the responding genes correlates with the estimated pseudotime in each iteration across all cells. Ideally, the cells would be sorted similarly in each iteration, and the correlation of the gene expression with the pseudotime would stay constant. For each iteration, we calculated the Pearson correlation of the pseudotime and each gene expression in cells. Two indexes were employed to assess the performance of the three methods, (1) variance of correlation coefficients for each gene across replications; (2). Jaccard index of the top 20 highly correlated genes between 1,225 pairwise replications.

### Data normalization for genetic analyses

For downstream genetic analyses, we first removed all lowly-expressed genes, defined as having less than 0.1 CPM in more than 20% of the samples in each condition, separately. We quantile-normalized the count data using the voom function in the `limma` v3.44.3 package [95] in R 4.0 and regressed out the following confounding factors: experimental batch, sex, age, and top three principal components of genotypes in each cell type-treatment combination, separately.

For gene expression variability, for each gene-treatment-cell type combination, we discarded data from batches represented by less than 3 individuals. We considered genes with gene expression variability measures available for at least 20% of remaining individuals in each treatment-cell type combination. We quantile-normalized the data using the voom function in the `limma` v3.44.3 package [95] in R 4.0. We regressed out the following confounding factors on the subset of genes with no missing data across all individuals in each condition: experimental batch, sex, age, and top three principal components of genotypes processing each cell type-condition combination and performed PCA on variability residuals to use as covariates in variability-eQTL mapping. Mean gene expression estimates were processed using the same pipeline as gene expression variability data.

### *cis*-eQTL mapping

We used `FastQTL` v2.076 [83] to perform eQTL mapping on gene expression residuals calculated for each cell type and condition as outlined above. For each gene, we tested all genetic variants within 50 kb of the transcription start site (TSS) and with cohort minor allele frequency (MAF)*>* 0.1. We optimized the number of gene expression PCs in the model to maximize the number of eGenes across all conditions combined. The model that yielded the largest number of eGenes included 4 gene expression PCs. eQTL discovery for mean and variability data was performed similarly, except we modelled quantile-normalized data and regressed out the effects of experimental batch, sex, age, and top three principal components of genotypes and PCs of residuals in the `FastQTL` model. The model that yielded the largest number of significant genes at 10% FDR included 5 PCs of residuals for variability and 7 PCs of residuals for mean data.

### Multivariate adaptive shrinkage

To improve power of eQTL discovery by taking advantage of parallel measures of genetic effects across many cell types and conditions, we employed the multivariate adaptive shrinkage (mash) method using the `mashr` package v.0.2.40 in R v4.0 [118]. As input, we provided the genetic effect size estimates from `FastQTL` analysis in each cell type-treatment combination, and their corresponding standard errors calculated as absolute effect size estimates divided by its *z*-score. We kept the gene-variant pairs with estimates across all cell types and conditions and eliminated the two genes causing a singular matrix. We fitted mash across all 20 cell type-condition combinations on a random subset of 200,000 gene-SNP pairs providing both canonical and data-driven covariance matrices (the latter estimated using mashr in-built cov ed function on strong eQTLs from full data set, defined as those with ashr local false sign rate (LFSR)*<* 0.05).The LFSR refers to the probability that we incorrectly inferred the sign of genetic effects size (*β_j_*), *LFSR_j_* = min {Pr (*β_j_ ≥* 0*|D*), Pr (*β_j_ ≤* 0*|D*)} [108]. We considered gene-variant pairs with posterior LFSR*<* 0.1 to be eQTLs. To analyze sharing of genetic effects across all conditions, we considered the direction of genetic effect across all conditions. Genetic effects are shared across conditions if they have the same sign and are significant in at least one of the conditions considered. To analyze treatment-or cell type-specificity, we focused on pairwise comparisons of genetic effect sizes. An eQTL is specific to a condition if it is significant at LFSR*<* 0.1 in either condition and either the direction of effect differs or the difference in the magnitude of the genetic effects is at least two-fold. A gene is considered shared/specific if at least one eQTL for that gene is shared/specific across the given set of conditions. To discover response eQTLs (reQTLs) we considered the pair-wise union of significant eQTLs in each of the 16 treatment-control combinations. We defined reQTLs as genetic variants whose effect size on gene expression differed by at least two-fold between treatment and control conditions. Mash analyses on mean and variability estimates were conducted following this pipeline.

### DLDA-eQTL mapping

For mapping dynamic eQTLs that have genetic effects on gene expression interacting with pseudotime, we first equally divided cells in one immune treatment into three tertiles of the corresponding DLDA pseudotime. The three tertiles denoted 1, 2 and 3, represent early, middle and latter response pseudotime, respectively. Then, we mapped eQTLs interacting with these representative tertiles as dynamic eQTLs across cell types in the corresponding immune treatments as follows:(1) eQTLs interacting with LPS pseudotime were identified in cells treated with LPS; (2) eQTLs interacting with LPS+DEX pseudotime were identified in cells treated with LPS+DEX; (3) eQTLs interacting with PHA pseudotime were identified in cells treated with PHA; and (4) eQTLs interacting with PHA+DEX pseudotime were identified in cells treated with PHA+DEX. To do this, we summed gene expression data in each of the three DLDA bins for individuals with at least two bins containing at least 5 cells for each treatment. We considered genes with *>*0.1 CPM for at least 20% of remaining individuals in each treatment-cell type combination. We quantile-normalized the data followed by regressing out the following confounding factors: experimental batch, sex, age, and top three principal components of genotypes processing each cell type-condition combination separately. To identify genetic variants interacting with each DLDA, we fitted a linear model using the lm function in R 4.0 that included both the genotype dosage and the DLDA bin (numerically encoded as 1-3), as well as their interaction: Expression *∼* dosage + DLDA bin + dosage *×* DLDA_bin in each condition separately. Using the same cis-regions to the above analysis, we perform the interaction analysis for all genetic variants within 50 kb of the transcription start site (TSS) in the genes. We then applied Storey’s q-value method on the p-values for the interaction term using a stratified FDR approach on significant and non-significant eGenes (from the `FastQTL` analysis, Figure S45).

### Integration with transcriptome-wide association studies results

We considered the results of probabilistic TWAS [133], where eQTL from whole blood tissue in GTEx v8 were integrated with the GWAS summary data of asthma. PTWAS identified 4, 16, 66 and 72 (union of 78 genes)genes associated with asthma for four GWAS studies: GABRIEL [74], TAGC [16]) and two from UK Biobank [86] (self reported asthma and asthma diagnosed by doctor asthma). To combine multiple TWAS Z-scores across studies for the same gene, we selected the one with the largest magnitude as representative. We then identified genes that were both associated with asthma risk and differentially expressed in any of the treatment conditions.

## Data and code availability

All the data produced in this manuscript is being submitted to dbGAP (accession number phs002182.v2.p1). Code and scripts used to make the results are available at https://github.com/piquelab/scaip and https://github.com/piquelab/SCAIP-genetic.

## Acknowledgments

This research has been supported by NIH grants from NHLBI (R01HL114097 to SZ and RS, 1R01HL162574 to FL, RP, and SZ) and NIGMS (R01GM109215 to FL and RP). We thank members of the Luca, Pique-Regi, Slatcher and Zilioli labs and Xiaoquan Wen for helpful discussions and comments. We also thank the anonymous reviewers for comments and suggestions that helped to improve the quality of this manuscript. Finally, we also greatly appreciate the ALOFT study participants and their families.

## Supplementary Figures

**Figure S1:**
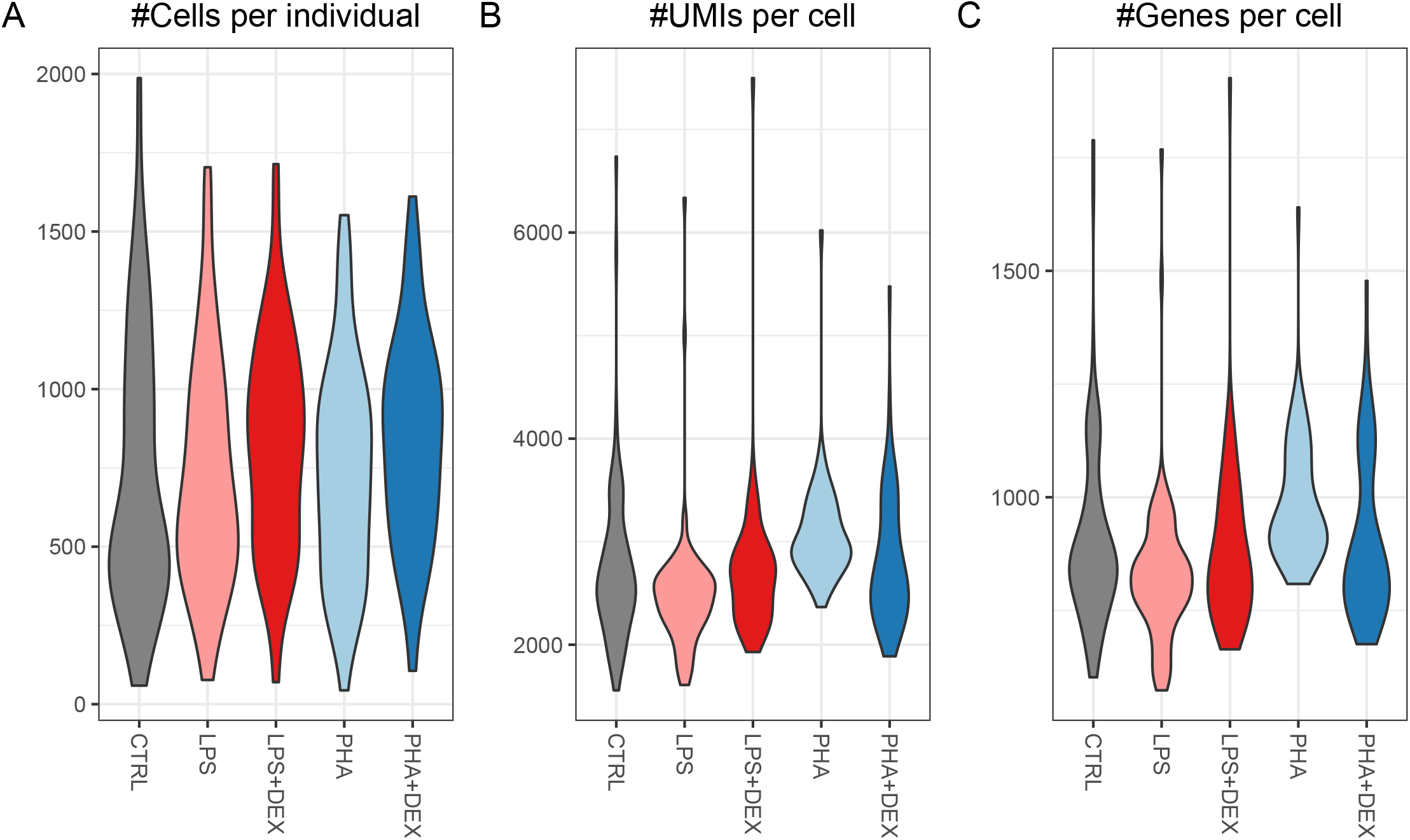
Violin plots of summary statistics per individual across five conditions. **(A)** number of measured cells per individual for each treatment. **(B)** average UMIs per cell from one individual for each treatment. **(C)** average number of detected genes per cell from one individual for each treatment.

**Figure S2:**
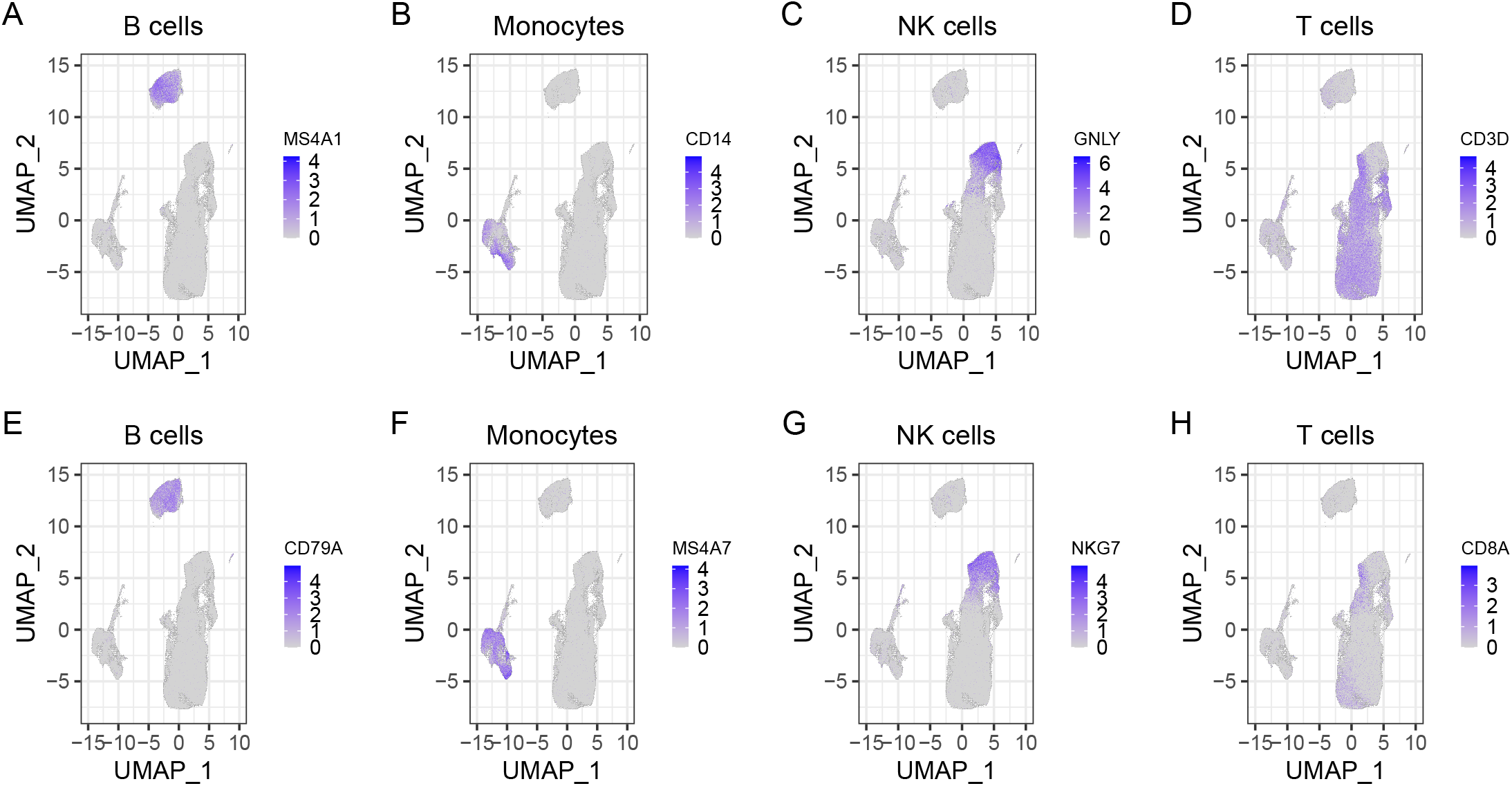
Canonical immune cell type-specific markers expressed on UMAP: **(A and E)** represent B cell-specific gene expressions (*MS4A1* and *CD79A*) on UMAP; **(B and F)** represent Monocyte-specific gene expressions (*MS4A7* and *CD14*) on UMAP; **(C and G)** represent NK cell-specific gene expressions (*GNLY* and *NKG7*) on UMAP; **(D and H)** represent T cell specific gene expressions (*CD3D* and *CD8A*) on UMAP.

**Figure S3:**
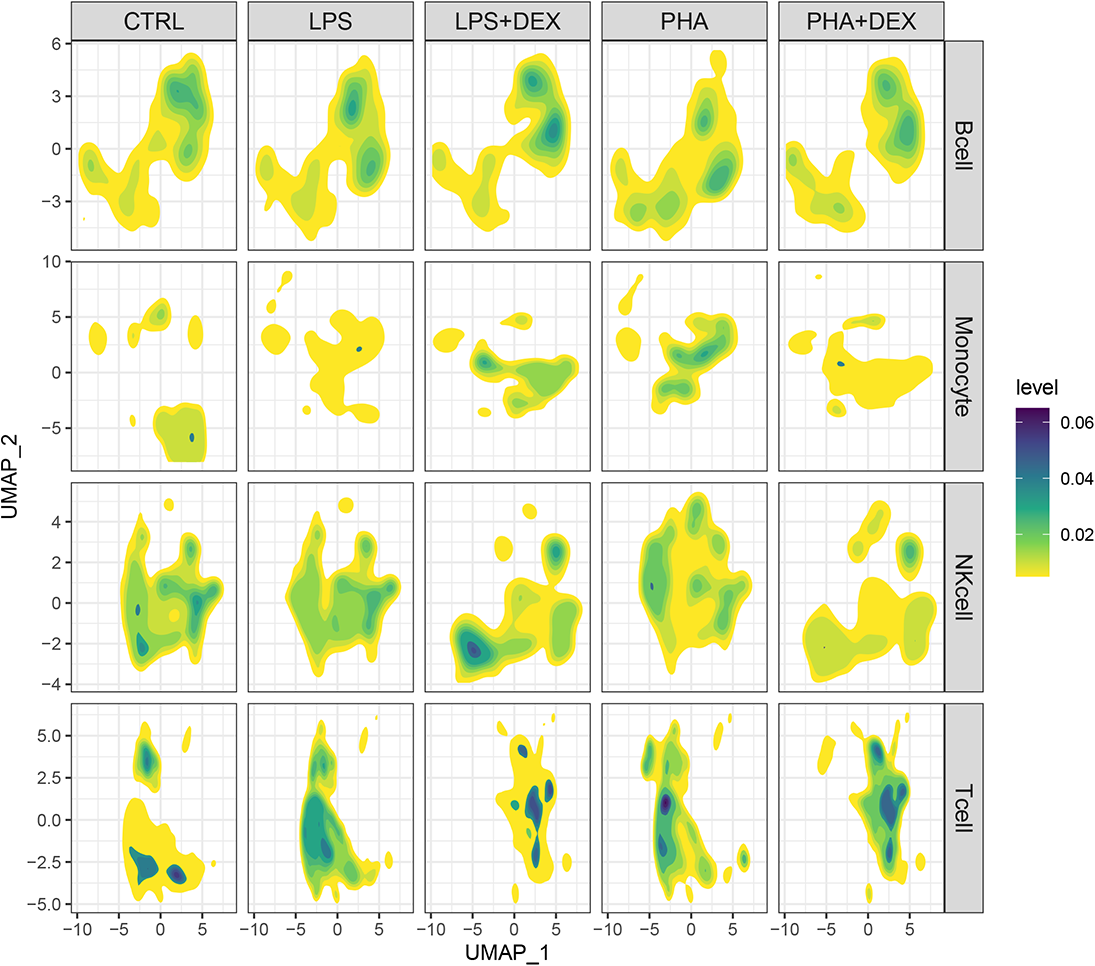
Density distribution of cells on UMAP for cell types and treatment separately. Blue color indicates high density of cells while yellow color indicates lower density.

**Figure S4:**
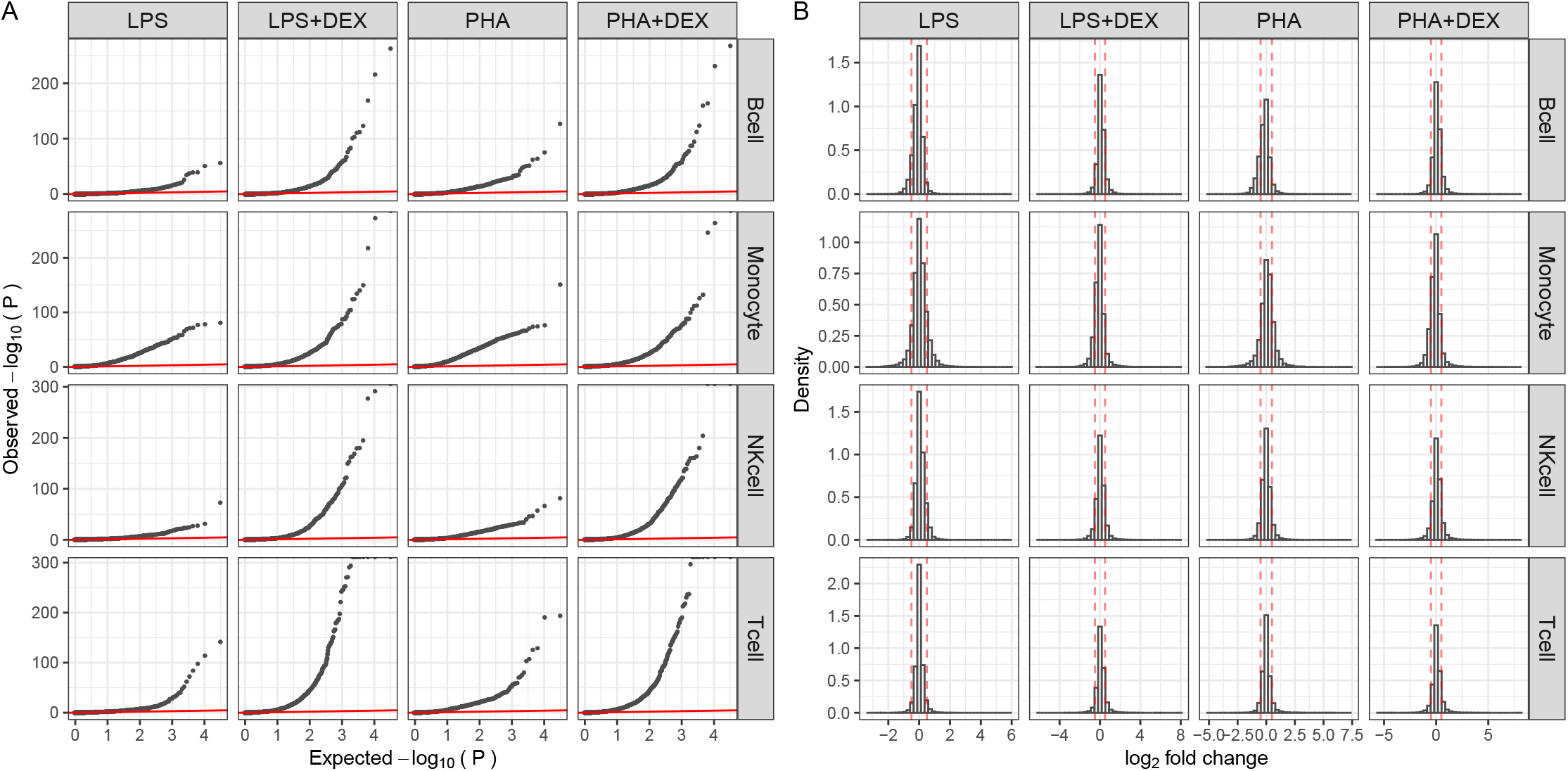
Overview of results of differential expression analysis: **(A)** Q-Q plots for 4 contrasts (LPS, LPS+DEX, PHA and PHA+DEX) across cell types, **(B)** Histograms of log_2_ fold changes of gene expression, red dashed lines denoting the —log2FoldChange— being 0.5, the cutoff value that is used for defining differentially expressed genes (DEGs).

**Figure S5:**
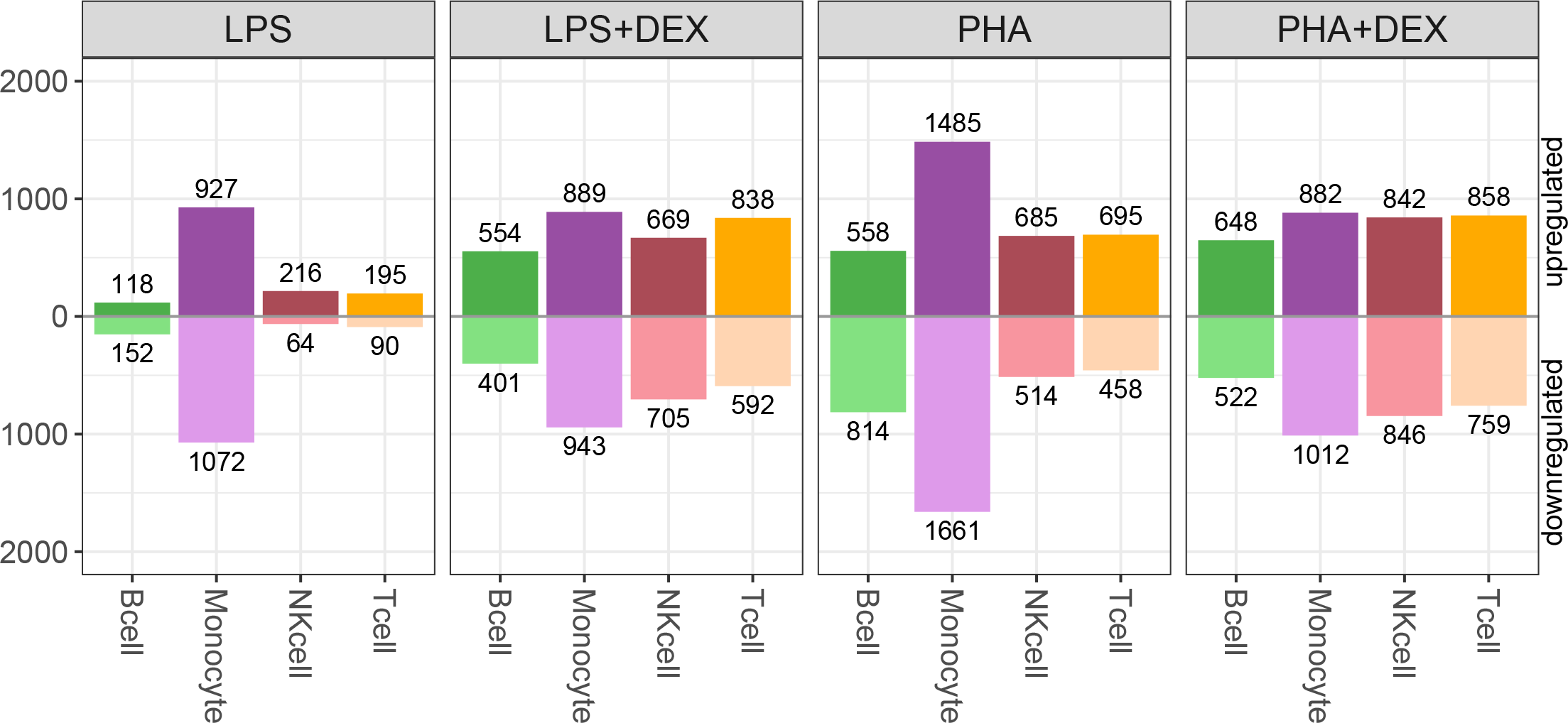
Number of DEGs across contrasts and cell-types; B cells (green), Monocytes (purple), NK cells (maroon) and T cells (orange), below axis representing down-regulated genes and above axis representing up-regulated genes (FDR*<*10% and —LFC—*>*0.5%).

**Figure S6:**
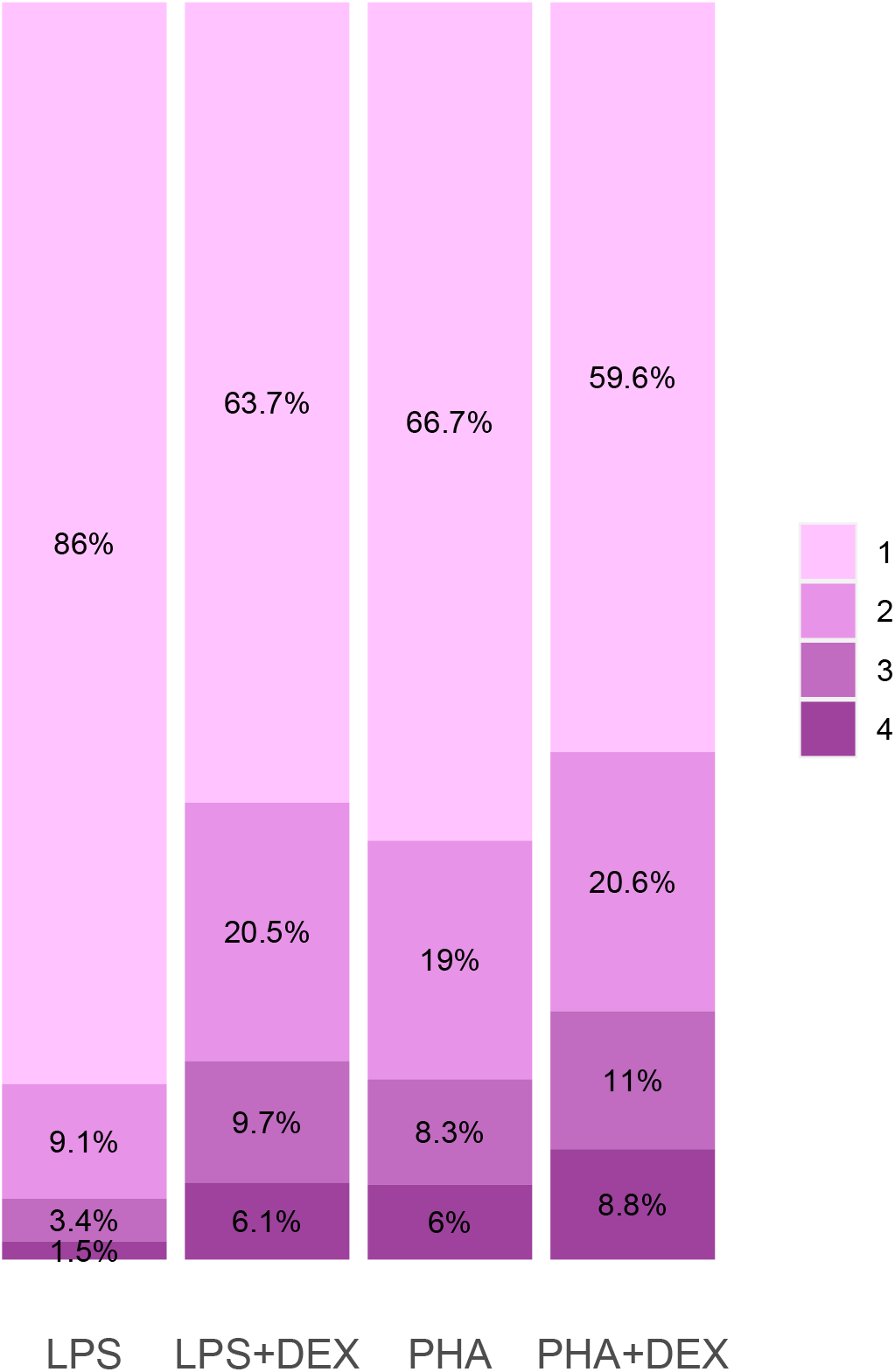
Cell type sharing of DEGs in four contrast conditions, 1=DEGs detected in only a single cell type and 4=DEGs detected across all the four cell types.

**Figure S7:**
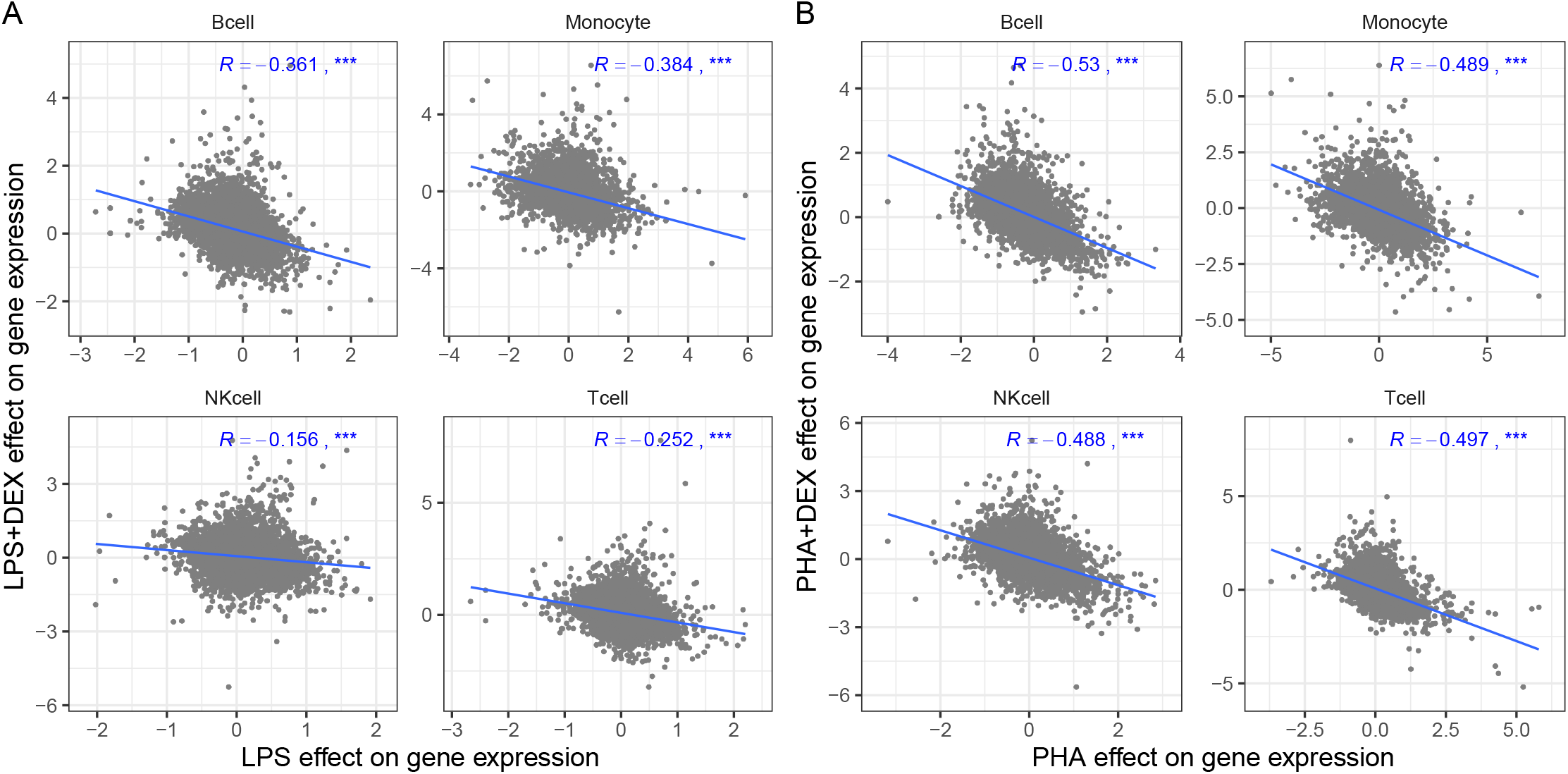
Dexamethasone (DEX) reverses effects of activation by immune stimuli on gene expression:. **(A)** Scatterplots of LPS effect on gene expression (x axis) against DEX effect on gene expression (y axis). **(B)** Scatterplots of PHA effect on gene expression (x axis) against DEX effect on gene expression (y axis). Blue lines represent the linear trend between the effects of immune stimuli and DEX treatments on gene expression. *R* represents the correlation coefficients of the effects of two types of immune treatments on gene expression, *denoting *p <* 0.05, ** denoting *p <* 0.01 and *** denoting *p <* 0.001.

**Figure S8:**
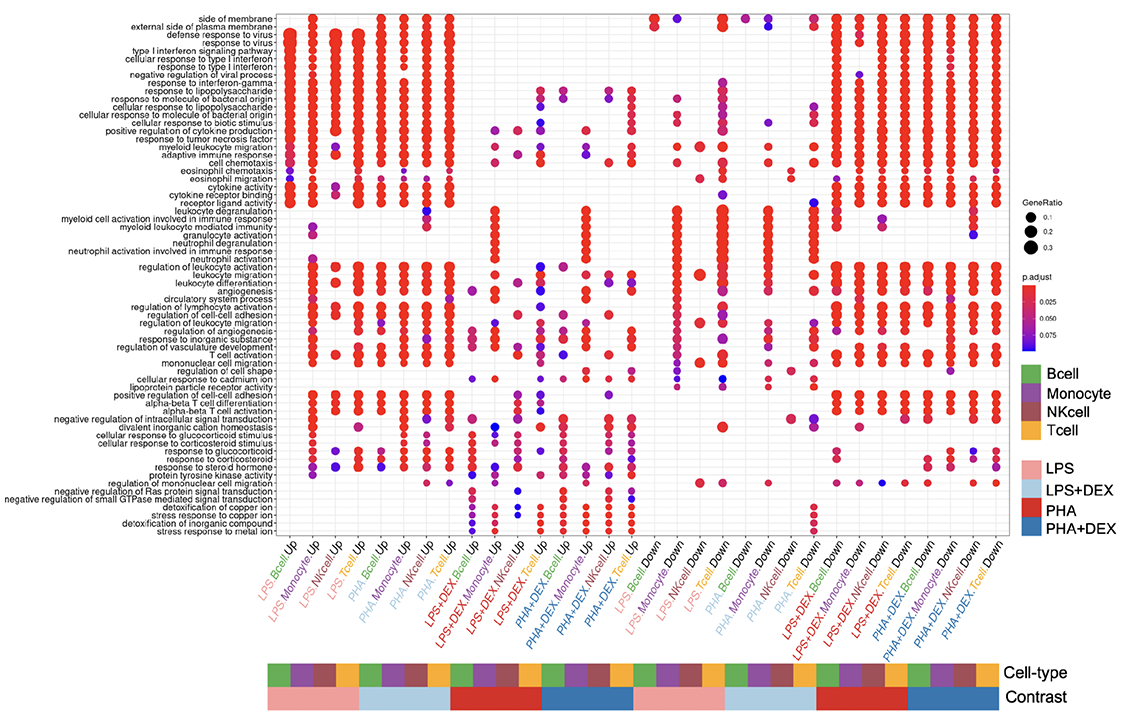
Visualization of Gene ontology (GO) enrichment analysis: The dotplots show the union of the top 5 GO terms for each condition (down-regulated and up-regulated separately) across 16 conditions (4 contrasts *×* 4 cell-types).

**Figure S9:**
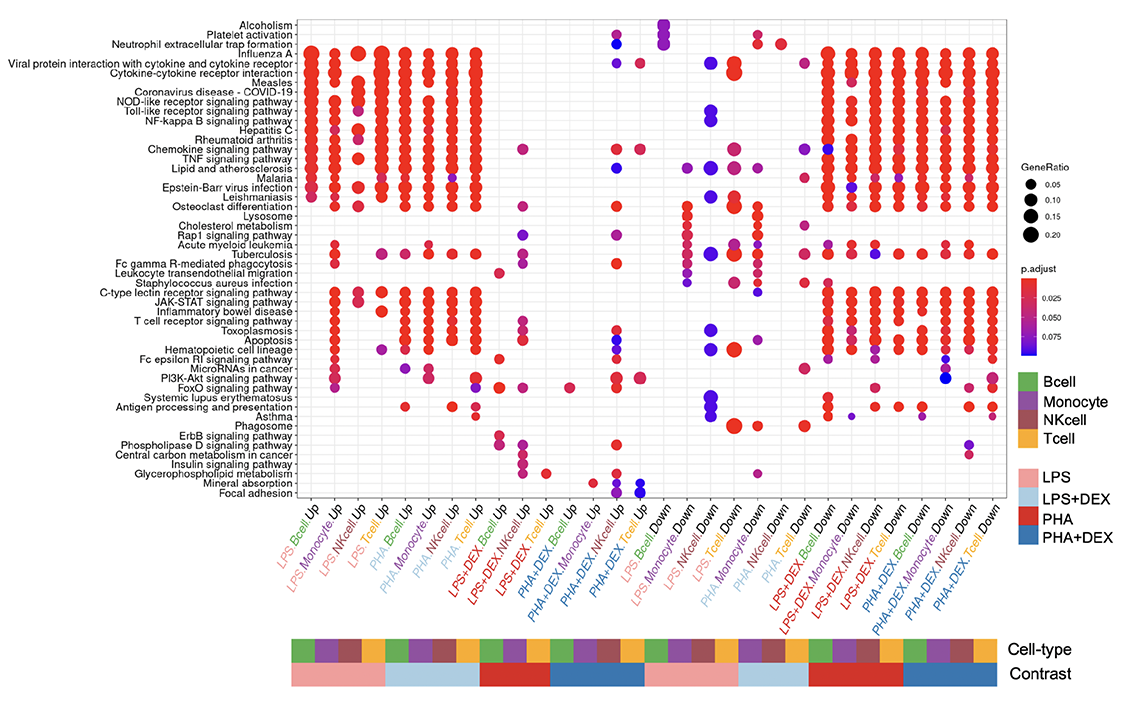
Visualization of Kyoto Encyclopedia of Genes and Genomes (KEGG) enrichment analysis: The dotplots show the union of the top 5 KEGG terms for each condition(down-regulated and up-regulated separately) across 16 conditions (4 contrasts *×* 4 cell-types).

**Figure S10:**
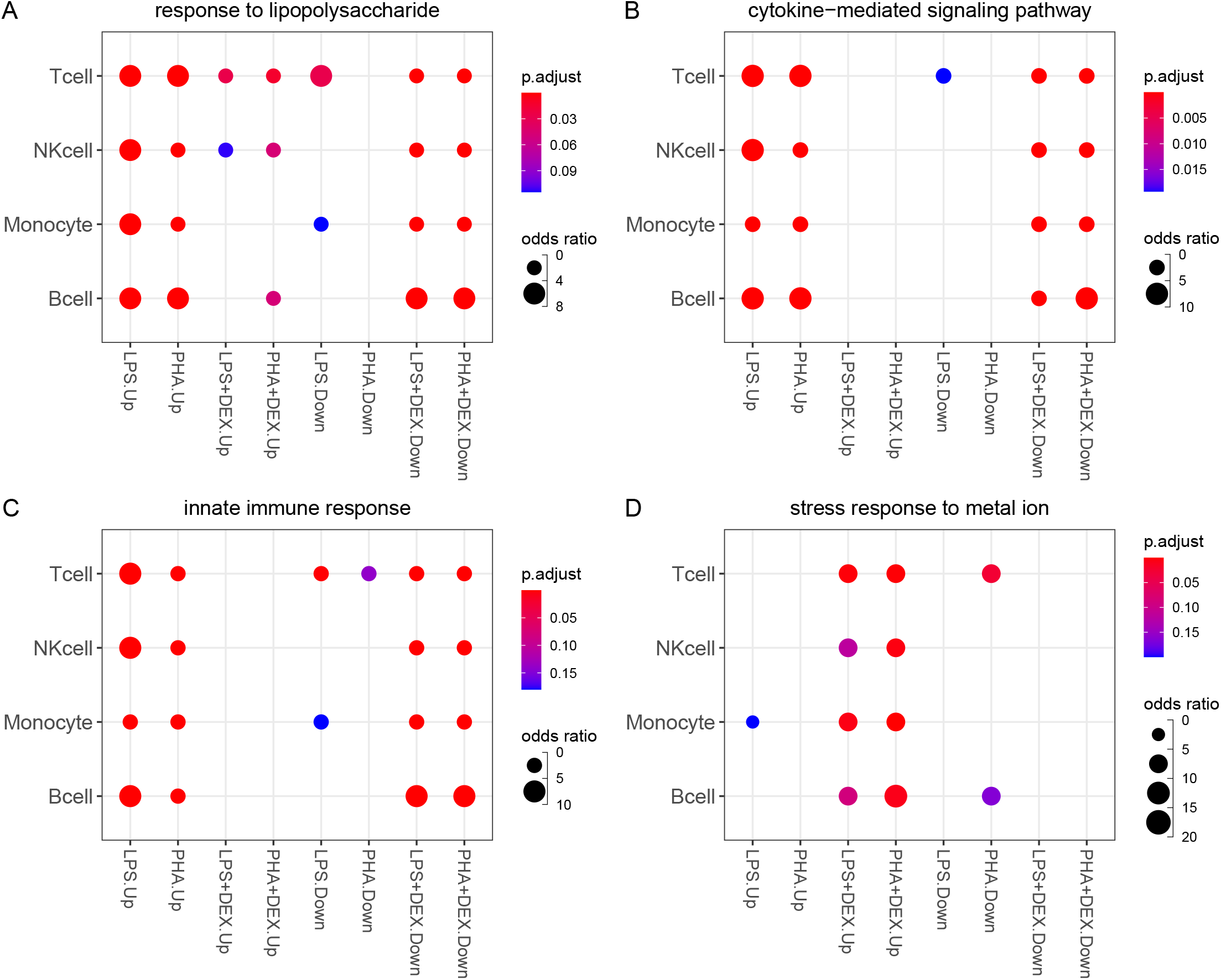
Examples of the enriched pathways for DEGs: **(A-D)** represents the enrichment results of the four example pathways across cell-types and conditions,, response to lipopolysaccharide, cytokine-mediated signaling pathway, innate immune response and cellular response to glucocorticoid stimulus, respectively.

**Figure S11:**
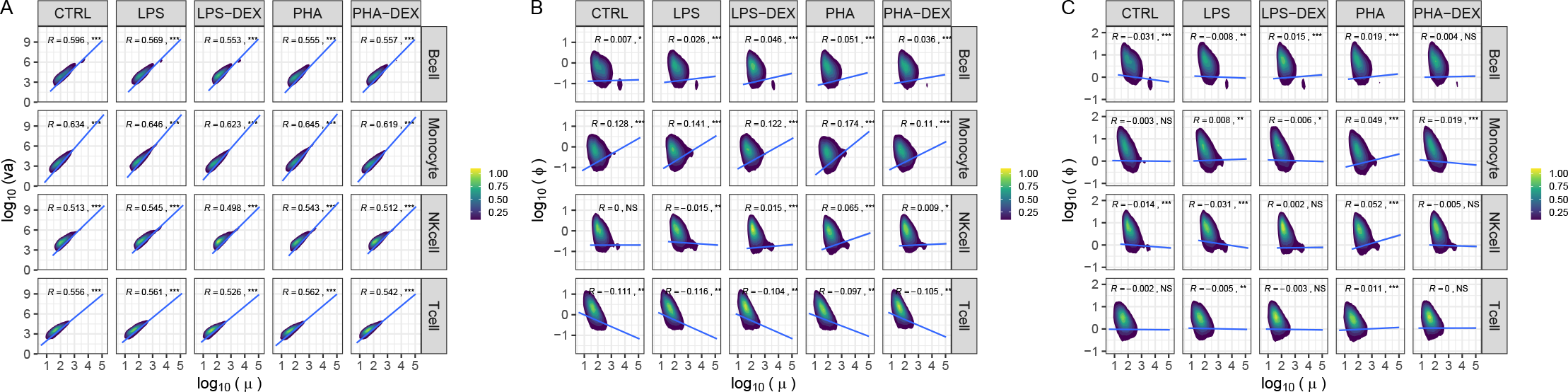
Both variance and dispersion of gene expression strongly depend on mean gene expression while mean-corrected dispersion is relatively independent on mean expression value: **(A)** Scatter plots of log_10_ mean gene expression (x axis) against log_10_ variance of gene expression. **(B)** Scatter plots of log_10_ mean gene expression (x axis) against log_10_ dispersion of gene expression. **(C)** Scatter plots of log_10_ mean gene expression (x axis) against log_10_ mean-corrected dispersion of gene expression, eventually used for measuring gene expression variability.

**Figure S12:**
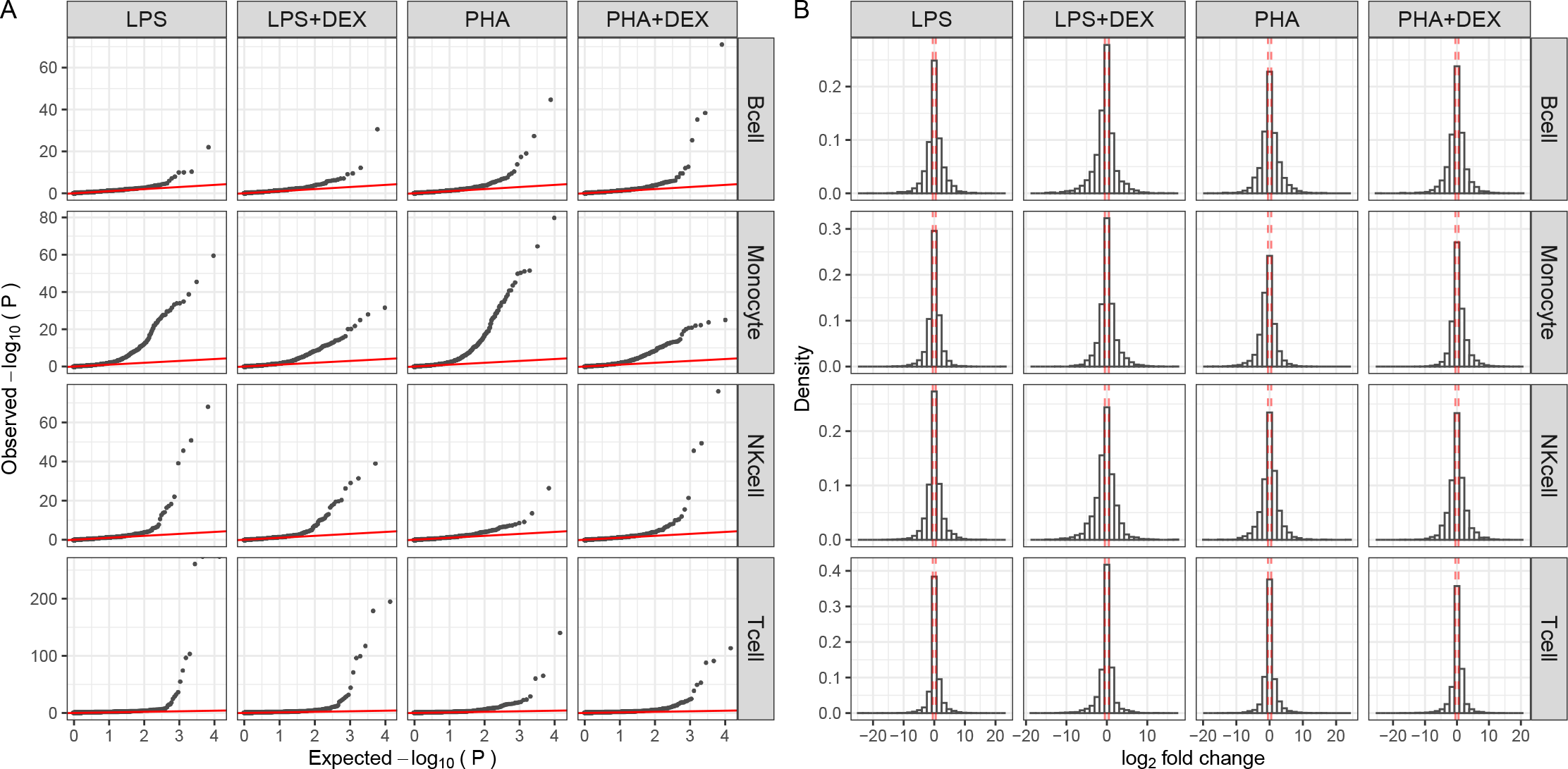
Overview of results of differential gene variability analysis: **(A)** Q-Q plots for 4 contrasts (LPS, LPS+DEX, PHA and PHA+DEX) across 4 cell types. **(B)** Histograms of log_2_ fold changes of gene variability, red dashed lines denoting the —log2FoldChange— being 0.5, the cutoff value that is used for defining differentially variable genes (DVGs)

**Figure S13:**
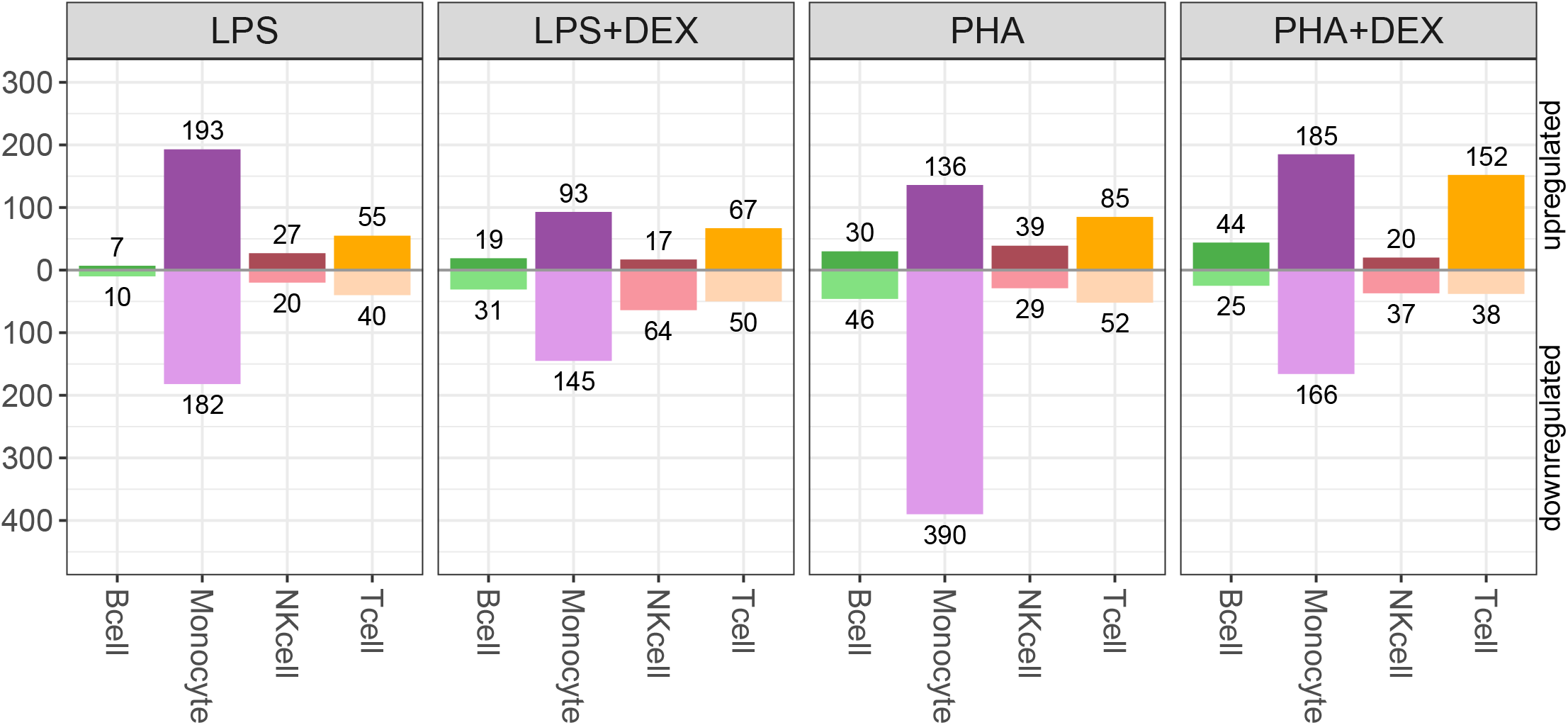
Number of DVGs across contrasts and cell types: B cells (green), Monocytes (purple), NK cells (maroon) and T cells (orange), below axis representing genes with decreased variability and above axis representing genes with increased variability (FDR*<*10% and —LFC—*>*0.5%)

**Figure S14:**
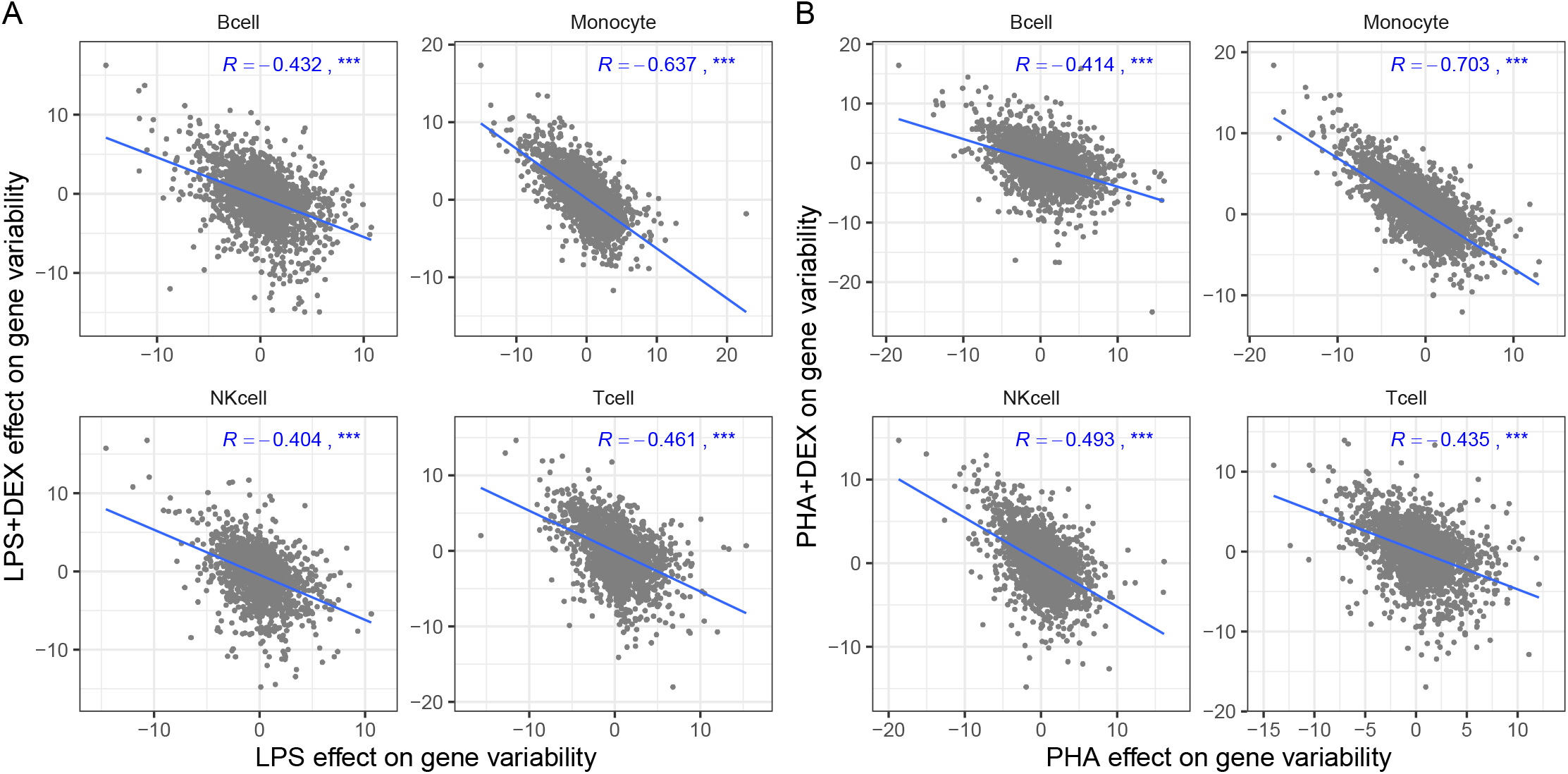
Dexamethasone (DEX) reverses effect of activation by immune stimuli on gene variability. **A** Scatterplots of LPS effect on gene variability (x axis) against DEX effect on gene variability (y axis). **B** Scatterplots of PHA effect on gene variability (x axis) against DEX effect on gene variability (y axis). Blue lines represent the linear trend between the effects of immune stimuli and DEX treatments on gene expression. *R* represents the correlation coefficients of the effects of two types of immune treatments on gene expression, *denoting *p ≤* 0.05, ** denoting *p ≤* 0.01 and *** denoting *p ≤* 0.001.

**Figure S15:**
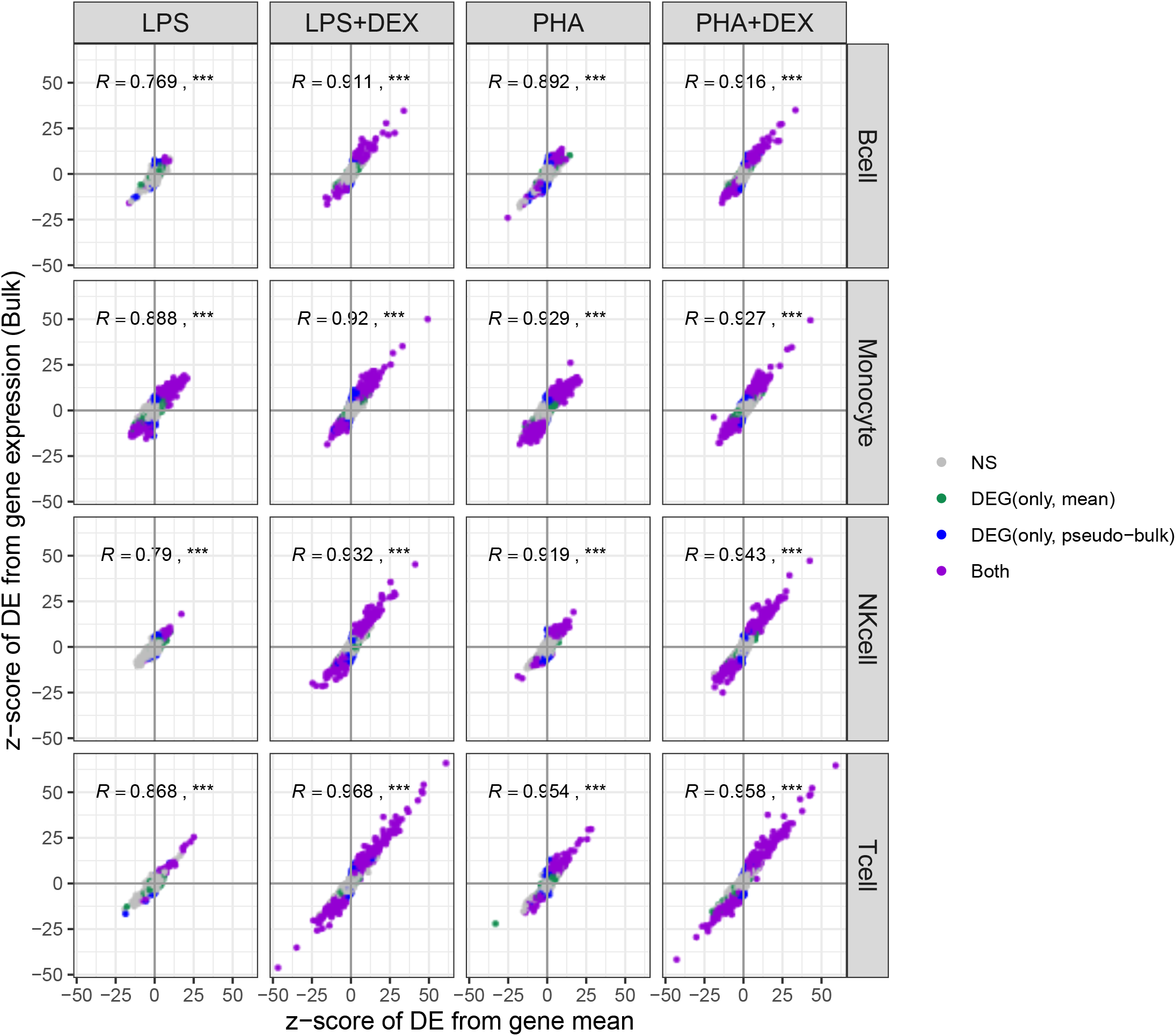
The scatter plots of the z score of treatment effects on gene mean (x axis) from Negative bionomial distribution against the z score of treatment effects on gene expression (y axis) from bulk data: Color represents significant treatment effects on gene mean only (green), bulk-derived DEGs only (blue), both (purple) and neither (gray). The *R* represents the correlation coefficients of the treatment effects on gene mean expression and gene bulk expression, *p ≥* 0.05 (NS), *p <* 0.05 (*), *p <* 0.01 (**) and *p <* 0.001(***).

**Figure S16:**
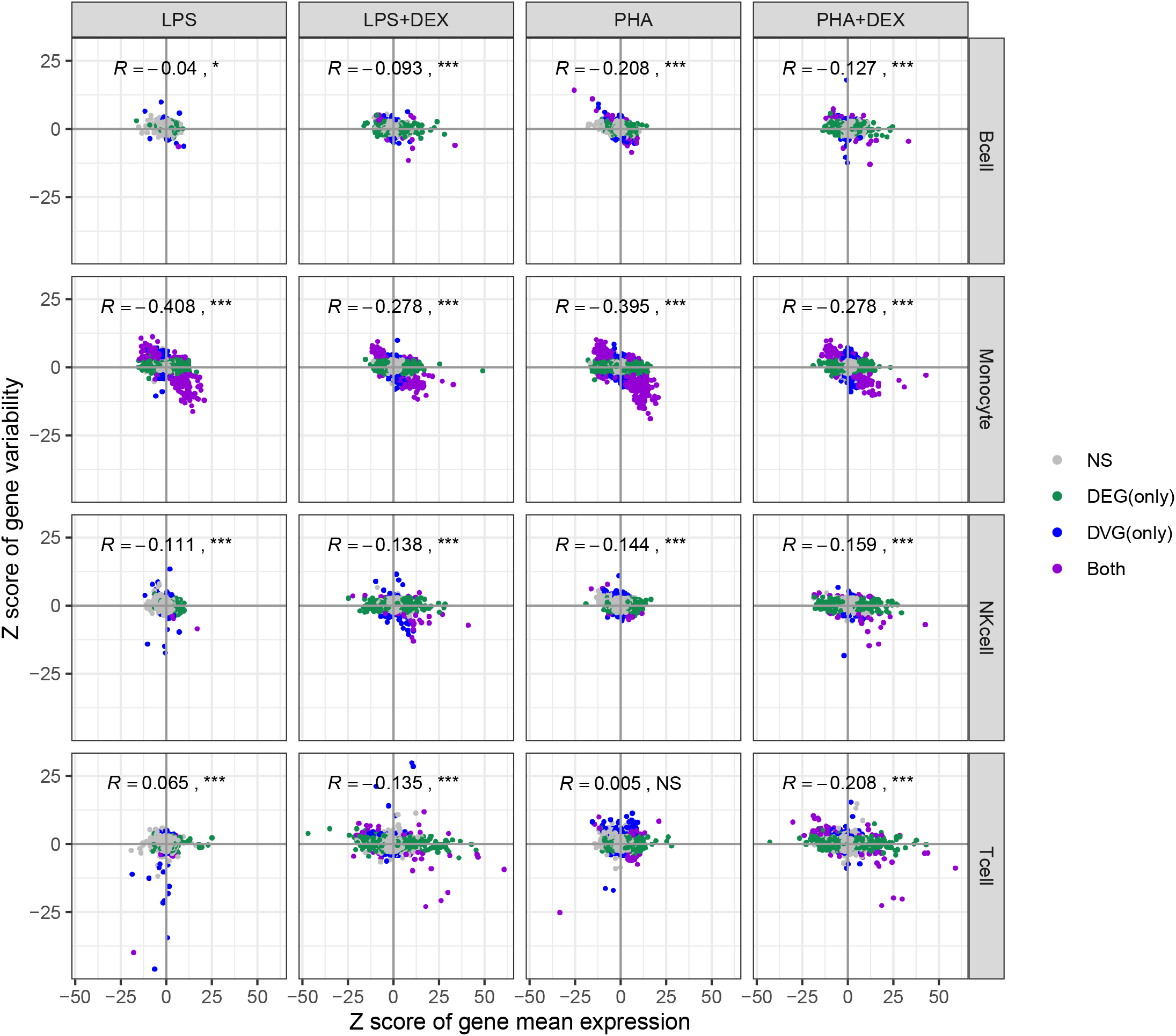
The scatter plots of the z score of treatment effects on gene mean (x axis) against the z score of treatment effects on gene variability (y axis): Color represents significant treatment effects on gene mean only (green), gene variability only (blue), both (purple) and neither (gray). The *R* represents the correlation coefficients of the treatment effects on gene mean and gene variability, *p ≥* 0.05 (NS), *p <* 0.05 (*), *p <* 0.01 (**) and *p <* 0.001(***).

**Figure S17:**
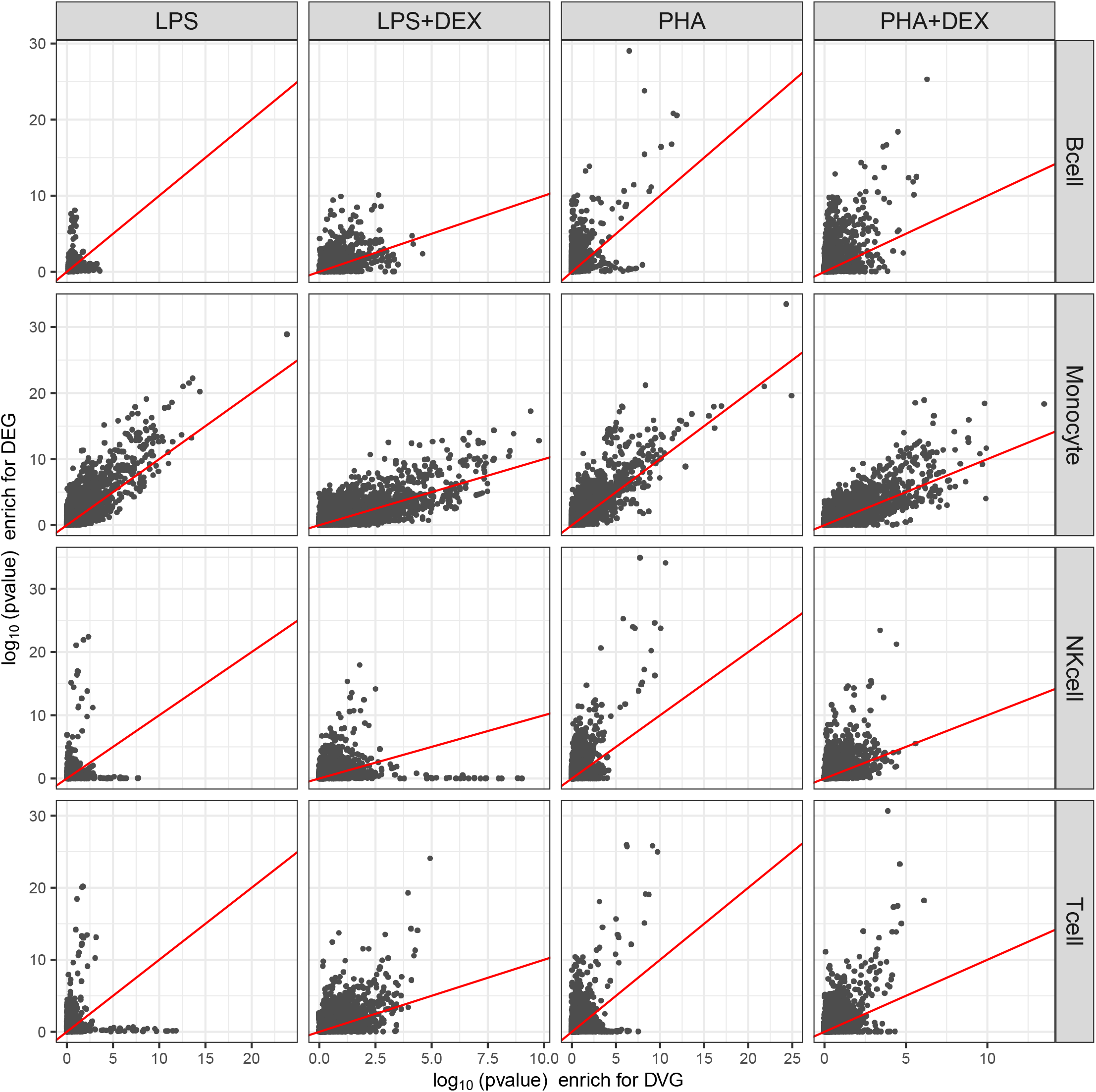
**Comparison of the results of GO enrichment between DEG and DVG**, x axis representing *−* log_10_(*p*) of the GO terms that are enriched by DVG and Y axis representing *−* log_10_(*p*) of the GO terms that are enriched by DEG.

**Figure S18:**
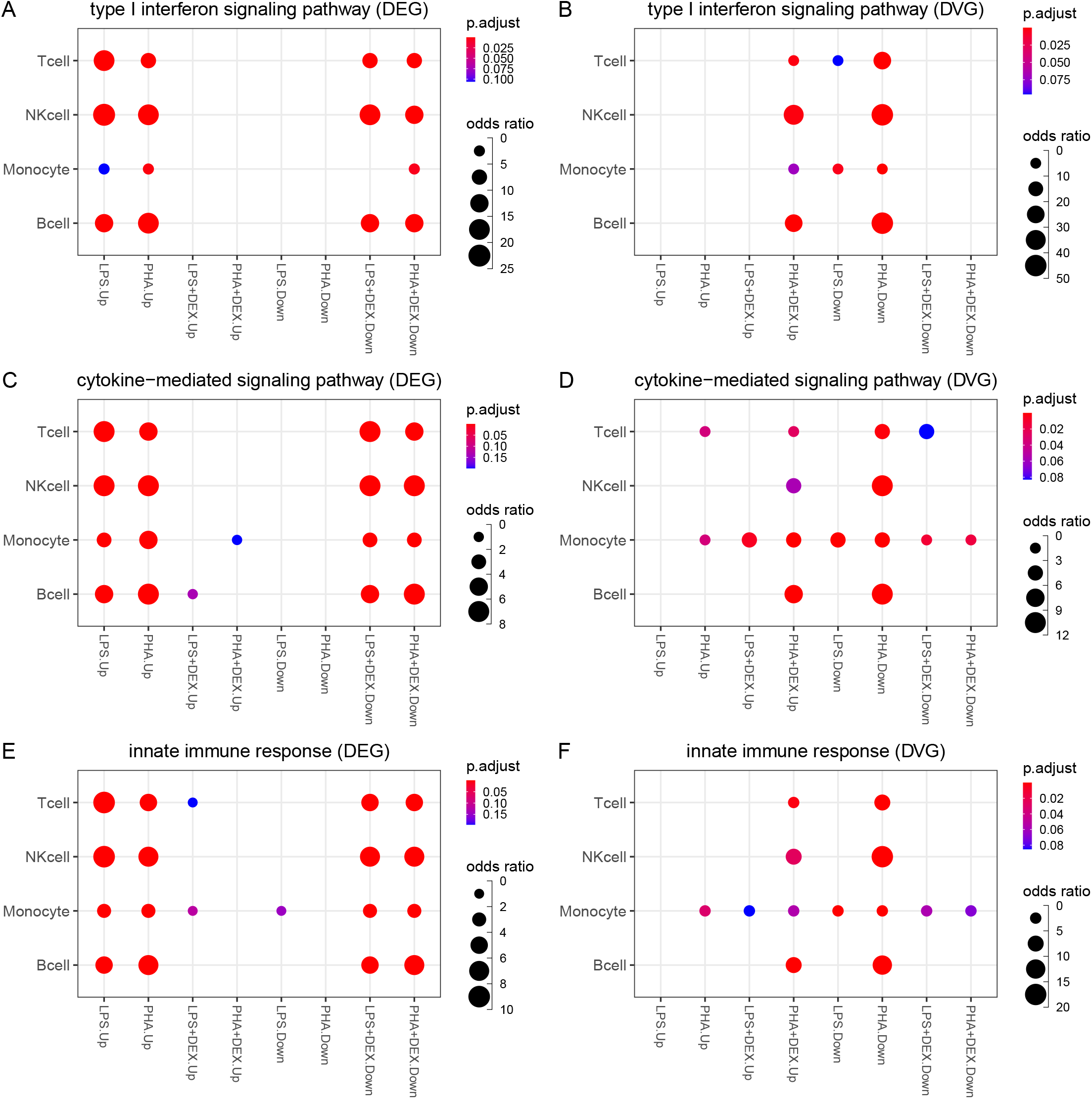
Examples of shared enrichment pathways for DEG and DVG: **(A and B)** represent enrichment results in the type I interferon signaling pathway for DEG and DVG, respectively, **(C and D)** represent enrichment results in the cytokine-mediated signaling pathway for DEG and DVG, respectively, and **(E and F)** represent enrichment results in innate immune response for DEG and DVG, respectively.

**Figure S19:**
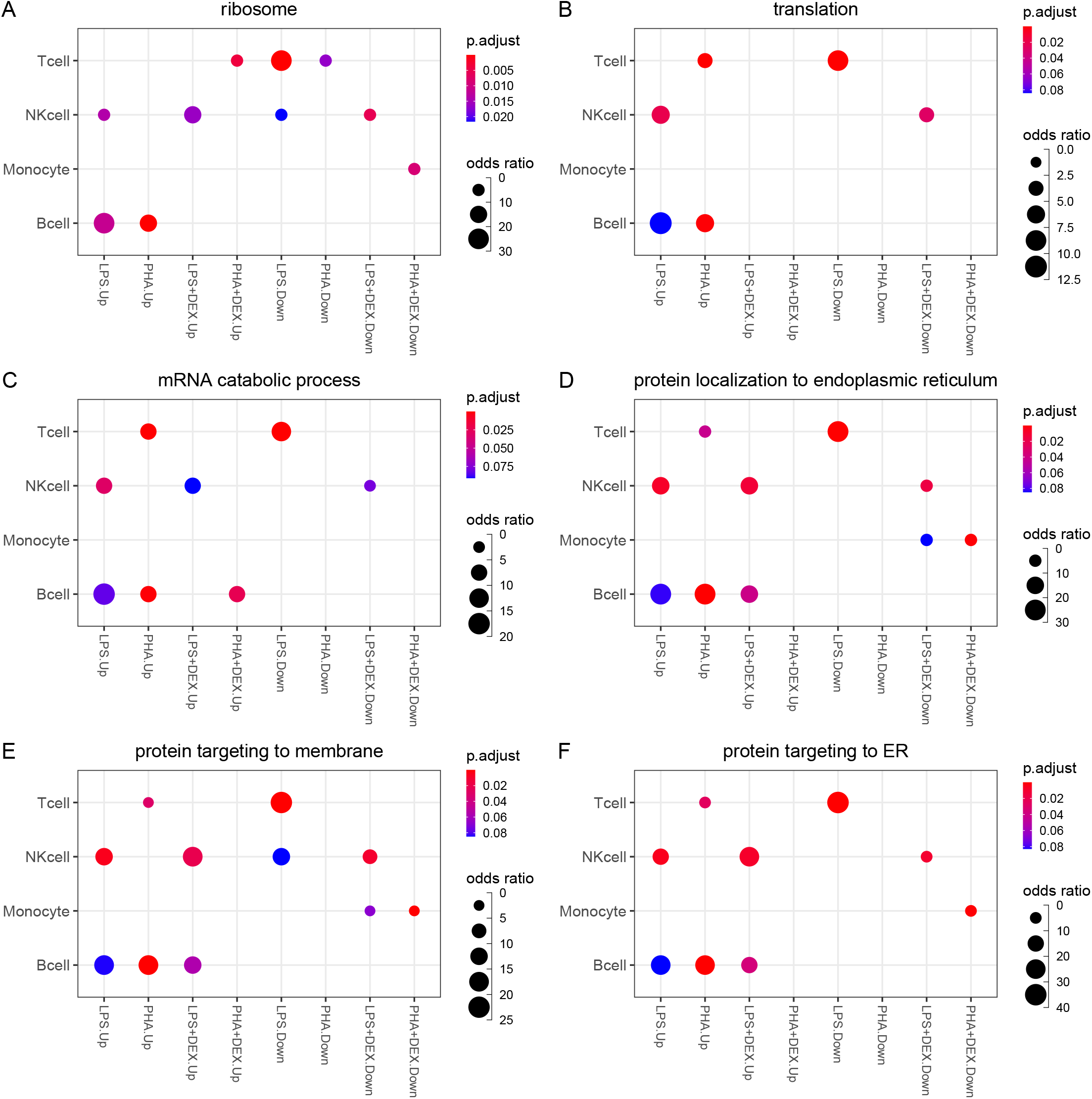
Examples of the pathways that are only enriched in DVGs: **(A-F)** represent enrichment results in ribosome, translation, mRNA catabolic process, protein localization to endoplasmic reticulum, protein targeting to membrane and protein targeting to ERrespectively.

**Figure S20:**
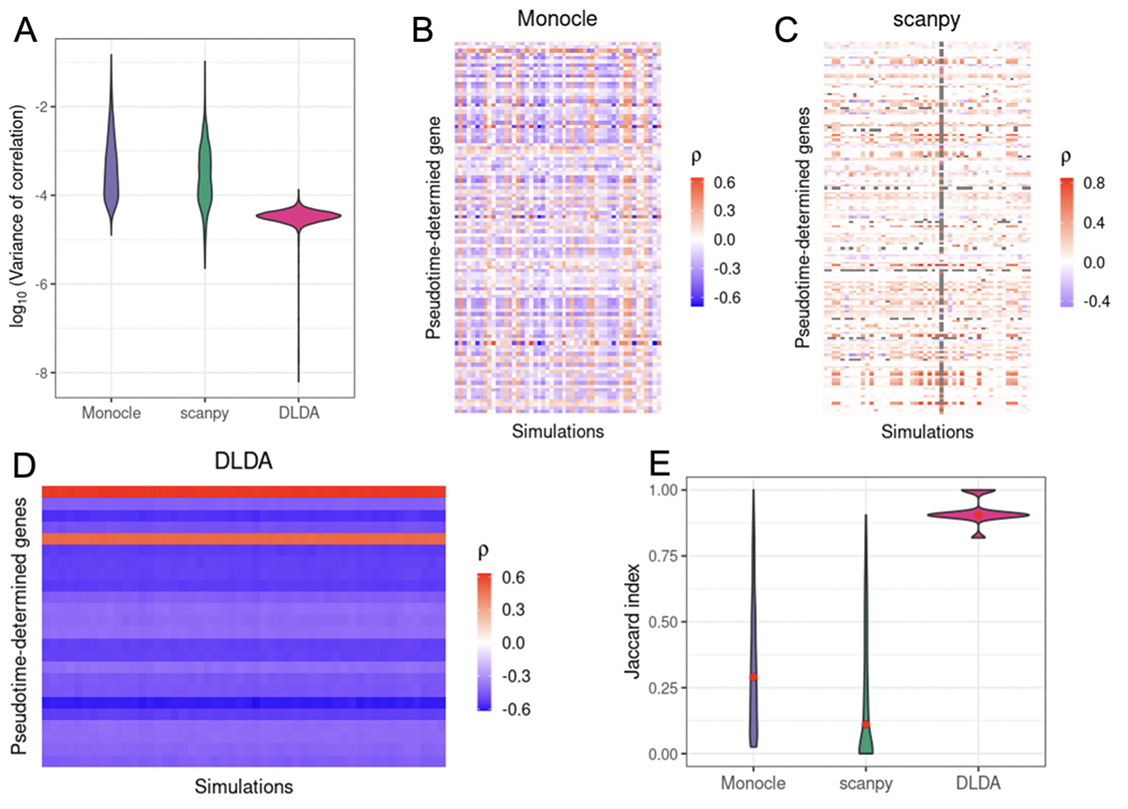
Validation of effectiveness and robustness of the treatment pseudotime trajectory using resampling. **(A)** Violin plot of the variance of correlation coefficients for each gene across 50 resamplings from Monocle, scanpy and DLDA. **(B-D)** Heatmaps of the correlation coefficients of top 20 ranking pseudotime driving genes across resampling. Each row represents the same gene and each column a resampling iteration with the color representing the magnitude of the correlation between gene expression and the derived pseudo-time variable; ideally we would like to see that correlation be approximately constant. The resamplings are from **(B)** Monocle, **(C)** scanpy and **(D)**DLDA, respectively. **(E)** Violin plot of jaccard index of the stability of the top 20 pseudotime correlated genes for each pair of resamplings across 1,225 pairwise comparisons from Monocle, scanpy and DLDA.

**Figure S21:**
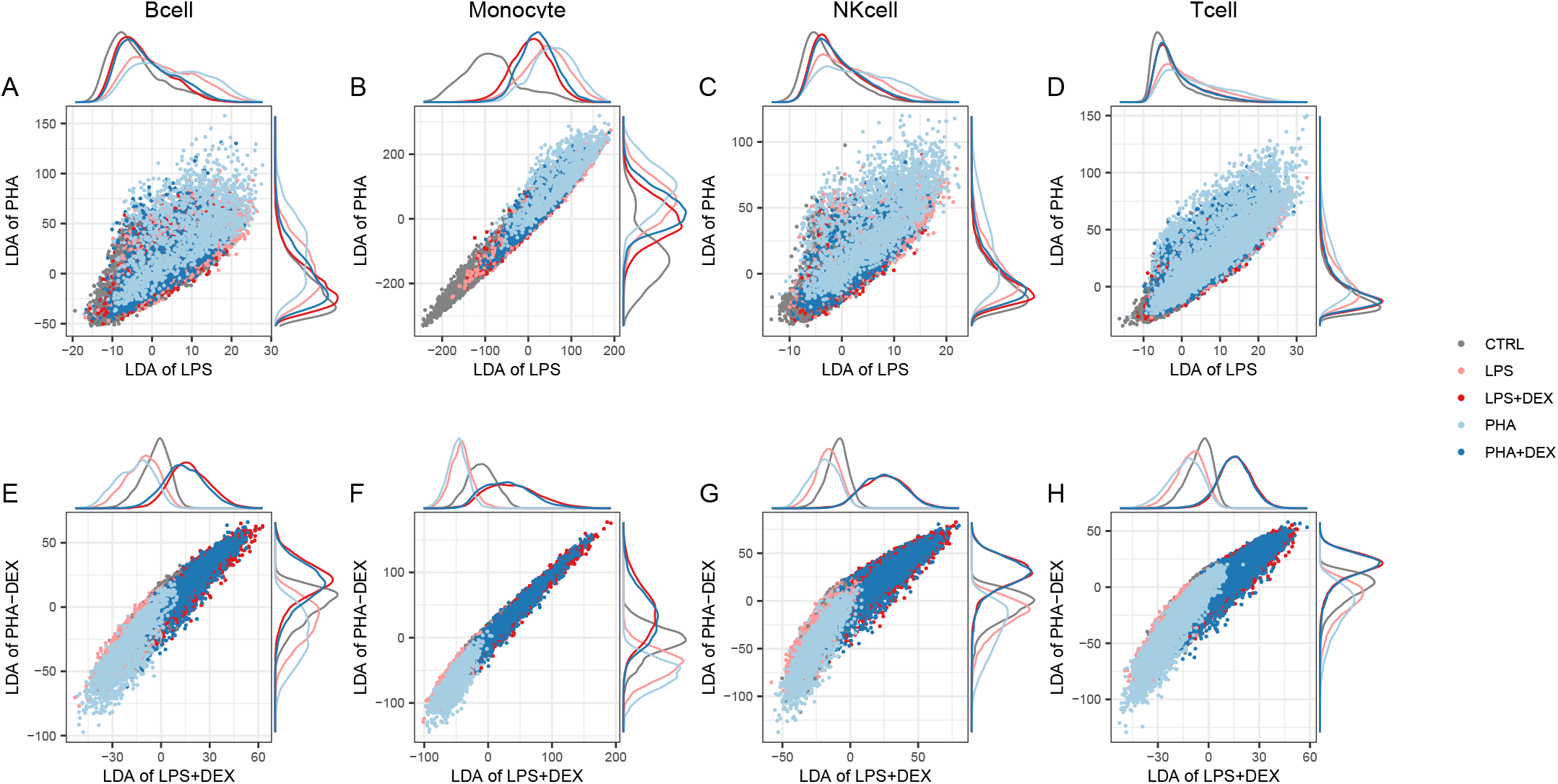
Low dimensional manifold plots from DLDA across cell types: **(A-D)** represent the scatter plots of the DLDA from LPS (x axis) against the DLDA from PHA (y axis) and **(E-H)** represent the scatter plots of the DLDA from LPS+DEX (x axis) against PHA+DEX (y axis).

**Figure S22:**
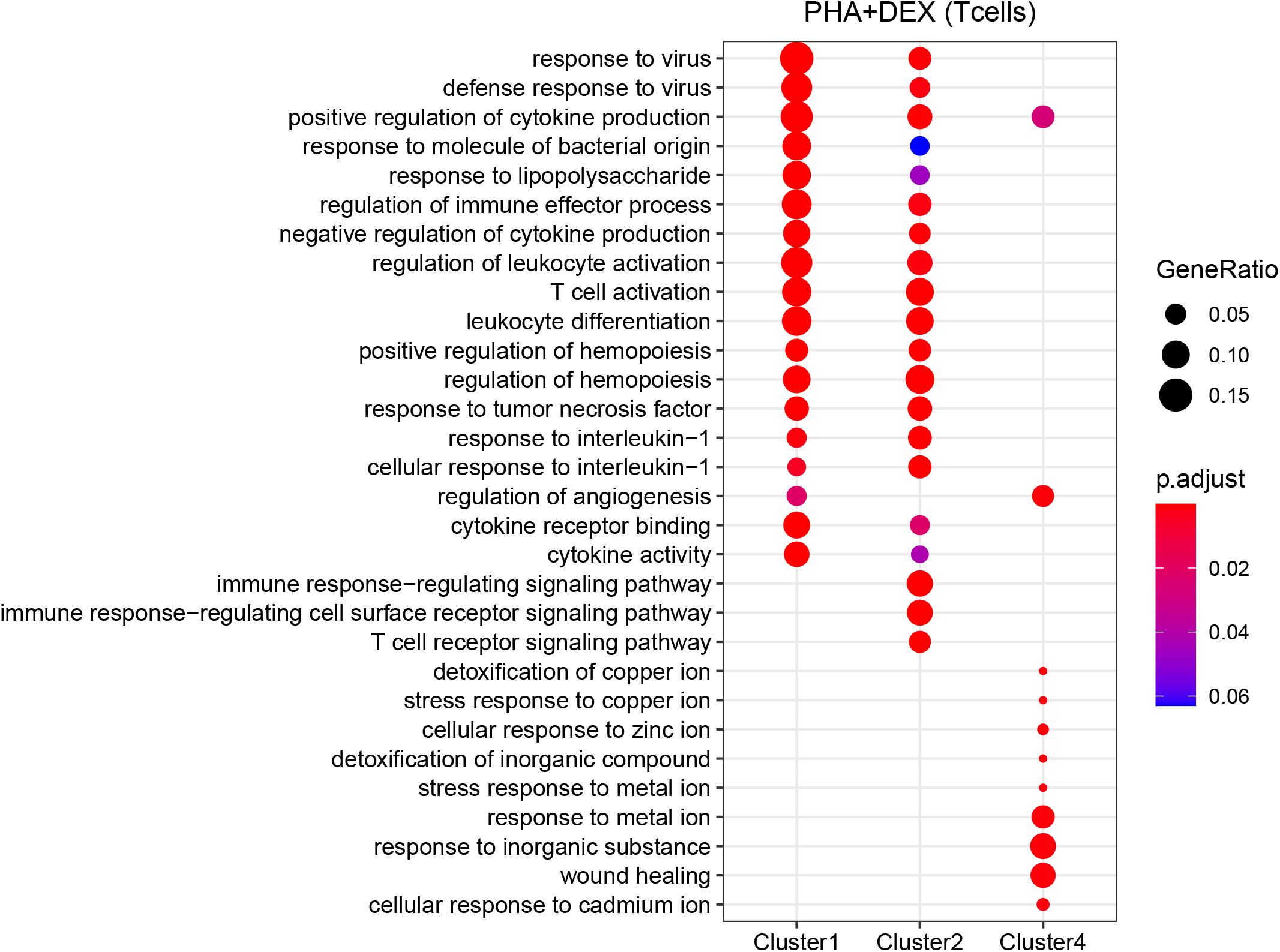
Gene ontology enrichment results for four dynamic patterns DEGs respectively.

**Figure S23:**
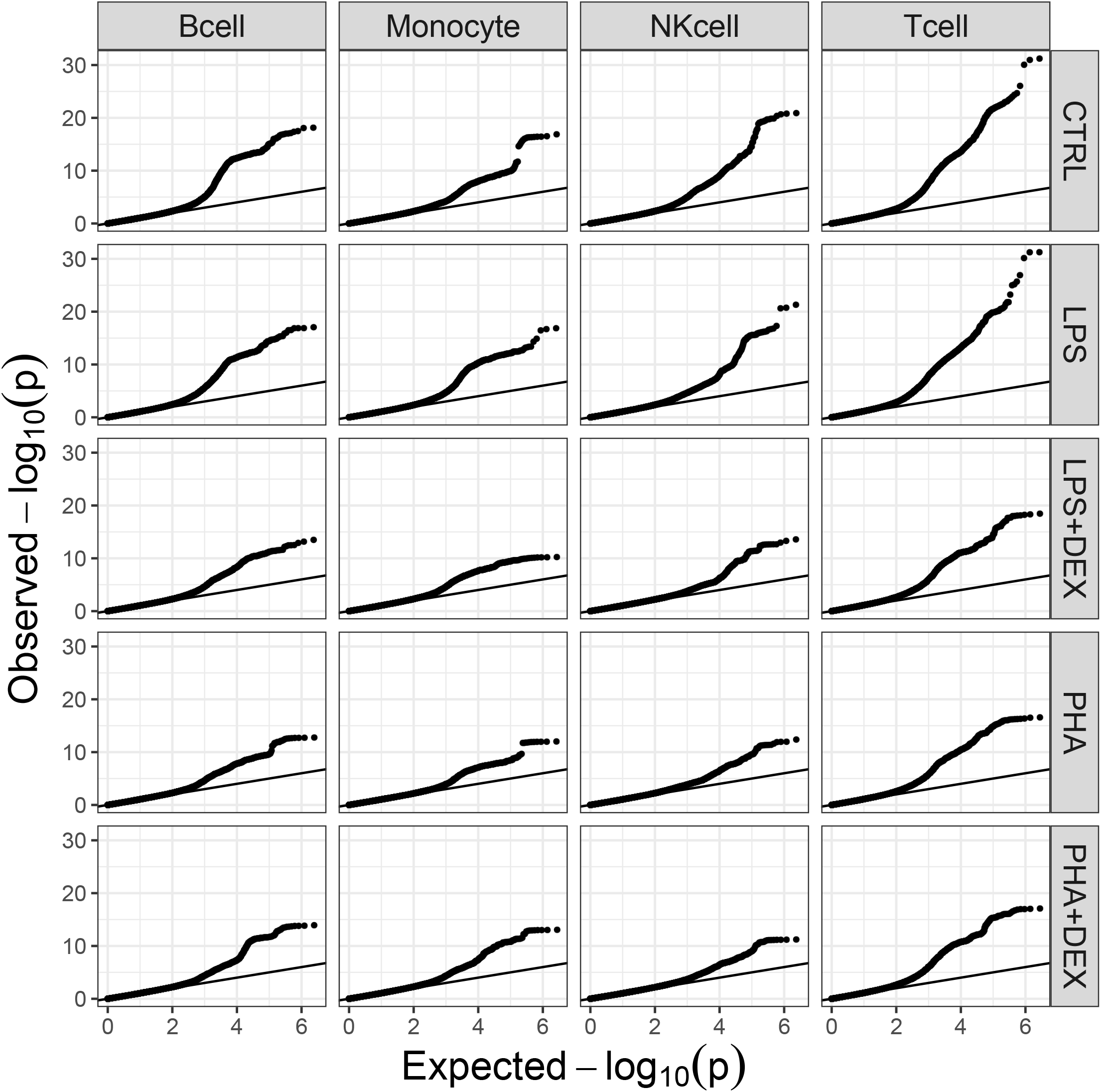
QQplots of p-values from eQTL mapping with FastQTL on pseudobulk gene expression data across the 20 conditions.

**Figure S24:**
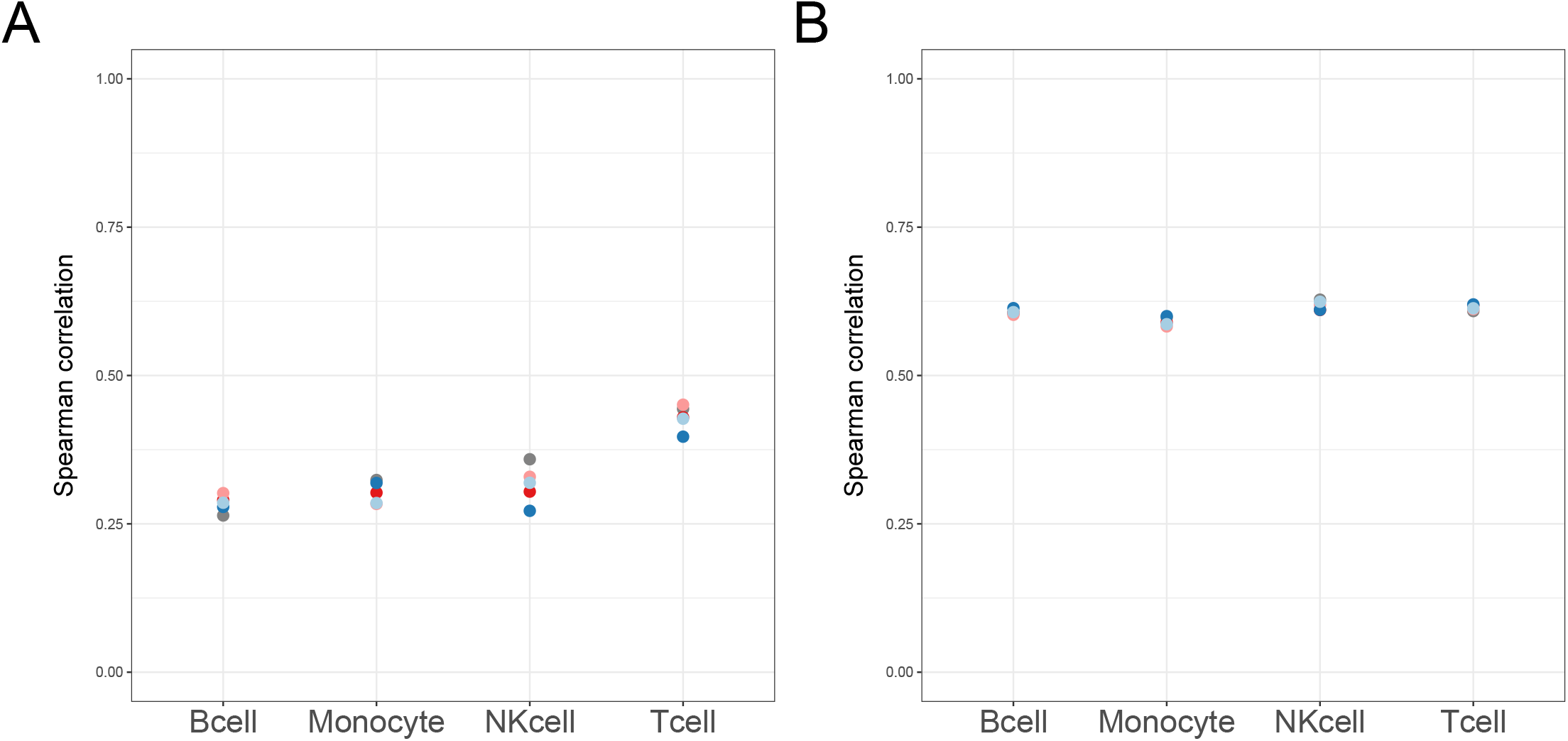
Correlations between genetic effect sizes assessed in single-cell data and in bulk RNA-seq. [93] **A** Spearman correlations for significant genetic effect sizes assessed by condition-by-condition analysis (FDR*<* 0.1) **B** Spearman correlations for significant genetic effect sizes assessed by multivariate adaptive shrinkage analysis (LFSR*<* 0.1). Color denotes treatment: grey – control, light blue – PHA, dark blue – PHA+DEX, pink – LPS, red – LPS+DEX. All values are significant (p-value*<* 0.05).

**Figure S25:**
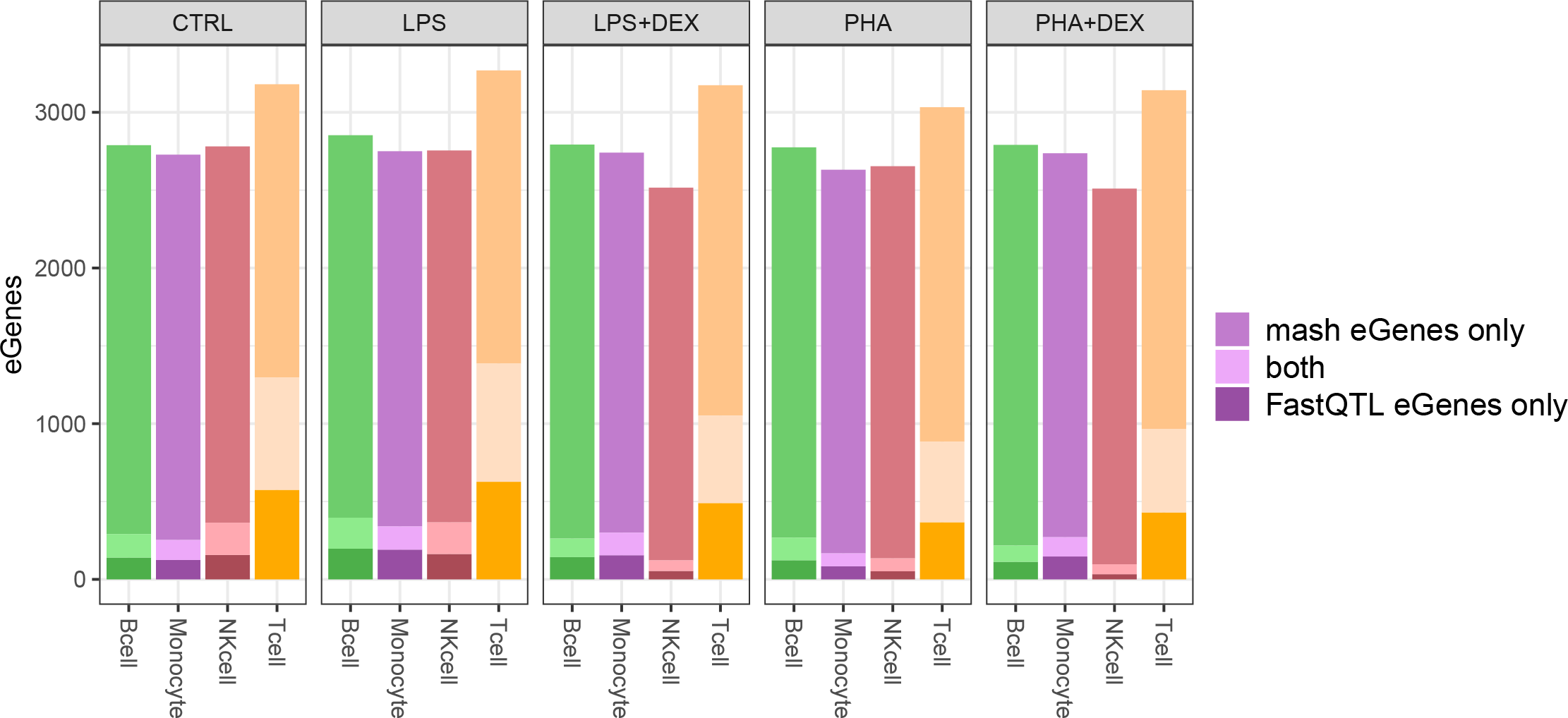
eQTL mapping results. Barplot represents the number of genes with eQTLs (eGenes) significant in only condition-by-condition analysis (FDR*<* 0.1, dark shading), significant in multivariate adaptive shrinkage analysis only (LFSR*<* 0.1, medium shading), and significant in both analyses (light shading).

**Figure S26:**
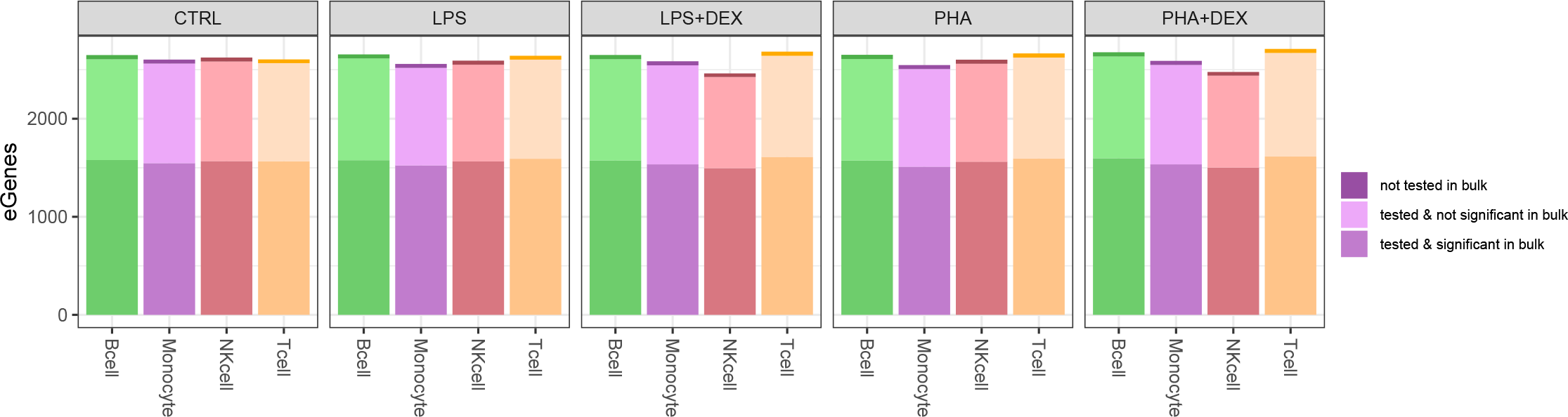
Barplot represents the number of genes with significant eQTLs as assessed by multivariate adaptive shrinkage analysis (eGenes, LFSR*<* 0.1). Shading denotes significance in bulk dataset: significant in bulk RNA-seq data citepmid34142656 (dark shading), not tested in bulk RNA-seq data (light shading), and not significant in bulk RNA-seq data (medium shading).

**Figure S27:**
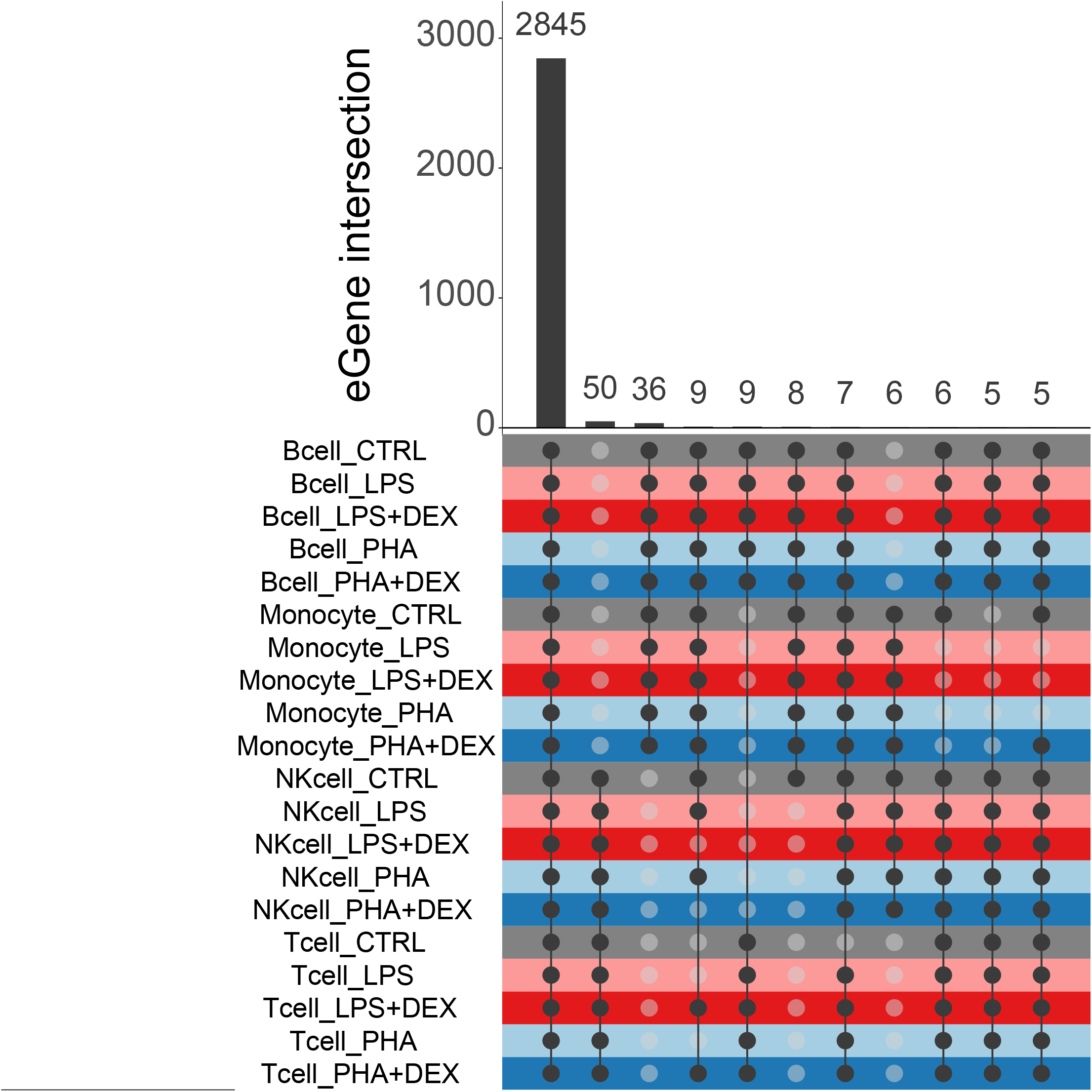
eGene sharing by sign across all conditions. Upset plot represents the number of eGenes with eQTLs shared by sign (i.e., with genetic effects in the same direction) for each unique set of conditions with intersection size of at least 5.

**Figure S28:**
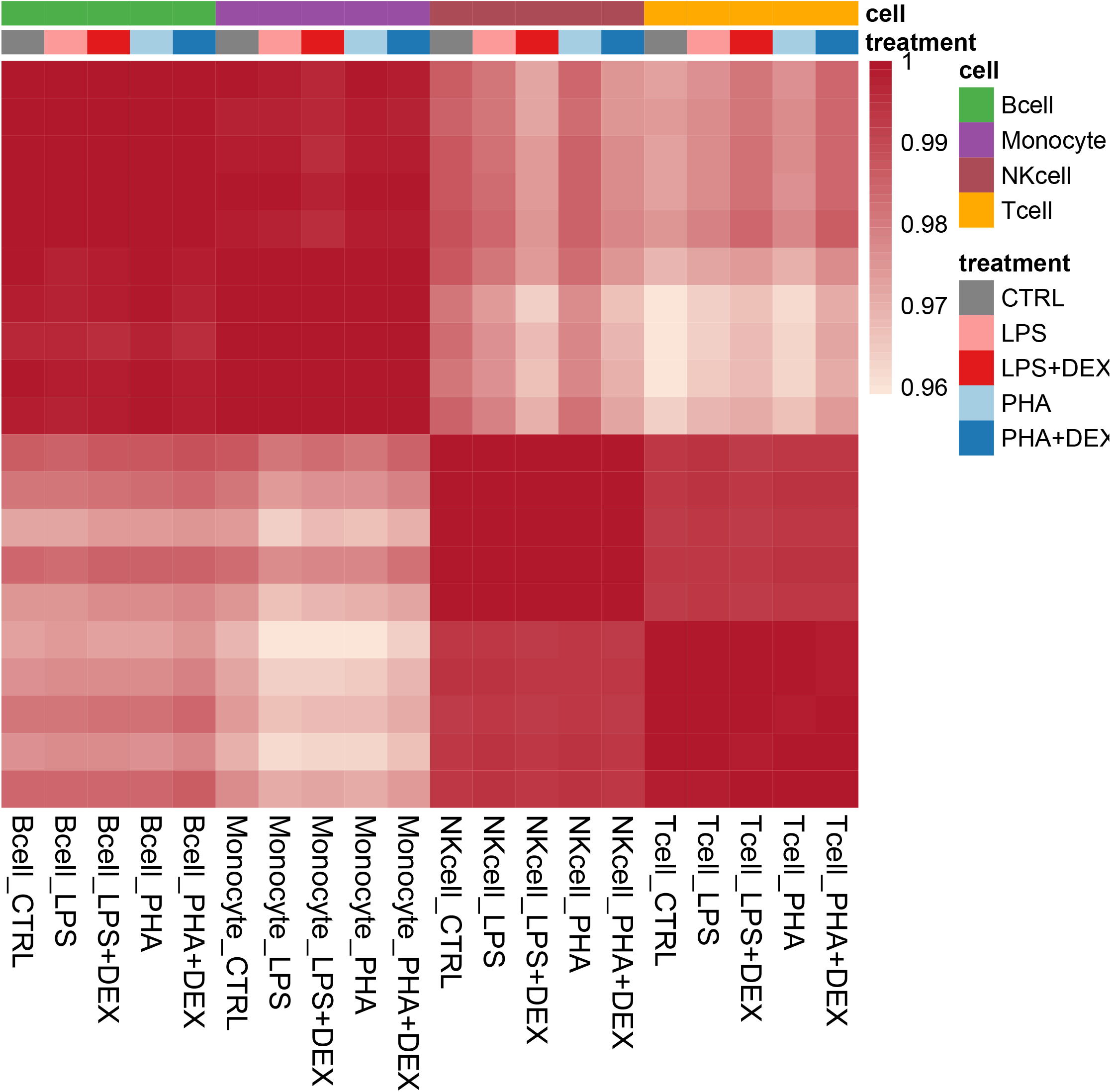
eGene differences in the direction of effect across all pairs of conditions. Heatmap represents proportion of eGenes significant in either of the two conditions which have at least one eQTL significant in either of the two conditions with opposite sign of genetic effect size estimated by multivariate adaptive shrinkage.

**Figure S29:**
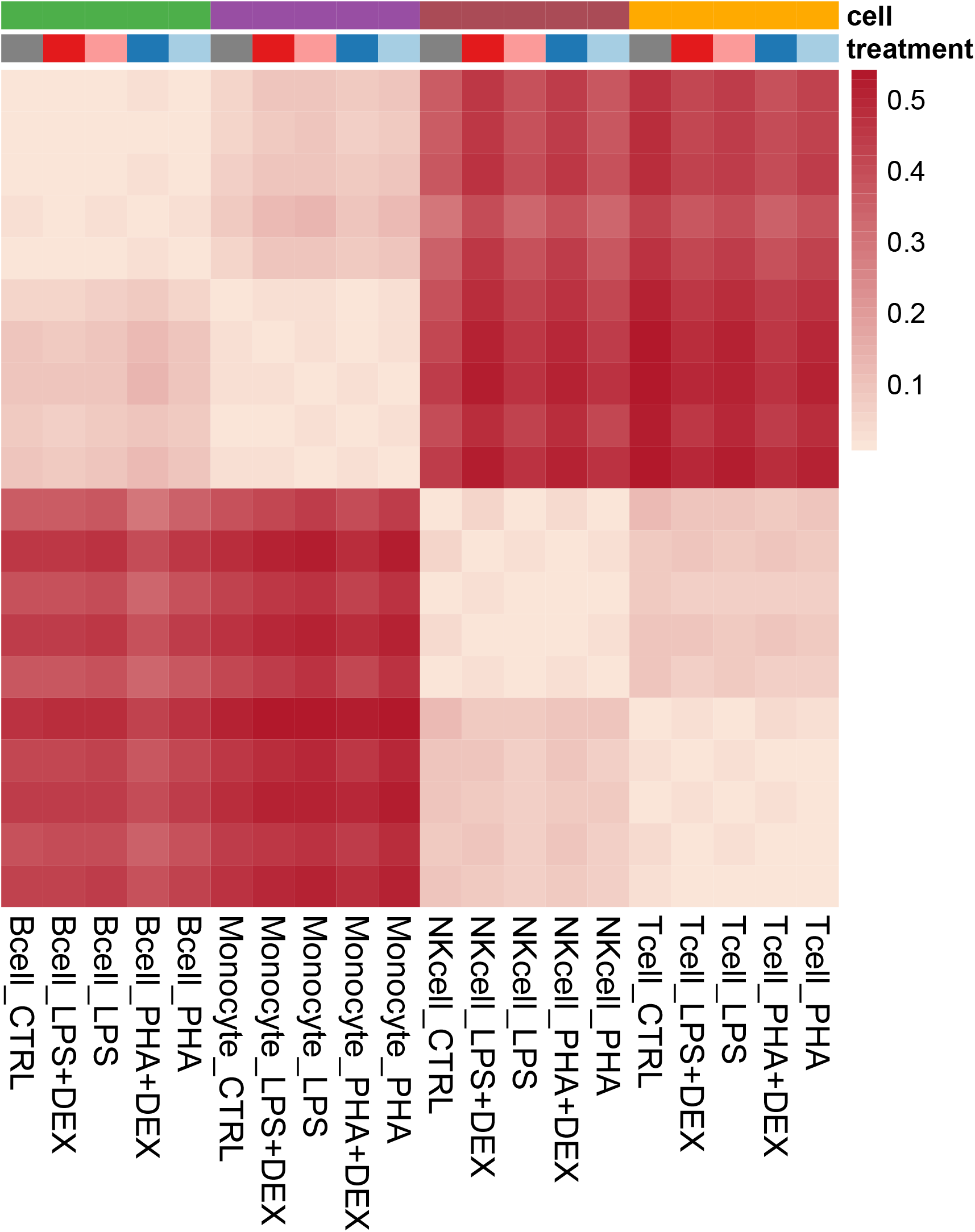
eGene differences across all pairs of conditions. Heatmap represents proportion of eGenes significant in either of the two conditions which have at least one eQTL significant in either of the two conditions with at least two-fold difference of genetic effect size estimated by multivariate adaptive shrinkage.

**Figure S30:**
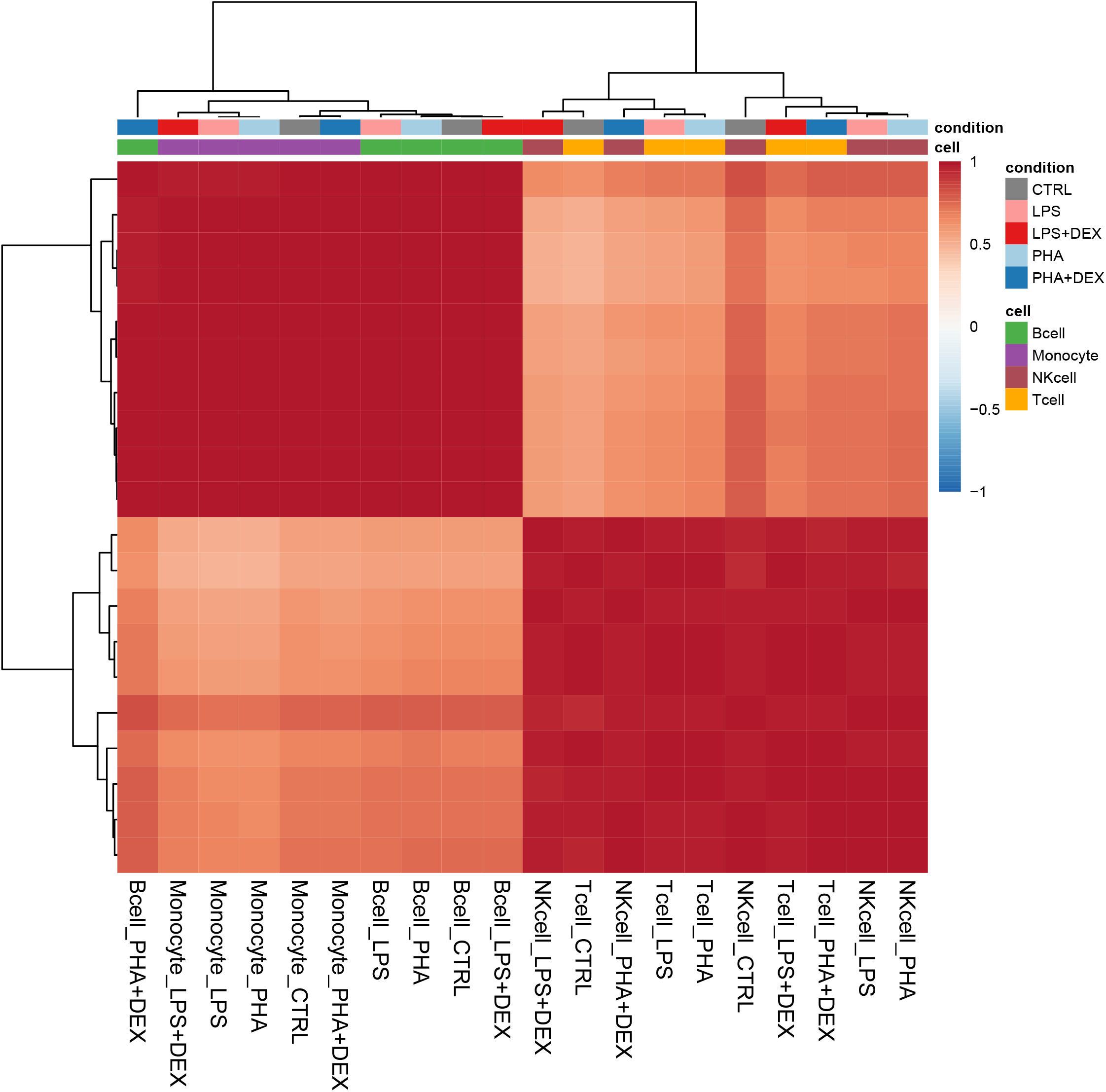
Correlations between genetic effect sizes estimated with multivariate adaptive shrinkage. Heatmap represents pair-wise Pearson correlations based on union of significant eQTLs in either of the two conditions (LFSR*<* 0.1). All values are significant (p-value*<* 0.05).

**Figure S31:**
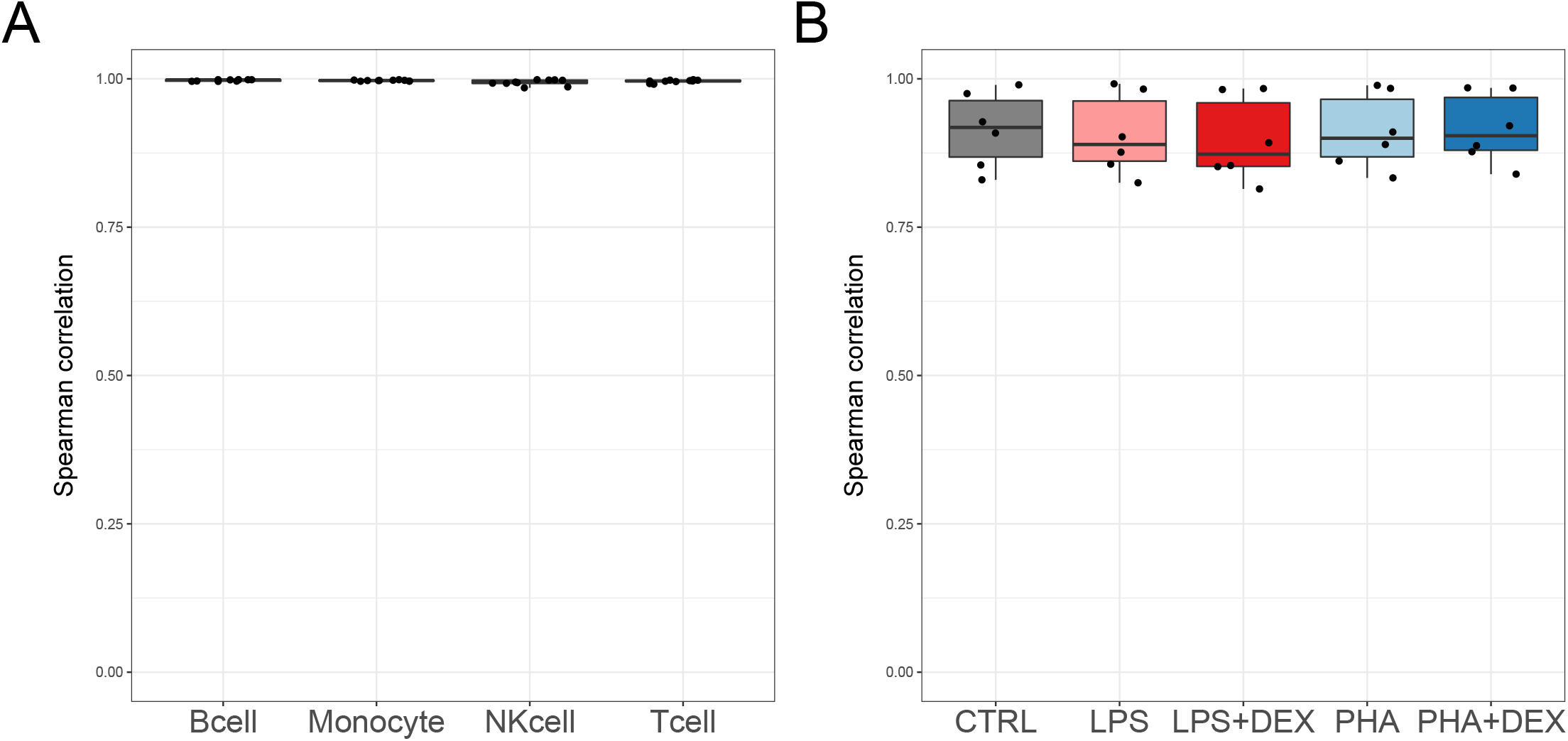
Spearman correlations between significant genetic effect sizes estimated with multivariate adaptive shrinkage (LFSR*<* 0.1). **A** pair-wise correlations between all treatment conditions within each cell type **B** pair-wise correlations between all cell types within each treatment. All values are significant (p-value*<* 0.05).

**Figure S32:**
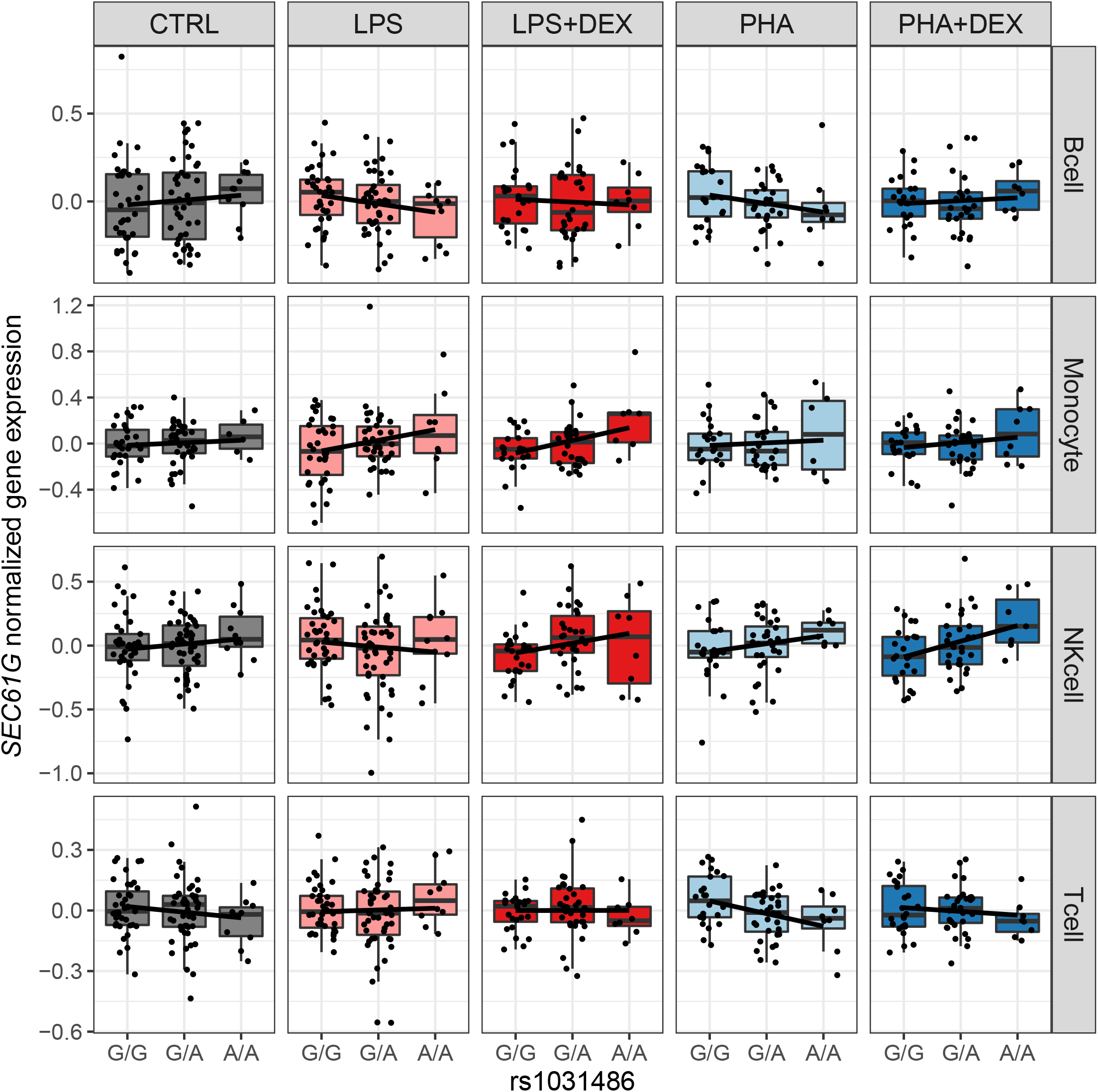
Genetic effect of rs1031486 on gene expression of *SEC61G*. Boxplots represent normalized expressions of the *SEC61G* gene (y axis) across the three genotype classes of rs1031486 (x axis) across all conditions.

**Figure S33:**
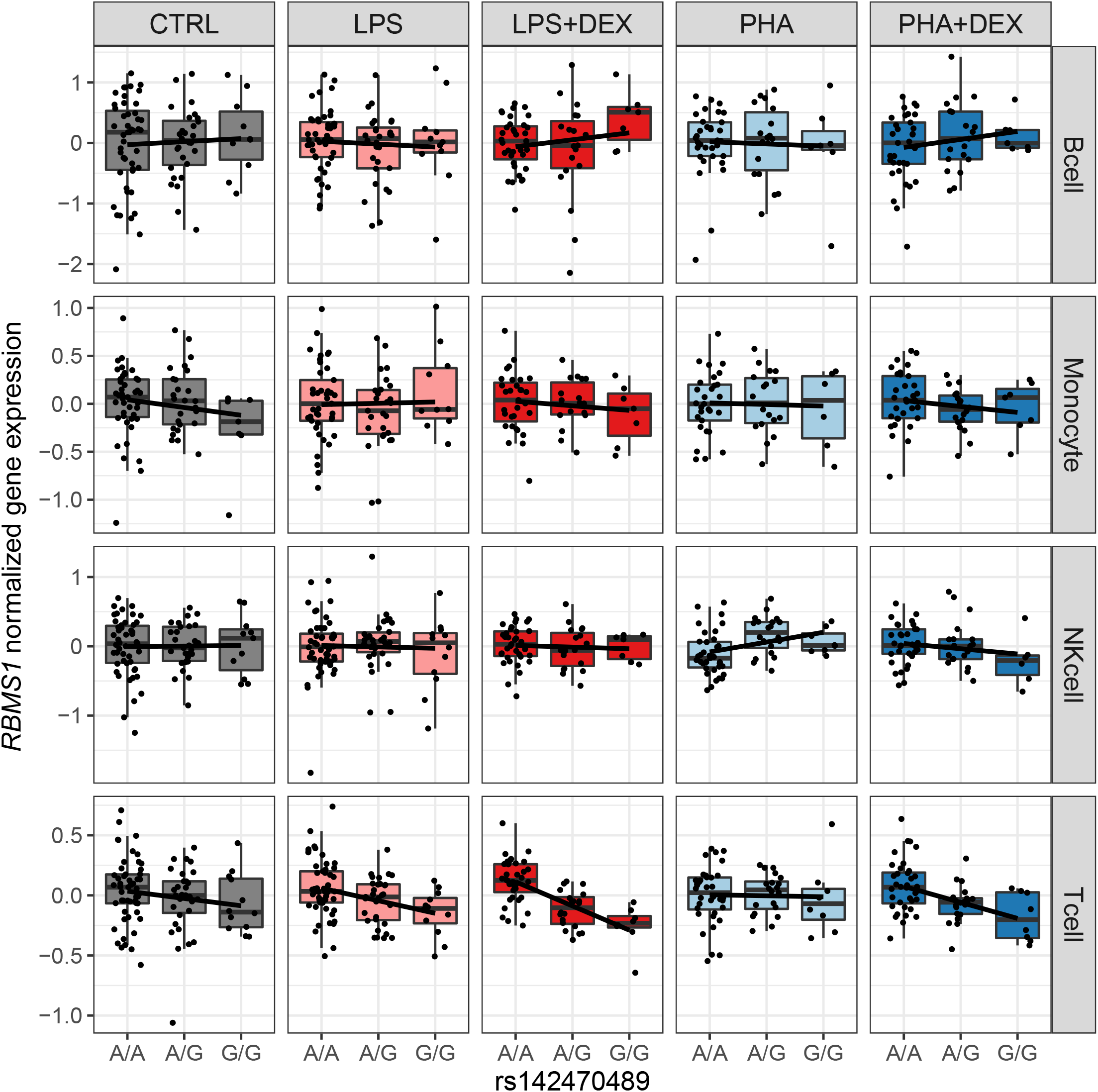
Genetic effect of rs142470489 on gene expression of *RBMS1*. Boxplots represent normalized expressions of the *RBMS1* gene (y axis) across the three genotype classes of rs142470489 (x axis) across all conditions.

**Figure S34:**
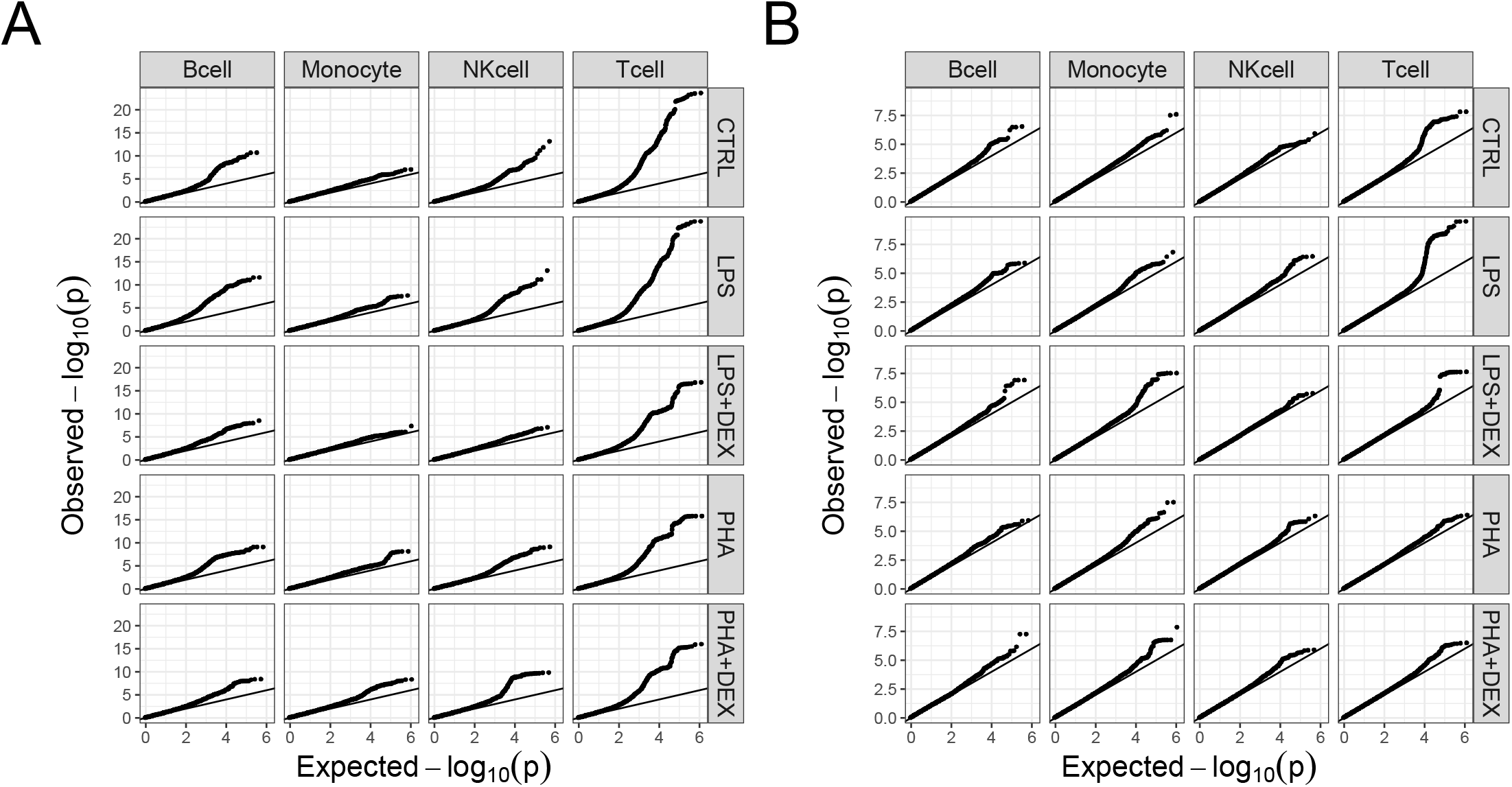
QQplots for genetic analyses of mean and variability. **A** p-values from mean-eQTL mapping with FastQTL **B** p-values from variability QTL mapping with FastQTL.

**Figure S35:**
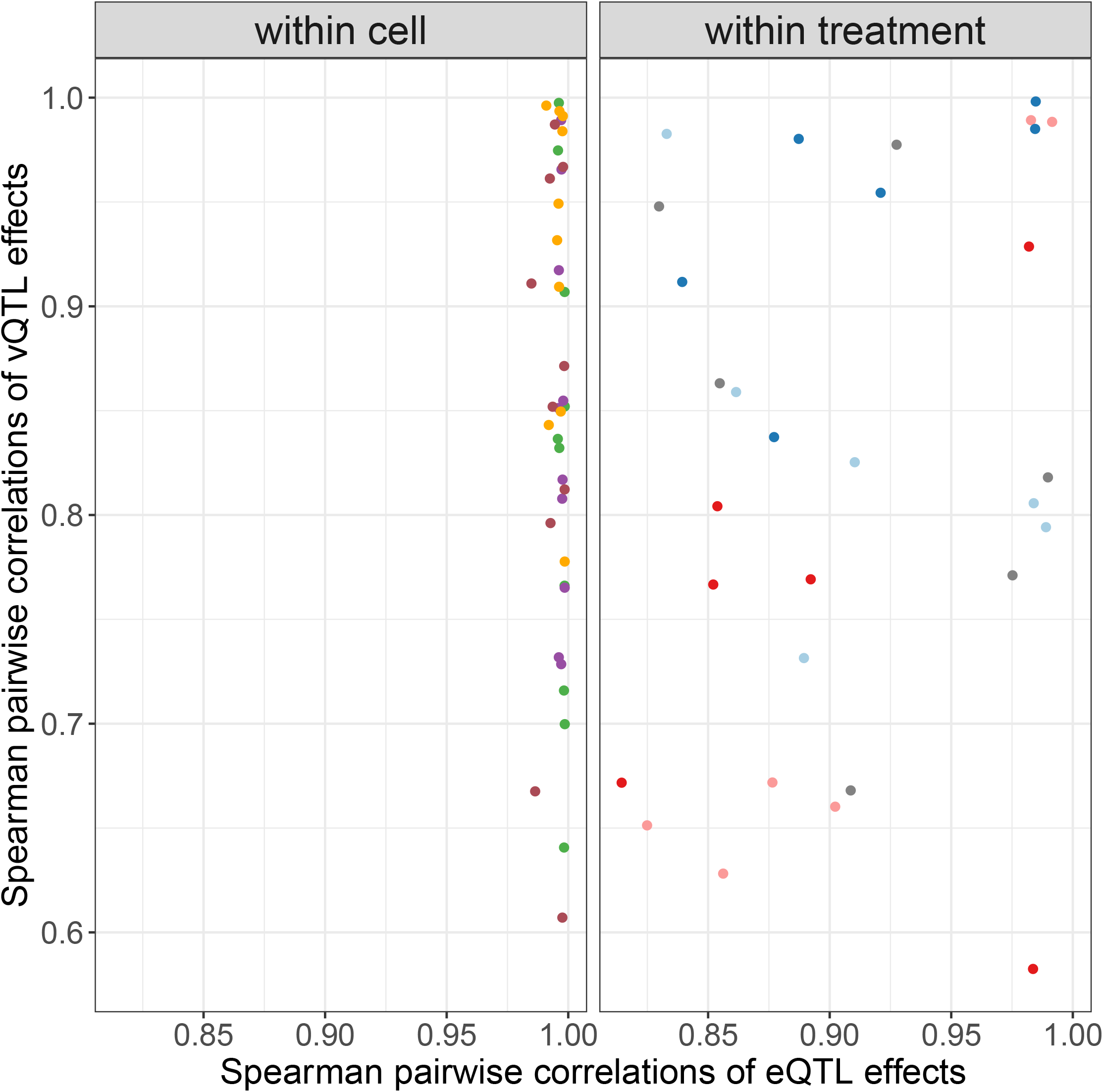
Cell type- and treatment-specificity of genetic effects on gene expression and gene expression variability. Scatterplots represent pairwise Spearman correlations between significant genetic effects (LFSR*<* 0.1) on gene expression (x axis), and gene expression variability (y axis) across all treatment conditions within each cell type (left panel) and across all cell types within each treatment condition (right panel).

**Figure S36:**
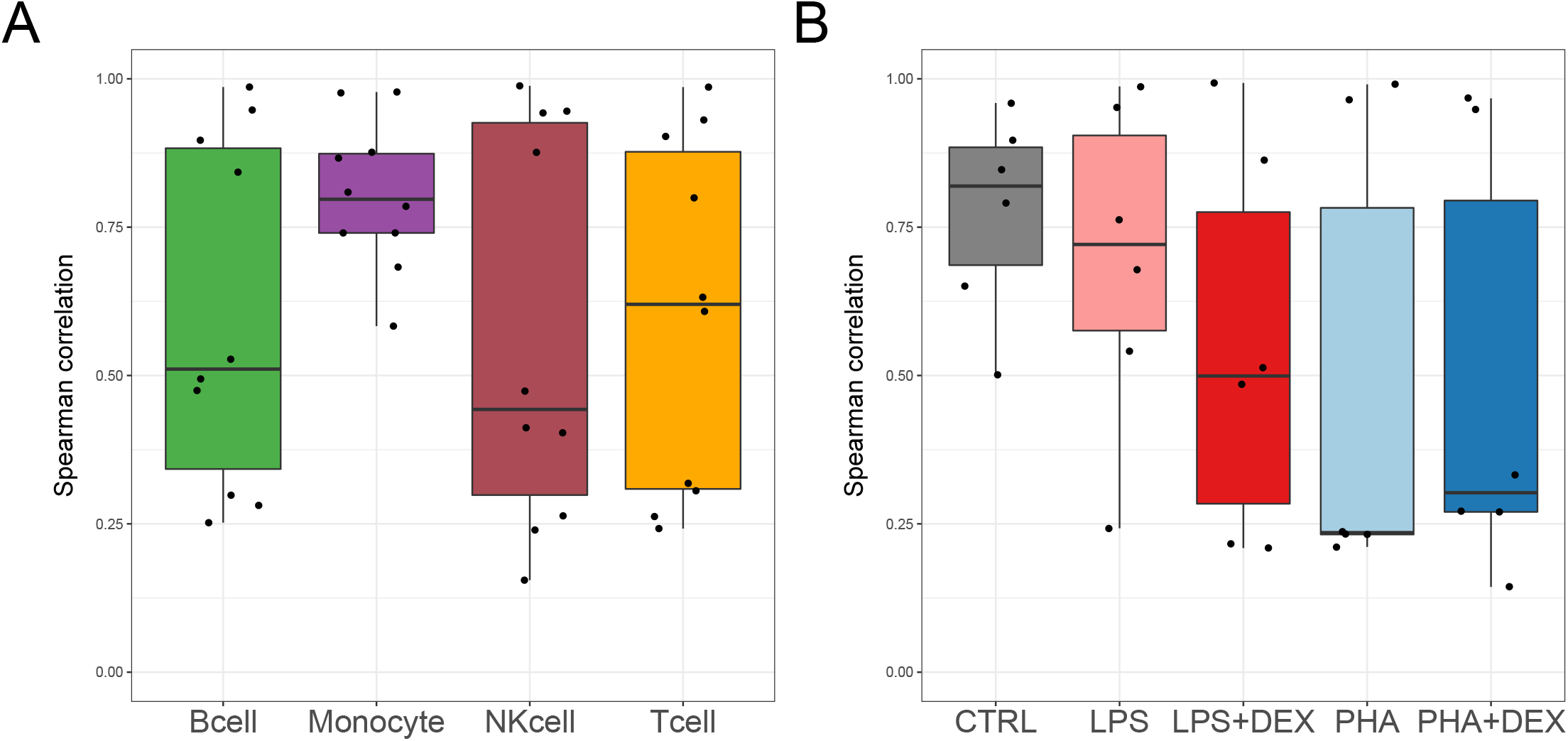
Spearman correlations between significant genetic effects on gene expression variability estimated with multivariate adaptive shrinkage (LFSR*<* 0.1). **A** – pair-wise correlations between all treatment conditions within each cell type **B** – pair-wise correlations between all cell typeswithin each treatment condition

**Figure S37:**
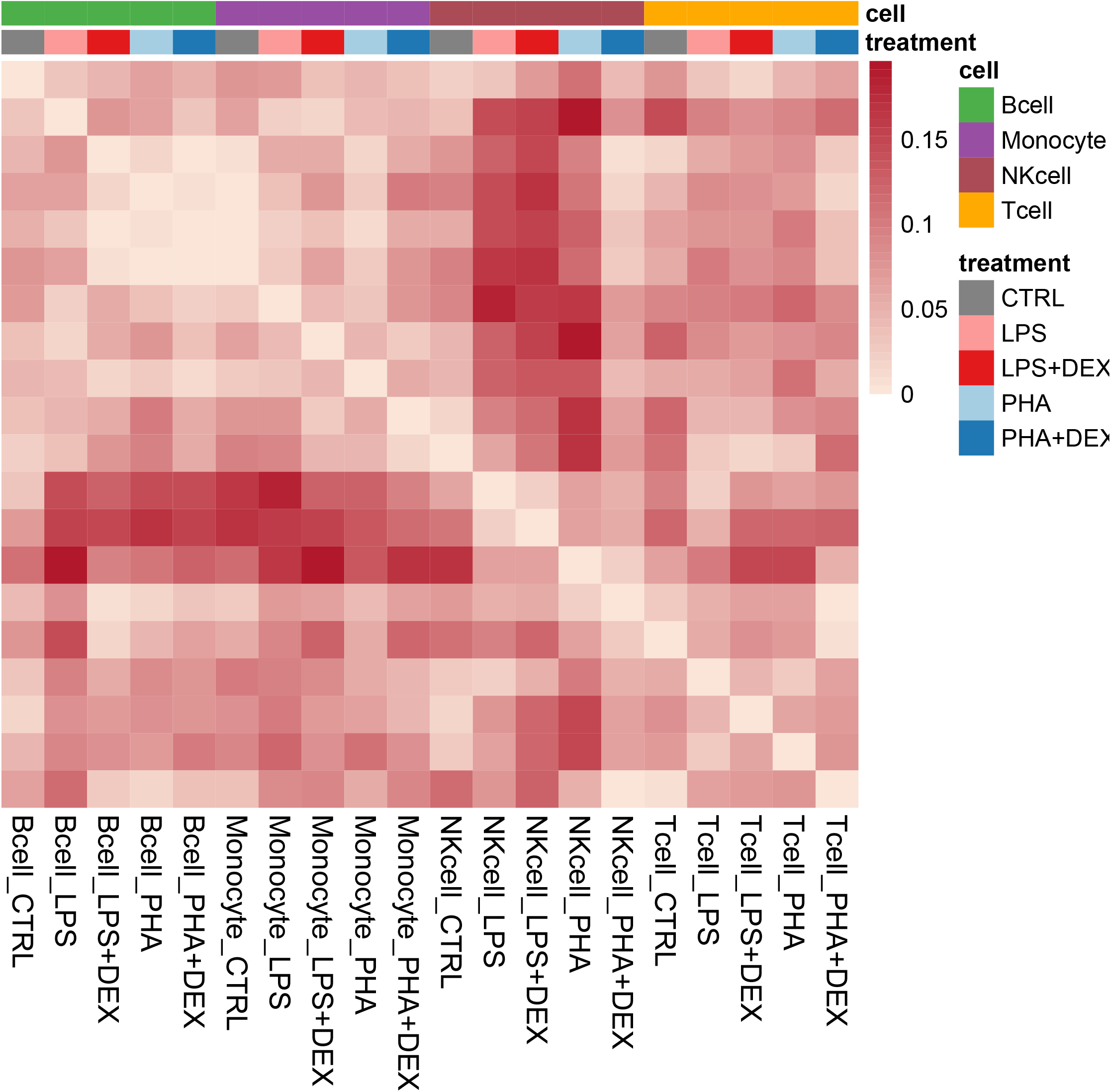
vGene differences in the direction of effect across all pairs of conditions. Heatmap represents proportion of vGenes significant in either of the two conditions which have at least one vQTL significant in either of the two conditions with opposite sign of genetic effect size estimated by multivariate adaptive shrinkage.

**Figure S38:**
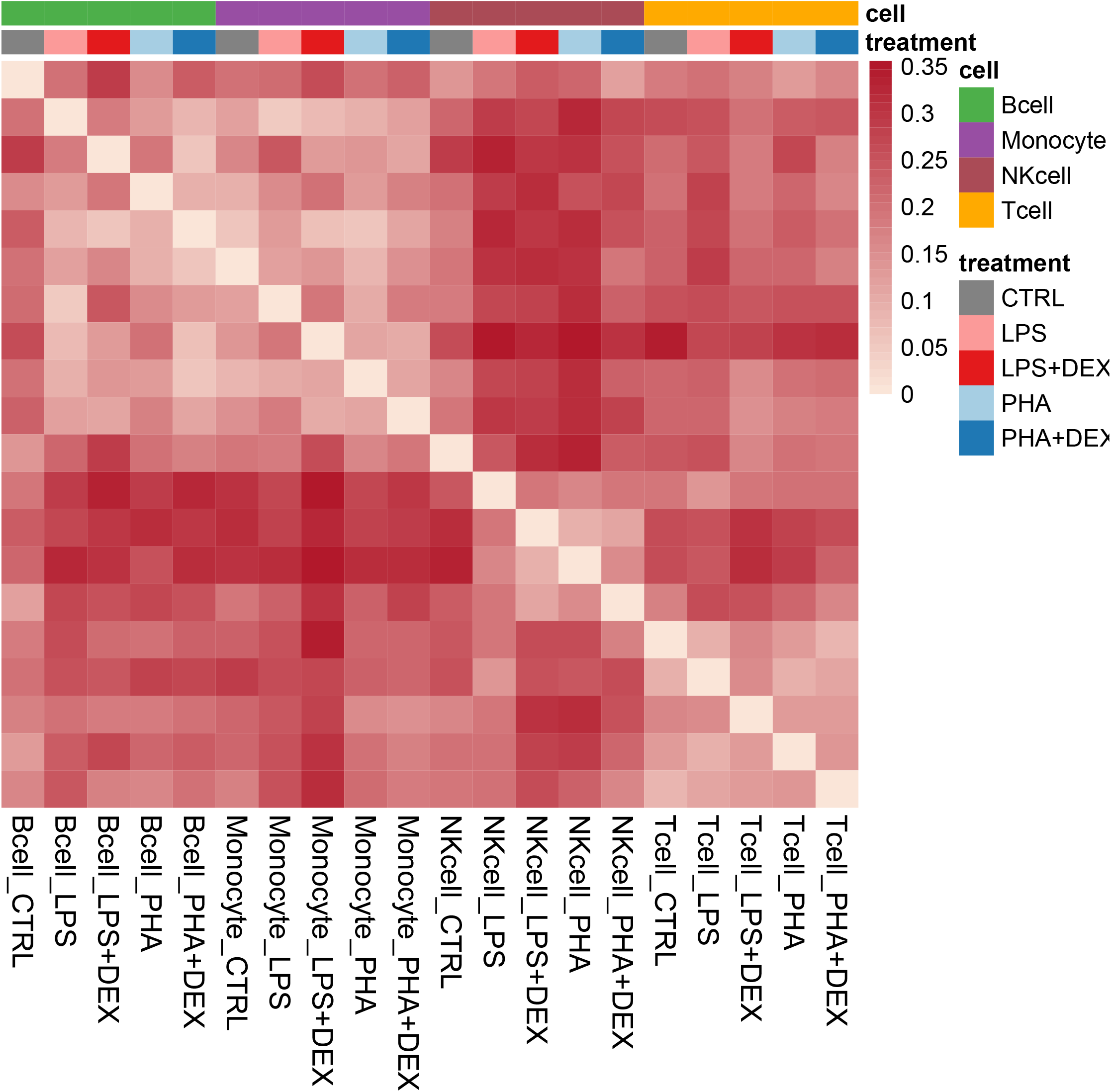
vGene differences across all pairs of conditions. Heatmap represents the proportion of vGenes not shared by magnitude (i.e., with at least 2-fold difference between genetic effects in each condition) over the union of all significant vGenes in each pair-wise comparison.

**Figure S39:**
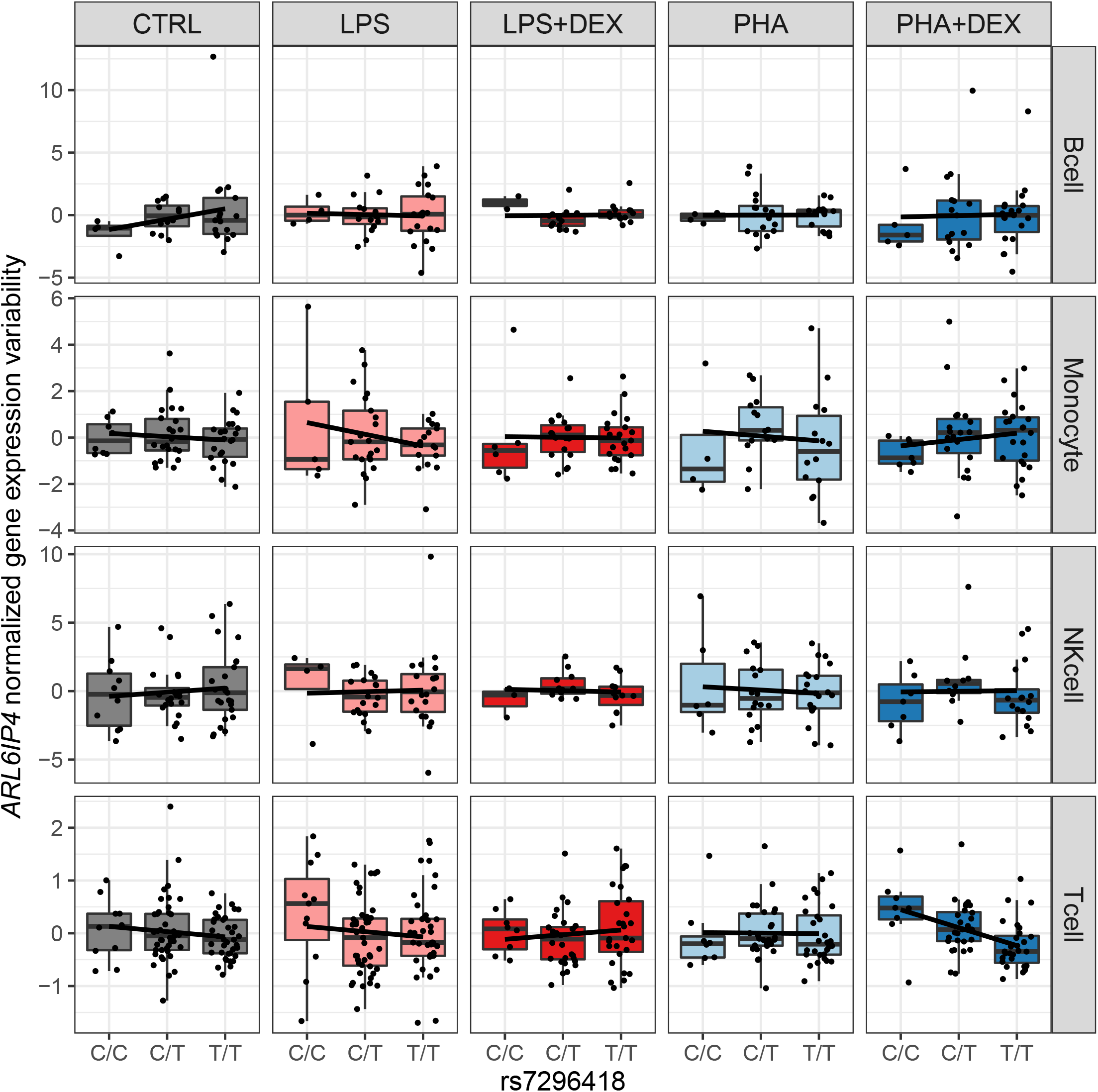
Genetic effect of rs7296418 on gene expression of ARL6IP4. **A** Boxplots represent normalized mean expression of the ARL6IP4 gene (y axis) across the three genotype classes of rs7296418 (x axis) across all conditions **B** Boxplots represent normalized gene expression variability of the ARL6IP4 gene (y axis) across the three genotype classes of rs7296418 (x axis) across all conditions.

**Figure S40:**
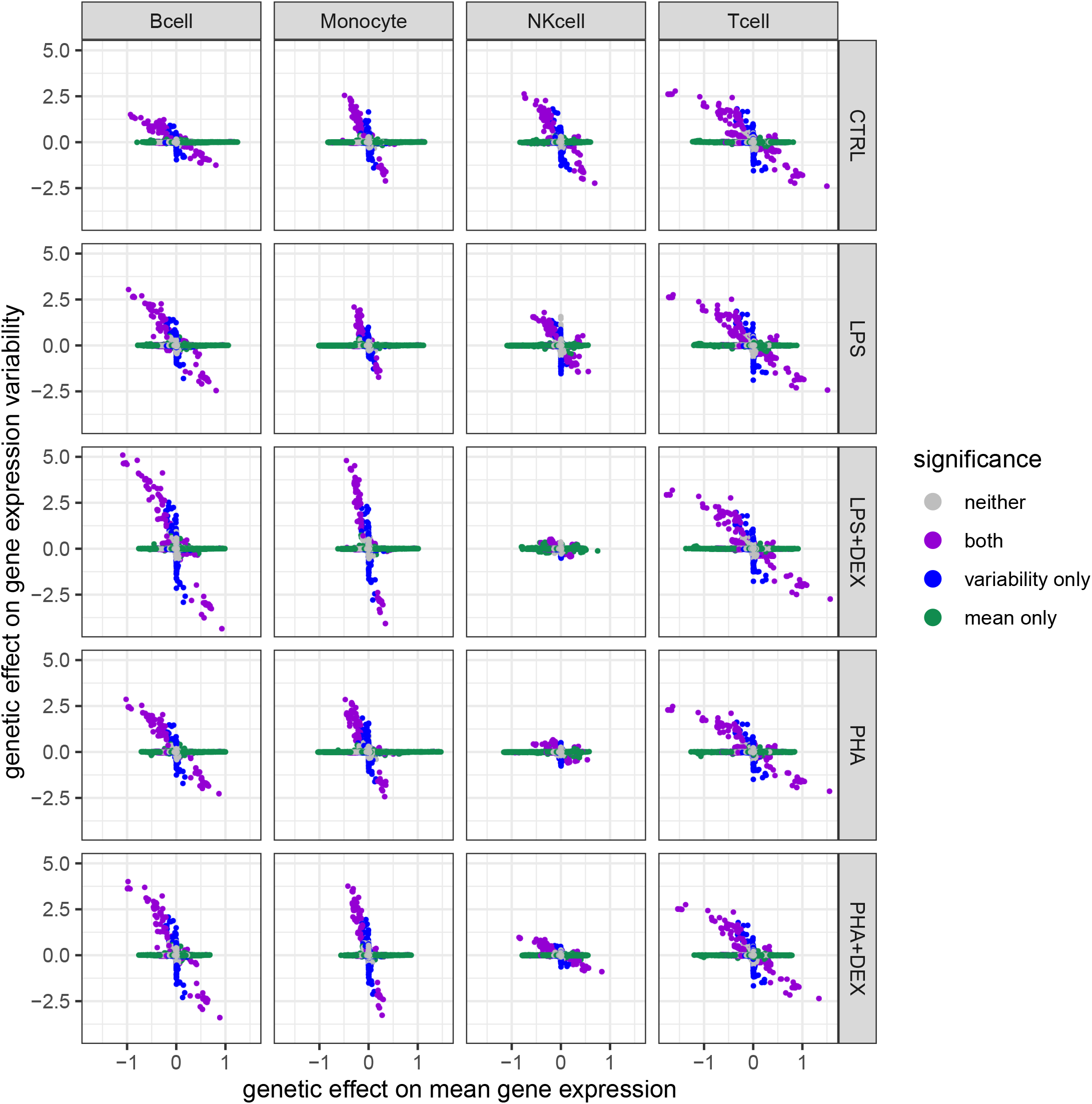
Genetic effects on gene expression variability. Scatterplots represent genetic effects on mean gene expression (x axis) versus genetic effects on gene expression variability (y axis) across all 20 conditions. Color represents genetic variants with significant effects on mean gene expression only (green), gene expression variability only (blue), both (purple), and neither (grey).

**Figure S41:**
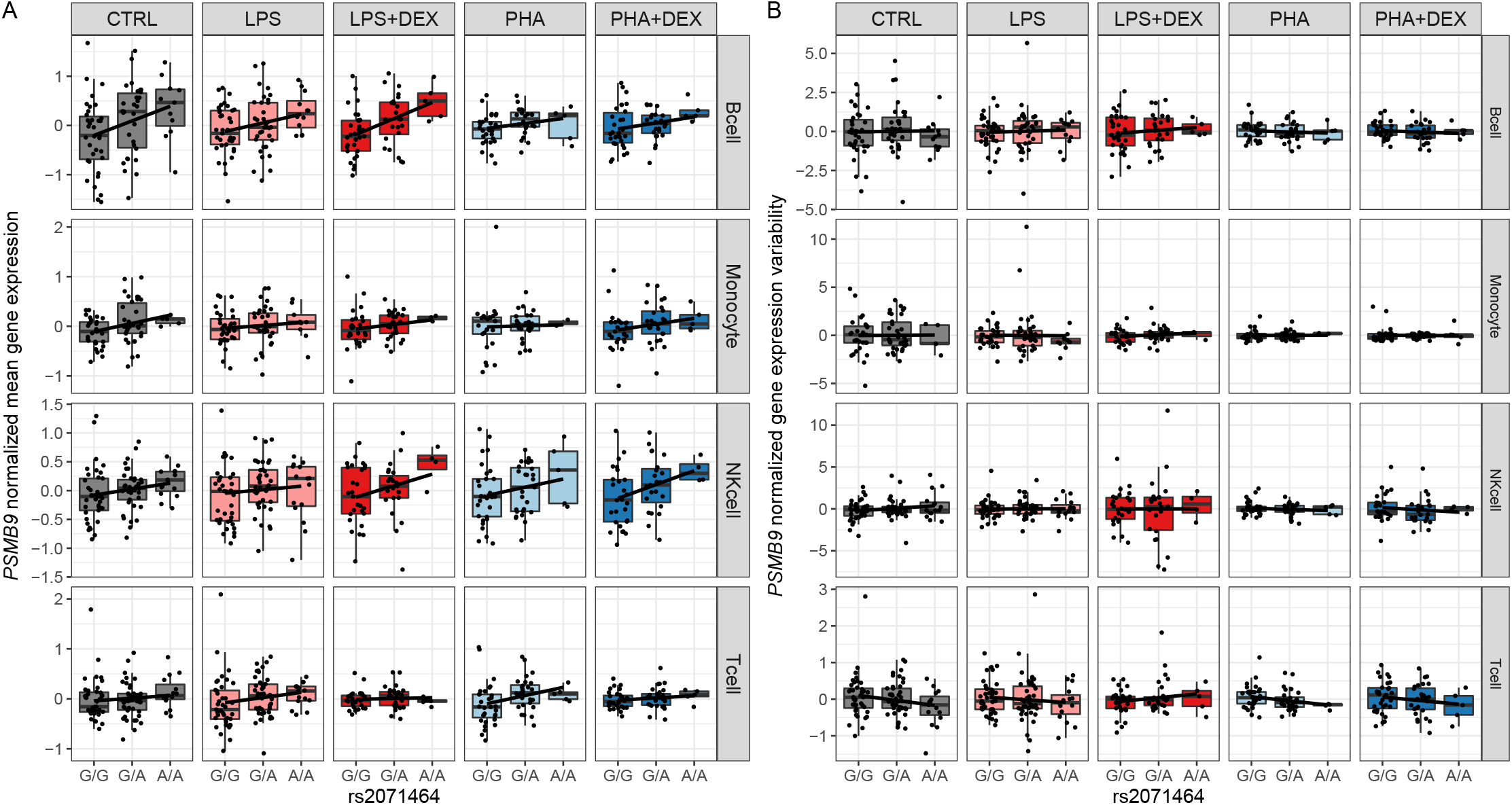
Genetic effect of rs2071464 on gene expression of PSMB9. **A** Boxplots represent normalized mean expression of the PSMB9 gene (y axis) across the three genotype classes of rs2071464 (x axis) across all conditions **B** Boxplots represent normalized gene expression variability of the PSMB9 gene (y axis) across the three genotype classes of rs2071464 (x axis) across all conditions.

**Figure S42:**
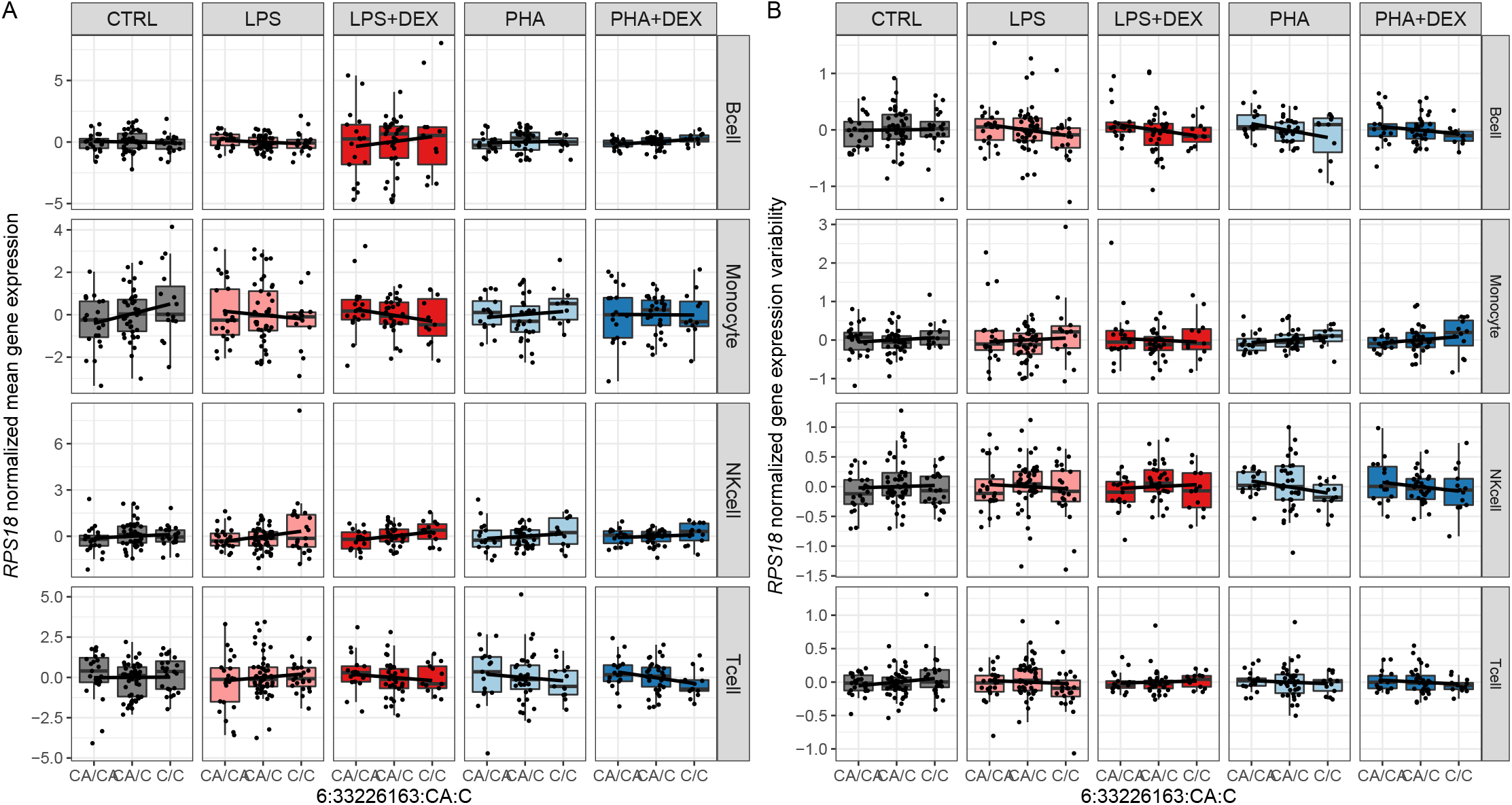
Genetic effect of the SNP at 6:33226163 on gene expression of RPS18. **A** Boxplots represent normalized mean expression of the RPS18 gene (y axis) across the three genotype classes of the SNP at 6:33226163 (x axis) across all conditions **B** Boxplots represent normalized gene expression variability of the RPS18 gene (y axis) across the three genotype classes of the SNP at 6:33226163 (x axis) across all conditions.

**Figure S43:**
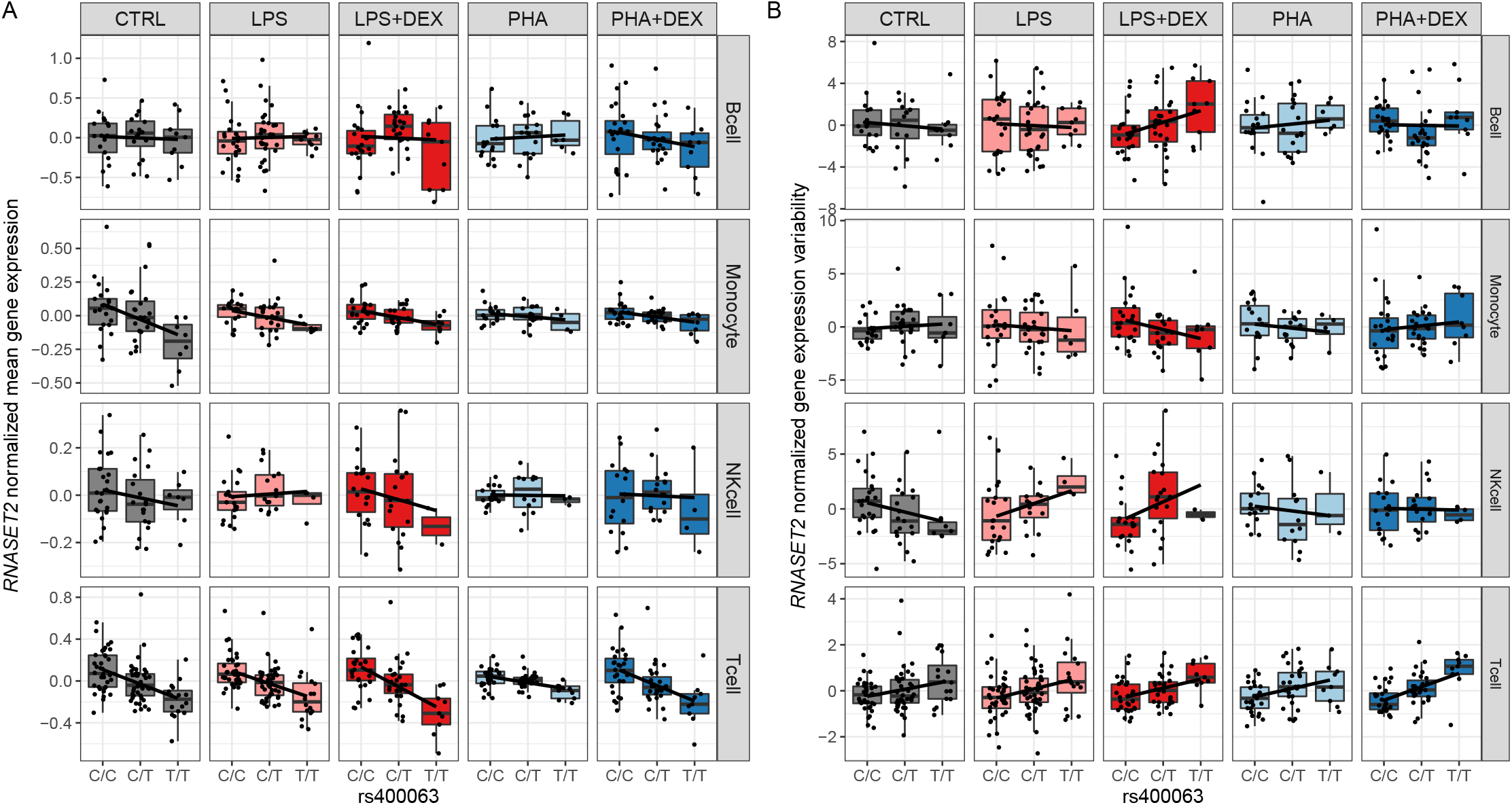
Genetic effect of rs12507413 on gene expression of RNASET2. **A** Boxplots represent normalized mean expression of the RNASET2 gene (y axis) across the three genotype classes of rs400063 (x axis) across all conditions **B** Boxplots represent normalized gene expression variability of the RNASET2 gene (y axis) across the three genotype classes of rsrs400063 (x axis) across all conditions.

**Figure S44:**
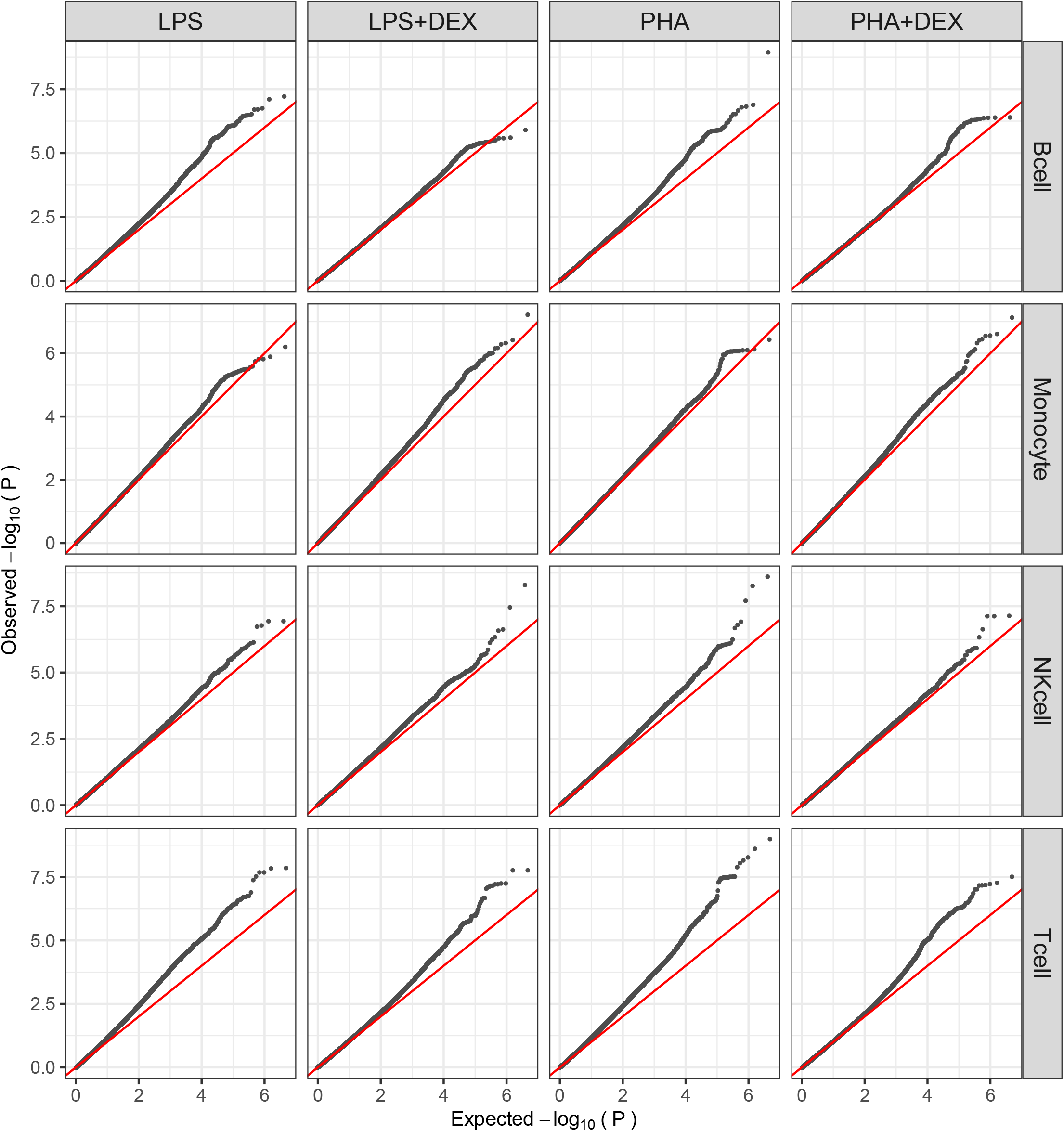
QQplots of p-values from DLDA-interacting eQTL mapping across the 16 conditions.

**Figure S45:**
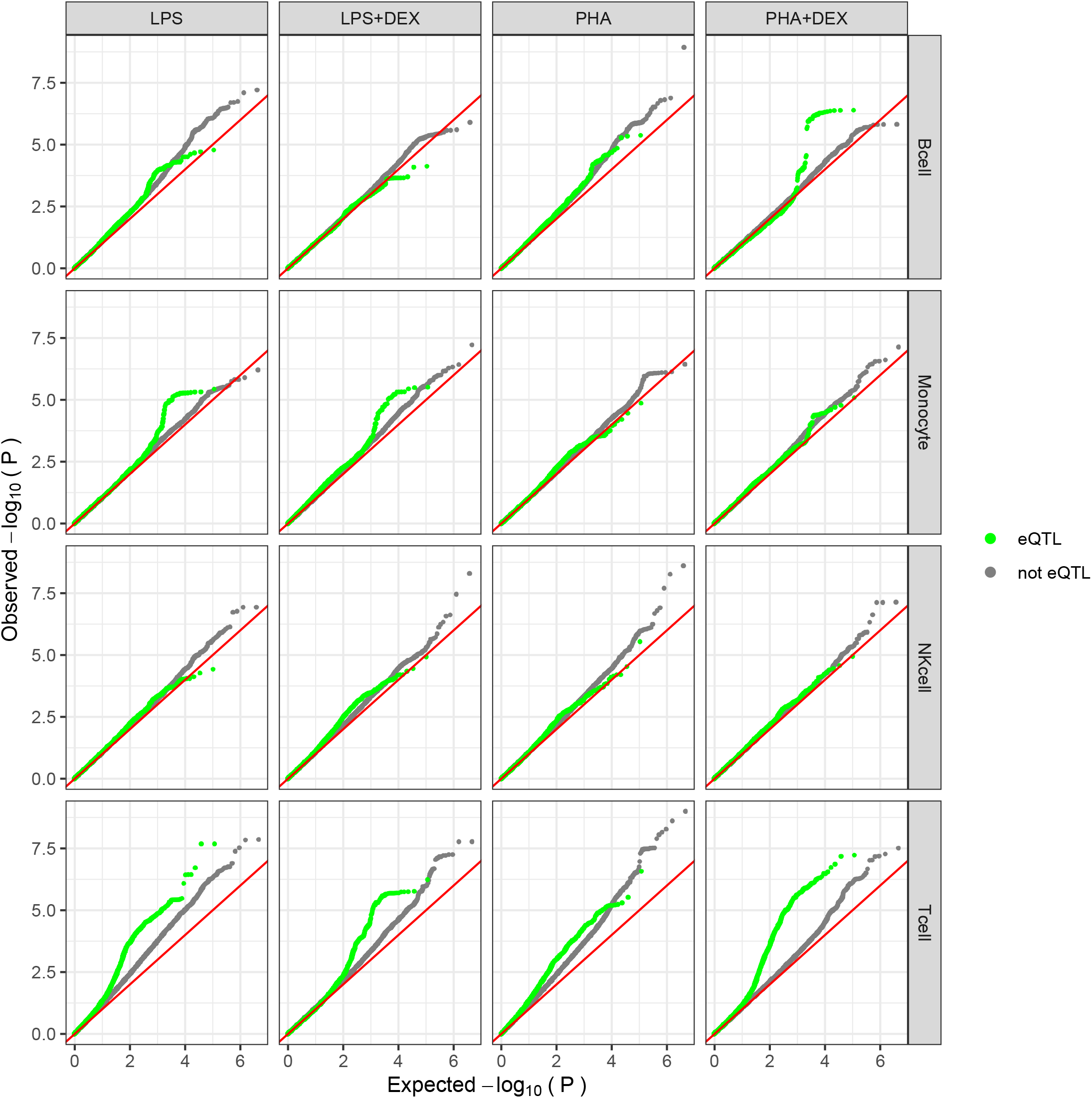
**Q-Qplots of p-values from DLDA-interacting eQTL mapping across the 16 conditions stratified by eQTL or not,** green colors representing the tested genetic variants are eQTLs that were detected in any condition.

**Figure S46:**
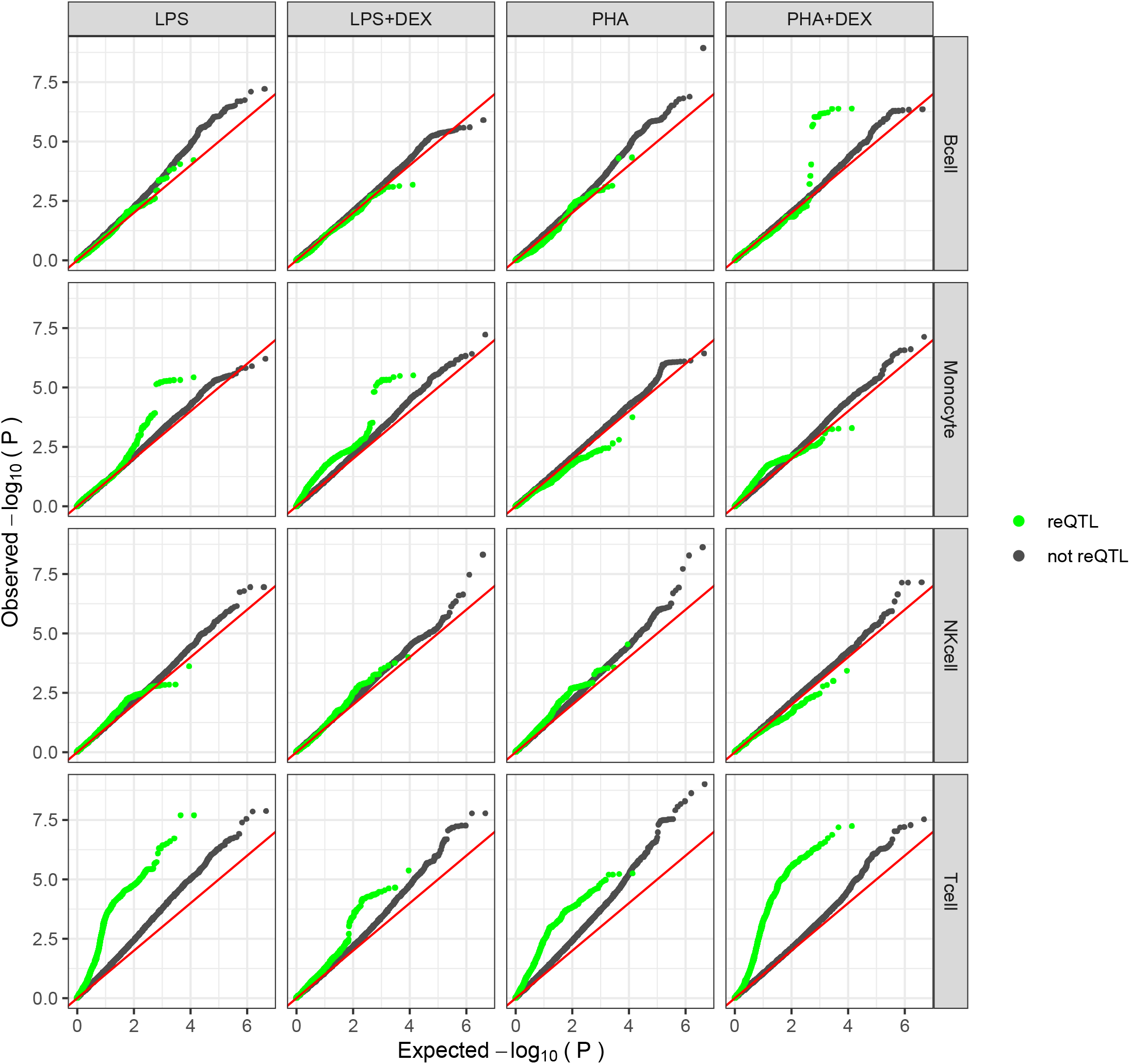
**Q-Qplots of p-values from DLDA-interacting eQTL mapping across the 16 conditions stratified by reQTL or not,** green colors representing the tested genetic variants are reQTLs that were detected in any condition.

**Figure S47:**
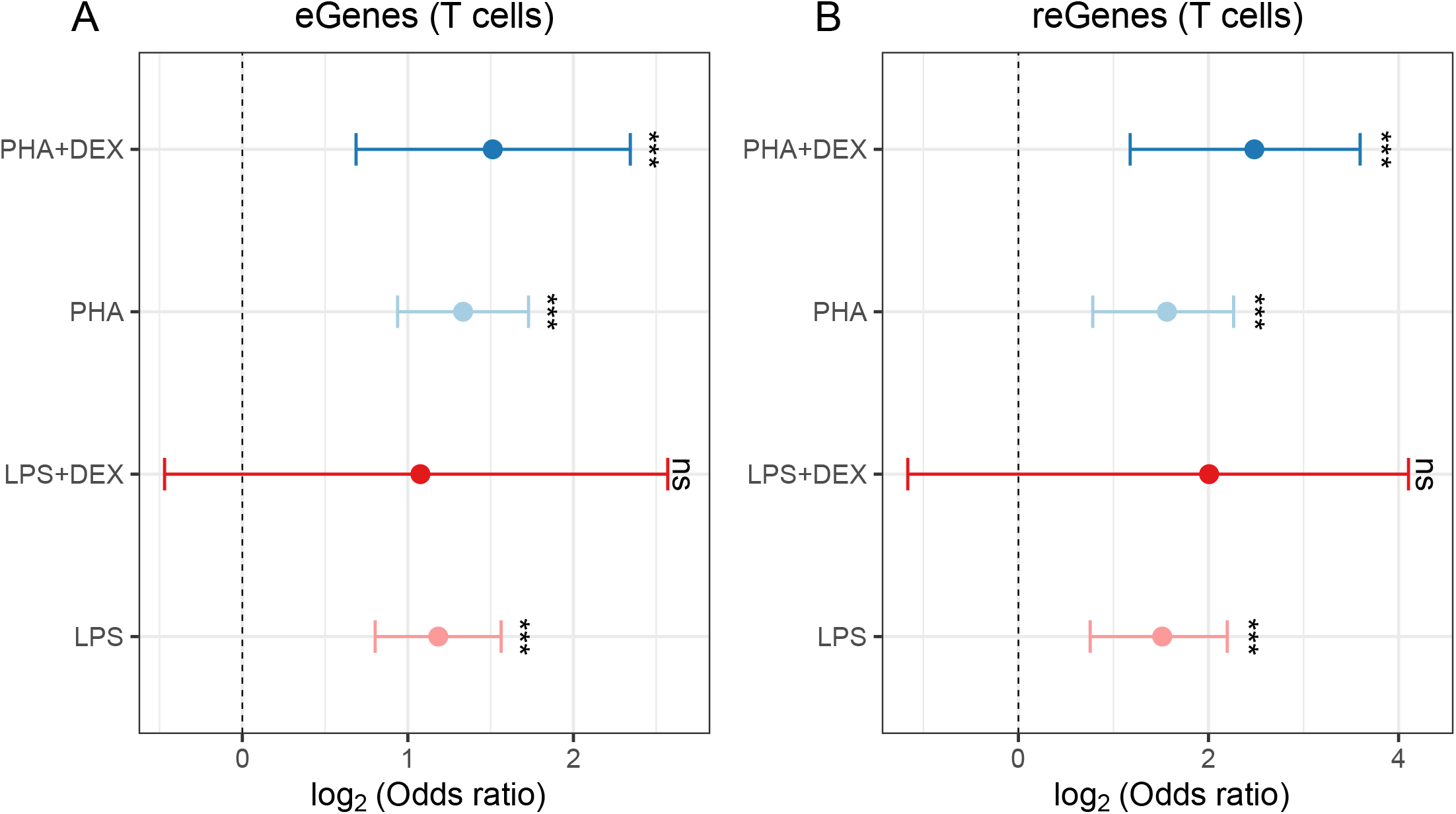
**Forest plots of log odds ratio across 4 conditions in the T cells**, to show if dynamic eGenes are significantly enriched in eGenes or reGenes across 4 conditions in the T cells. **(A)** test if dynamic eGenes are enriched in the eGenes discovered in the same condition, **(B)** test if dynamic eGenes are enriched in the reGenes discovered in the same condition

**Figure S48:**
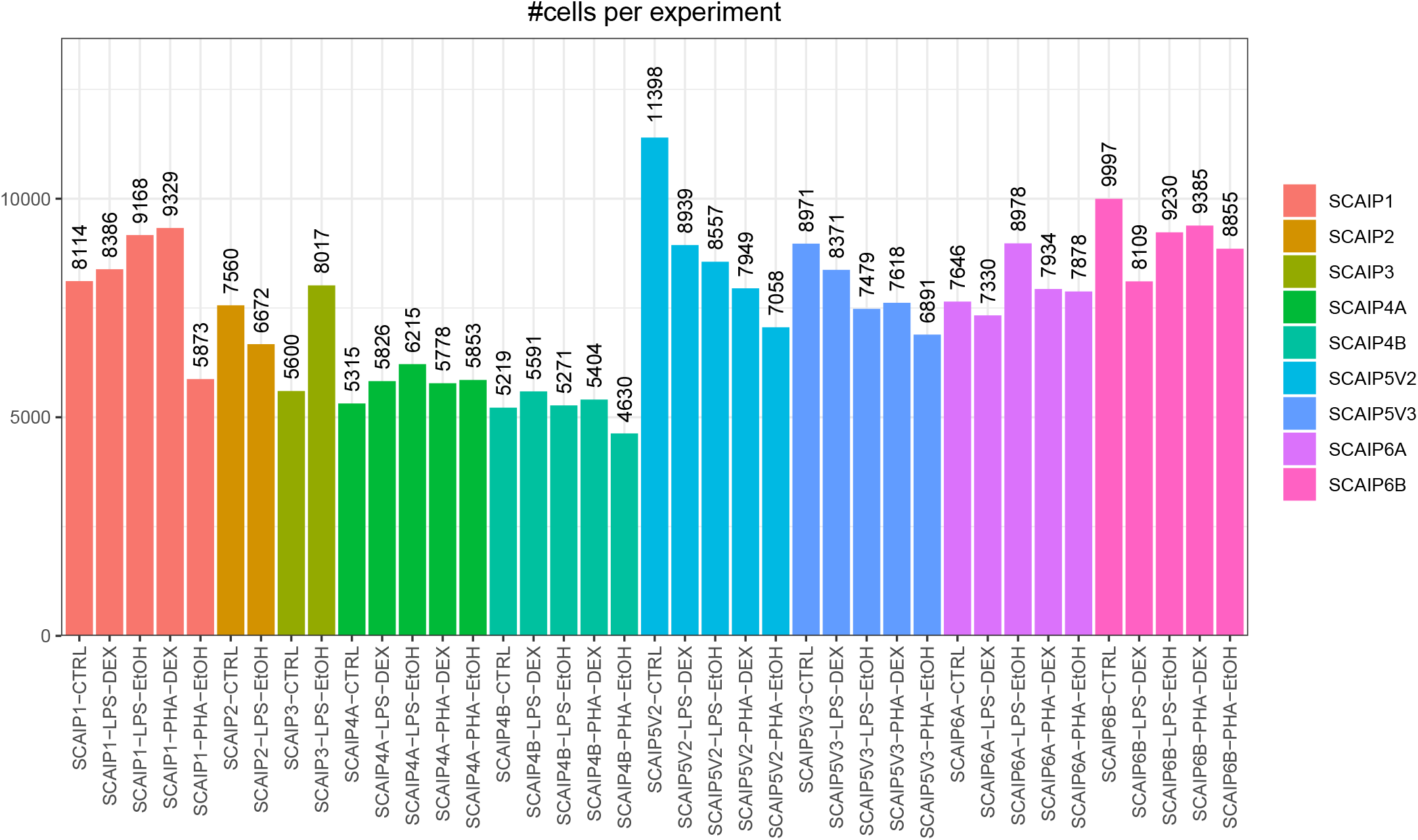
Number of measured cells across 39 experiments.

**Figure S49:**
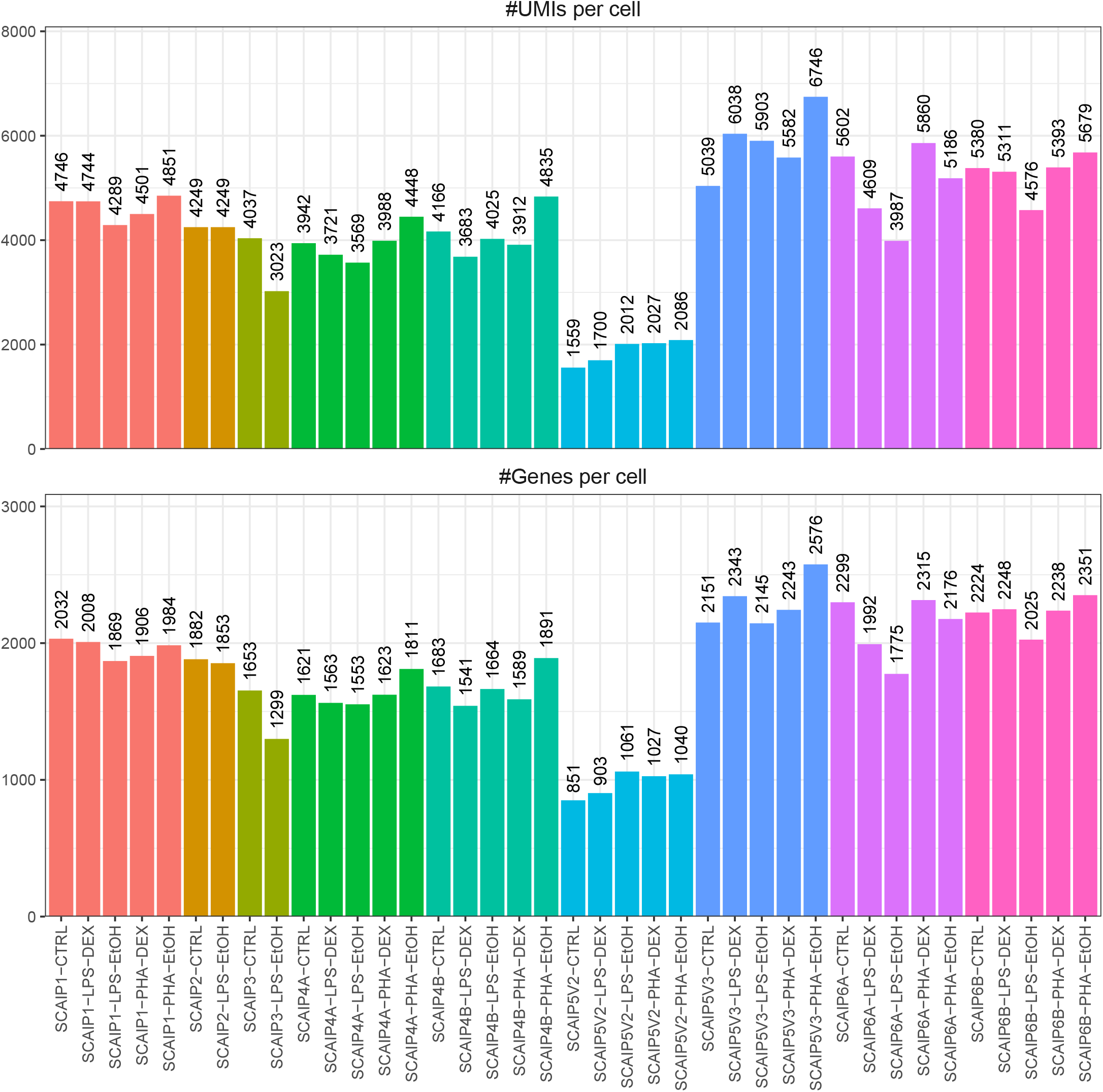
Unique molecular identifiers(UMIs) and gene features per cell across 39 experiments, including spliced and unspliced reads.

**Figure S50:**
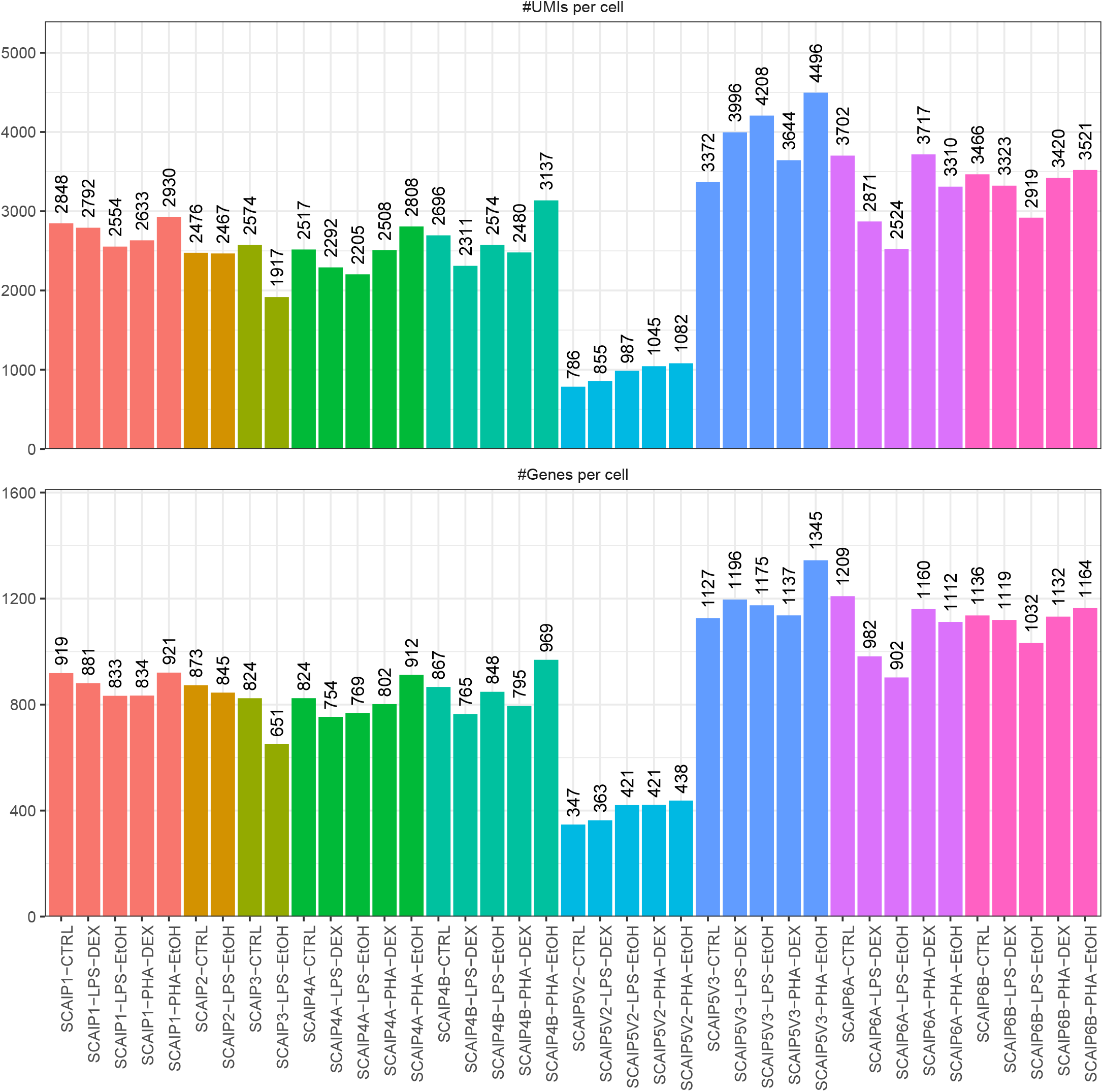
Unique molecular identifiers(UMIs) and gene features per cell across 39 experiments for spliced reads.

**Figure S51:**
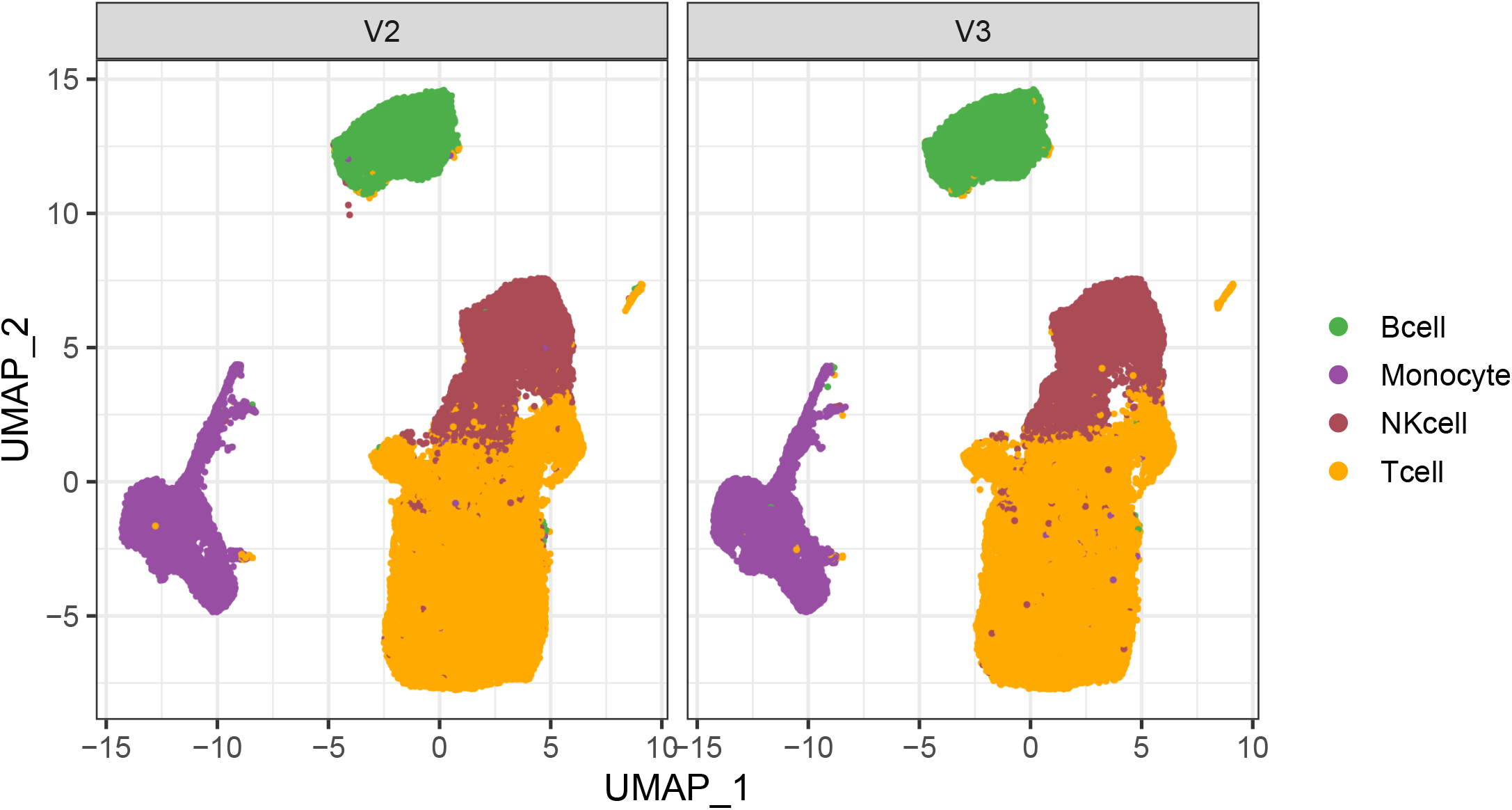
UMAP of cells split by chemistry.

**Figure S52:**
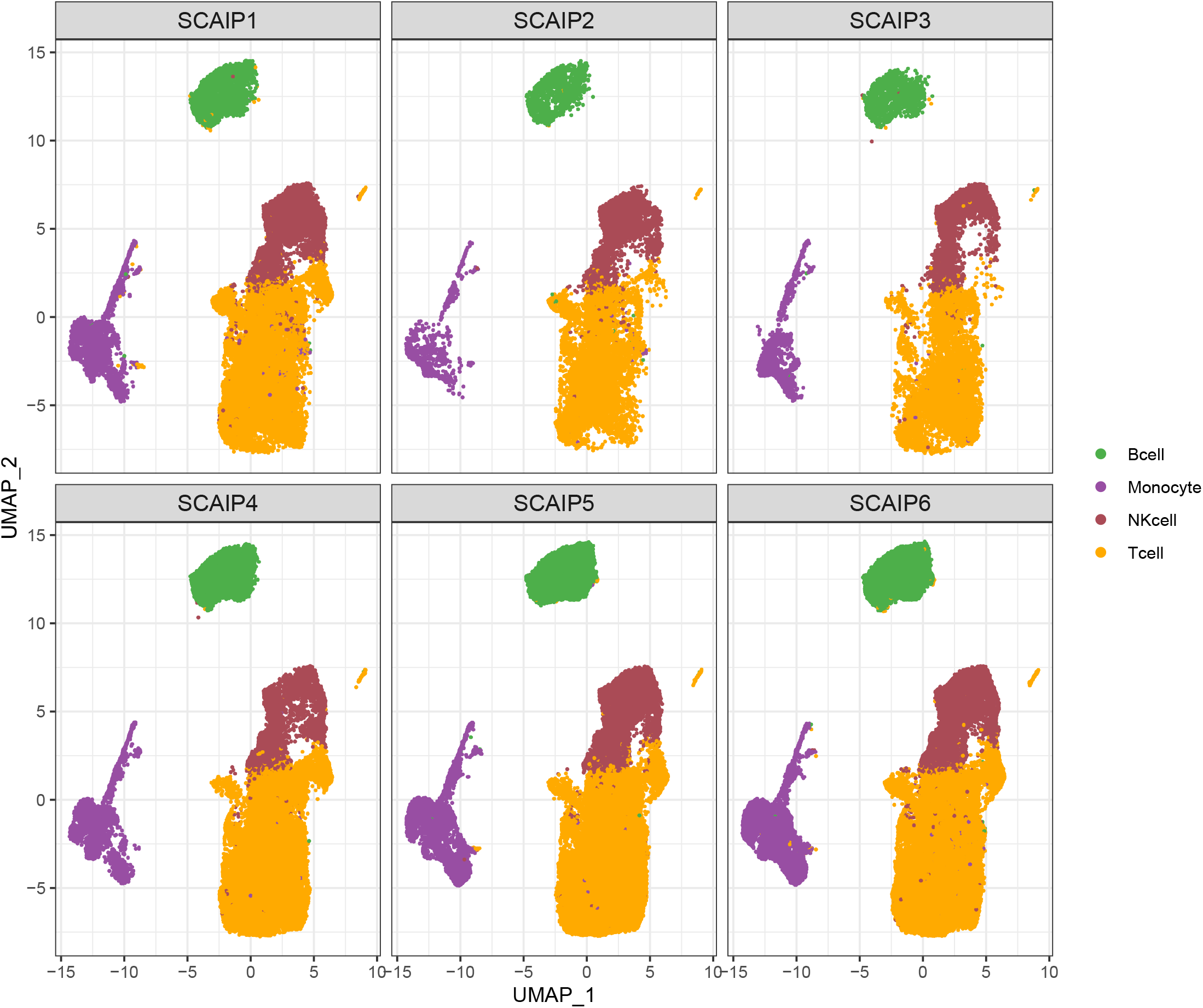
UMAP of cells split by BATCH.

**Figure S53:**
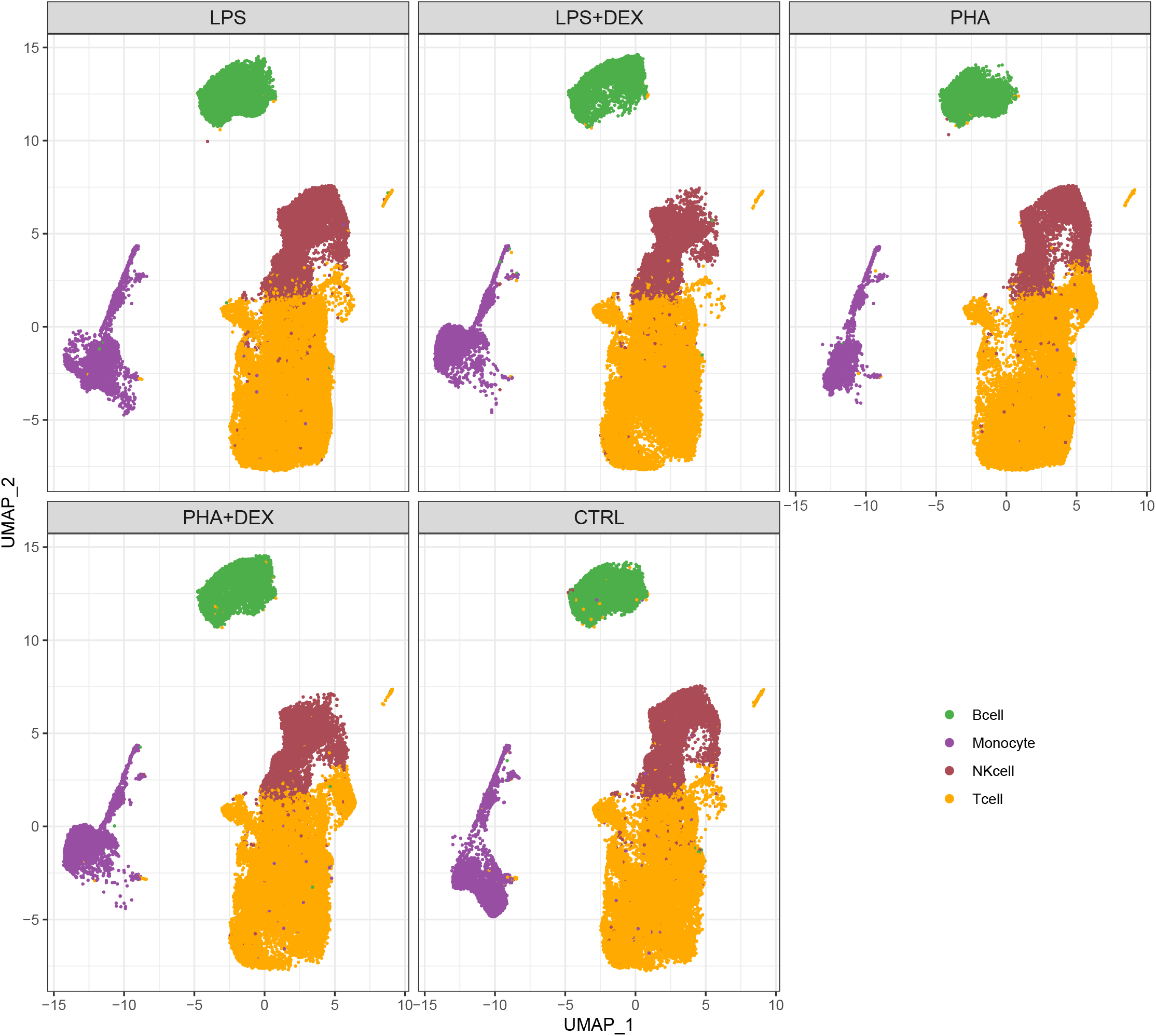
UMAP of cells split by treatment.

**Figure S54:**
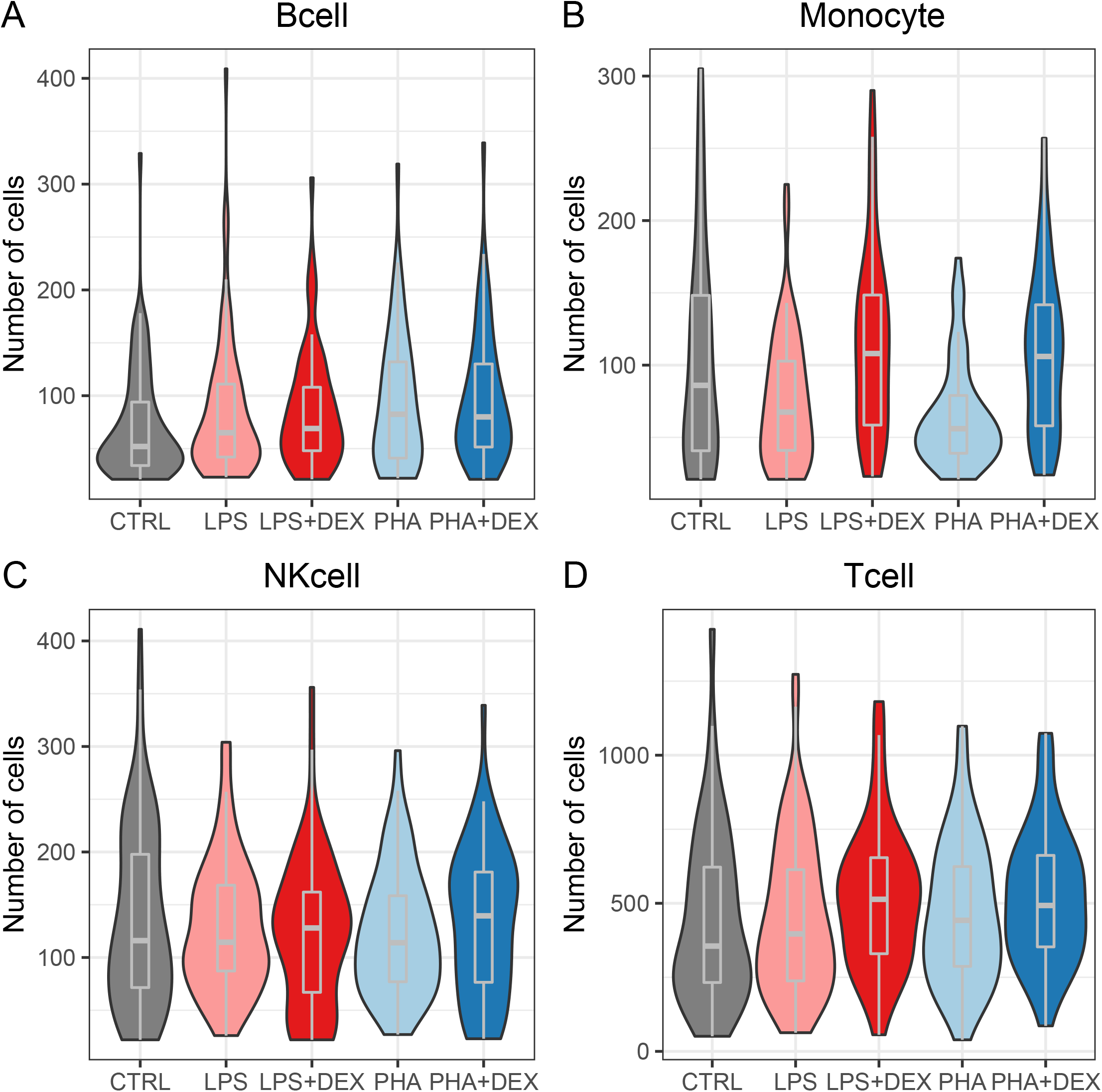
Violin plots of number of cells in each combination (cell-type+treatment+individual) **A-D** represents the distribution of the number of cells in each combination across individuals in different treatments for B cells, monocytes, NK cells and T cells.

## Supplementary Tables

**Table S1:** Summary statistics of single-cell data for each treatment. Column 2, number of individuals for each treatment; Column 3, median values of number of cells per individual across individuals in each treatment; Column 4, median values of average UMIs per cell from the same individual across individuals in each treatment, Column 5, median values of average detection genes per cell from the same individual across individuals in each treatment

|  | #Individuals | # Cells per indi-<br>vidual (median) | #UMIs per cell<br>(median) | #Genes per cell<br>(median) |
| --- | --- | --- | --- | --- |
| CTRL | 96 | 586 | 2,588 | 859 |
| LPS | 96 | 658 | 2,451 | 818 |
| LPS+DEX | 64 | 821 | 2,672 | 852 |
| PHA | 64 | 714 | 2,981 | 944 |
| PHA+DEX | 64 | 858 | 2,618 | 852 |

**Table S2:** Summary cell-type composition of individuals for each treatment. Column 2-5, number of cells from B-cells, Monocyte, NK-cell and T-cells of the individual for each treatment (median value). The full table detailing the number of cells in each cell-type/treatment/individual combination can be found in Table S23.

|  | #B-cell per individual<br>(median) | #Monocyte<br>per individual<br>(median) | #NK-cell<br>per individual<br>(median) | #T-cell<br>per individual<br>(median) |
| --- | --- | --- | --- | --- |
| CTRL | 52 | 86 | 116 | 356 |
| LPS-EtOH | 65 | 67.5 | 114 | 396 |
| LPS-DEX | 69 | 108 | 128 | 514 |
| PHA-EtOH | 82.5 | 56 | 114 | 442 |
| PHA-DEX | 80 | 106 | 140 | 492 |

**Table S3:** eQTL mapping results. Number of significant eQTLs and eGenes based on FastQTL results (FDR*<*0.1, columns 2 and 3), and multivariate adaptive shrinkage results (LFSR*<*0.1), columns 4 and 5.

|  | FastQTL FDR<0.1 |  | mash LFSR<0.1 |  |
| --- | --- | --- | --- | --- |
|  | eQTLs | eGenes | eQTLs | eGenes |
| Bcell CTRL | 5378 | 291 | 110276 | 2649 |
| Bcell LPS+DEX | 4465 | 262 | 109921 | 2656 |
| Bcell LPS | 8135 | 396 | 109869 | 2650 |
| Bcell PHA+DEX | 3671 | 217 | 113168 | 2652 |
| Bcell PHA | 4347 | 268 | 109853 | 2678 |
| Monocyte CTRL | 4186 | 253 | 106399 | 2603 |
| Monocyte LPS+DEX | 4720 | 300 | 103217 | 2559 |
| Monocyte LPS | 5518 | 340 | 101642 | 2586 |
| Monocyte PHA+DEX | 4019 | 271 | 104661 | 2547 |
| Monocyte PHA | 2407 | 167 | 101235 | 2590 |
| NKcell CTRL | 5986 | 364 | 113162 | 2624 |
| NKcell LPS+DEX | 1566 | 122 | 103316 | 2592 |
| NKcell LPS | 5379 | 366 | 110434 | 2462 |
| NKcell PHA+DEX | 1427 | 97 | 104507 | 2602 |
| NKcell PHA | 1621 | 135 | 110859 | 2477 |
| Tcell CTRL | 20187 | 1297 | 109130 | 2605 |
| Tcell LPS+DEX | 15636 | 1052 | 114176 | 2642 |
| Tcell LPS | 22603 | 1386 | 112189 | 2684 |
| Tcell PHA+DEX | 14568 | 967 | 115410 | 2666 |
| Tcell PHA | 13279 | 884 | 112915 | 2712 |
| Total Unique | 62694 | 5190 | 129539 | 2984 |
| Total Tested | 4163269 | 13876 | 2,095,831 | 9556 |

**Table S4:** Overlap of eGenes with differentially expressed genes (DEGs, FDR<0.1) and differentially variable genes (DVGs, FDR<0.1). Columns represent: 1 – condition, 2 – number of eGenes which are also DEGs for the condition against its paired control, 3 – odds ratio of the overlap in column 2, 4 – p-value of the Fisher’s exact test for enrichment of eGenes in DEGs, 5 - number of eGenes which are also DVGs for the condition against its paired control, 6 – odds ratio of the overlap in column 5, 7 – p-value of the Fisher’s exact test for enrichment of eGenes in DVGs.

| Condition | eGenes that are DEGs |  |  | eGenes that are DVGs |  |  |
| --- | --- | --- | --- | --- | --- | --- |
|  | Overlap | OR | p-value | overlap | OR | p-value |
| Bcell LPS | 111 | 2.19 | 5.1E-09 | 8 | 1.25 | 8.0E-01 |
| Bcell LPS+DEX | 297 | 1.61 | 1.5E-09 | 33 | 1.90 | 3.2E-02 |
| Bcell PHA | 406 | 1.46 | 1.8E-08 | 52 | 3.27 | 3.2E-06 |
| Bcell PHA+DEX | 355 | 1.58 | 2.2E-10 | 41 | 1.98 | 6.3E-03 |
| Monocyte LPS | 382 | 1.15 | 3.7E-02 | 199 | 2.33 | 2.1E-13 |
| Monocyte LPS+DEX | 368 | 1.19 | 1.2E-02 | 131 | 2.63 | 3.2E-11 |
| Monocyte PHA | 660 | 1.20 | 6.4E-04 | 292 | 2.78 | 3.5E-25 |
| Monocyte PHA+DEX | 400 | 1.27 | 2.9E-04 | 178 | 2.00 | 3.4E-09 |
| NKcell LPS | 104 | 1.99 | 3.0E-07 | 27 | 1.73 | 9.4E-02 |
| NKcell LPS+DEX | 410 | 1.61 | 1.2E-12 | 54 | 2.02 | 4.0E-03 |
| NKcell PHA | 376 | 1.54 | 7.4E-10 | 49 | 3.33 | 8.1E-06 |
| NKcell PHA+DEX | 493 | 1.60 | 4.2E-14 | 36 | 2.24 | 5.5E-03 |
| Tcell LPS | 79 | 1.98 | 9.1E-06 | 57 | 3.22 | 4.4E-08 |
| Tcell LPS+DEX | 347 | 1.43 | 5.5E-07 | 66 | 2.58 | 7.9E-07 |
| Tcell PHA | 282 | 1.48 | 7.9E-07 | 89 | 3.88 | 2.2E-14 |
| Tcell PHA+DEX | 408 | 1.50 | 1.7E-09 | 100 | 2.31 | 3.8E-08 |

**Table S5:** Overlap of reGenes with differentially expressed genes (DEGs, FDR*<*0.1) and differentially variable genes (DVGs, FDR*<*0.1). Columns represent: 1 – condition, 2 – number of reGenes which are also DEGs for the condition against its paired control, 3 – odds ratio of the overlap in column 2, 4 – p-value of the Fisher’s exact test for enrichment of reGenes in DEGs, 5 - number of reGenes which are also DVGs for the condition against its paired control, 6 – odds ratio of the overlap in column 5, 7 – p-value of the Fisher’s exact test for enrichment of reGenes in DVGs.

| Condition | reGenes that are DEGs |  |  | reGenes that are DVGs |  |  |
| --- | --- | --- | --- | --- | --- | --- |
|  | Overlap | OR | p-value | overlap | OR | p-value |
| Bcell LPS | 2 | 1.95 | 2.8E-01 | 1 | 14.80 | 7.3E-02 |
| Bcell LPS+DEX | 11 | 3.12 | 2.4E-03 | 3 | 7.44 | 1.1E-02 |
| Bcell PHA+DEX | 10 | 1.76 | 1.3E-01 | 2 | 3.33 | 1.3E-01 |
| Bcell PHA | 3 | 0.72 | 7.9E-01 | 1 | 3.51 | 2.6E-01 |
| Monocyte LPS | 14 | 1.28 | 4.2E-01 | 8 | 2.21 | 5.8E-02 |
| Monocyte LPS+DEX | 5 | 0.57 | 2.7E-01 | 2 | 1.10 | 7.1E-01 |
| Monocyte PHA+DEX | 11 | 1.56 | 1.7E-01 | 7 | 3.08 | 1.4E-02 |
| Monocyte PHA | 16 | 1.01 | 1.0E+00 | 6 | 1.39 | 4.5E-01 |
| NKcell LPS | 3 | 3.56 | 6.1E-02 | 0 | NA | NA |
| NKcell LPS+DEX | 13 | 1.84 | 5.6E-02 | 1 | 0.97 | 1.0E+00 |
| NKcell PHA | 3 | 1.09 | 7.5E-01 | 0 | NA | NA |
| NKcell PHA+DEX | 12 | 1.44 | 2.7E-01 | 4 | 6.14 | 6.2E-03 |
| Tcell LPS | 0 | NA | NA | 1 | 2.07 | 3.9E-01 |
| Tcell LPS+DEX | 7 | 1.21 | 6.6E-01 | 0 | NA | NA |
| Tcell PHA+DEX | 6 | 1.09 | 8.2E-01 | 1 | 0.94 | 1.0E+00 |
| Tcell PHA | 3 | 0.61 | 6.3E-01 | 3 | 3.32 | 7.0E-02 |

**Table S6:** vQTL mapping results. Number of significant vQTLs and vGenes based on FastQTL results (FDR<0.1, columns 2 and 3), and multivariate adaptive shrinkage results (LFSR<0.1), columns 4 and 5.

|  | FastQTL FDR<0.1 |  | mash LFSR<0.1 |  |
| --- | --- | --- | --- | --- |
|  | vQTLs | vGenes | vQTLs | vGenes |
| Bcell CTRL | 37 | 6 | 3086 | 96 |
| Bcell LPS+DEX | 9 | 4 | 3042 | 98 |
| Bcell LPS | 0 | 0 | 2638 | 104 |
| Bcell PHA+DEX | 2 | 1 | 2728 | 100 |
| Bcell PHA | 0 | 0 | 2866 | 99 |
| Monocyte CTRL | 2 | 2 | 2816 | 101 |
| Monocyte LPS+DEX | 86 | 12 | 2827 | 96 |
| Monocyte LPS | 67 | 12 | 2573 | 100 |
| Monocyte PHA+DEX | 20 | 7 | 2937 | 100 |
| Monocyte PHA | 50 | 9 | 2856 | 103 |
| NKcell CTRL | 0 | 0 | 3211 | 99 |
| NKcell LPS+DEX | 0 | 0 | 2632 | 91 |
| NKcell LPS | 16 | 7 | 3010 | 82 |
| NKcell PHA+DEX | 0 | 0 | 2941 | 87 |
| NKcell PHA | 19 | 3 | 2819 | 90 |
| Tcell CTRL | 188 | 24 | 3052 | 106 |
| Tcell LPS+DEX | 23 | 4 | 3230 | 98 |
| Tcell LPS | 154 | 22 | 2989 | 107 |
| Tcell PHA+DEX | 17 | 3 | 3151 | 106 |
| Tcell PHA | 0 | 0 | 3089 | 101 |
| Total unique | 545 | 102 | 3563 | 123 |
| Total tested | 1549665 | 7055 | 242,494 | 1086 |

**Table S7:** Mean-eQTL mapping results. Number of significant mean-eQTLs and mean-eGenes based on FastQTL results (FDR*<*0.1, columns 2 and 3), and multivariate adaptive shrinkage results (LFSR*<*0.1), columns 4 and 5.

|  | FastQTL FDR<0.1 |  | mash LFSR<0.1 |  |
| --- | --- | --- | --- | --- |
|  | mean-eQTLs | mean-eGenes | mean-eQTLs | mean-eGenes |
| Bcell CTRL | 604 | 50 | 9420 | 222 |
| Bcell LPS+DEX | 1068 | 73 | 10252 | 238 |
| Bcell LPS | 4075 | 225 | 10882 | 232 |
| Bcell PHA+DEX | 812 | 54 | 11098 | 228 |
| Bcell PHA | 2018 | 125 | 10313 | 249 |
| Monocyte CTRL | 56 | 12 | 8616 | 203 |
| Monocyte LPS+DEX | 1 | 1 | 7995 | 187 |
| Monocyte LPS | 401 | 39 | 7925 | 187 |
| Monocyte PHA+DEX | 410 | 59 | 8294 | 200 |
| Monocyte PHA | 15 | 1 | 8239 | 186 |
| NKcell CTRL | 783 | 71 | 10929 | 245 |
| NKcell LPS+DEX | 104 | 14 | 11250 | 214 |
| NKcell LPS | 1089 | 94 | 8635 | 254 |
| NKcell PHA+DEX | 617 | 64 | 11446 | 202 |
| NKcell PHA | 702 | 56 | 7891 | 256 |
| Tcell CTRL | 11912 | 792 | 12404 | 285 |
| Tcell LPS+DEX | 9226 | 653 | 12463 | 286 |
| Tcell LPS | 11744 | 781 | 12465 | 302 |
| Tcell PHA+DEX | 7515 | 562 | 12156 | 288 |
| Tcell PHA | 6477 | 506 | 12441 | 287 |
| Total unique | 30493 | 2430 | 13164 | 366 |
| Total tested | 1549634 | 7055 | 241568 | 1084 |

**Table S8:** Overlap of vGenes with differentially expressed genes (DEGs, FDR<0.1) and differentially variable genes (DVGs, FDR<0.1). Columns represent: 1 – condition, 2 – number of vGenes which are also DEGs for the condition against its paired control, 3 – odds ratio of the overlap in column 2, 4 – p-value of the Fisher’s exact test for enrichment of vGenes in DEGs, 5 - number of vGenes which are also DVGs for the condition against its paired control, 6 – odds ratio of the overlap in column 5, 7 – p-value of the Fisher’s exact test for enrichment of vGenes in DVGs.

| Condition | vGenes that are DEGs |  |  | vGenes that are DVGs |  |  |
| --- | --- | --- | --- | --- | --- | --- |
|  | Overlap | OR | p-value | overlap | OR | p-value |
| Bcell LPS | 7 | 6.81 | 5.5E-04 | 4 | 6.94 | 8.6E-03 |
| Bcell LPS+DEX | 10 | 2.27 | 3.1E-02 | 3 | 1.59 | 4.4E-01 |
| Bcell PHA | 18 | 2.70 | 1.1E-03 | 15 | 6.49 | 8.9E-07 |
| Bcell PHA+DEX | 12 | 1.81 | 7.5E-02 | 6 | 2.48 | 5.5E-02 |
| Monocyte LPS | 5 | 0.45 | 1.1E-01 | 7 | 0.89 | 1.0E+00 |
| Monocyte LPS+DEX | 3 | 0.45 | 2.7E-01 | 7 | 1.32 | 4.9E-01 |
| Monocyte PHA | 14 | 0.75 | 4.1E-01 | 16 | 1.28 | 4.4E-01 |
| Monocyte PHA+DEX | 9 | 1.25 | 5.5E-01 | 16 | 2.55 | 3.0E-03 |
| NKcell LPS | 5 | 2.35 | 8.8E-02 | 7 | 5.08 | 1.9E-03 |
| NKcell LPS+DEX | 9 | 1.11 | 7.0E-01 | 11 | 5.38 | 6.9E-05 |
| NKcell PHA | 8 | 1.21 | 5.4E-01 | 6 | 2.17 | 1.2E-01 |
| NKcell PHA+DEX | 13 | 1.21 | 5.1E-01 | 2 | 1.17 | 6.9E-01 |
| Tcell LPS | 6 | 4.53 | 6.3E-03 | 11 | 3.65 | 1.1E-03 |
| Tcell LPS+DEX | 13 | 1.69 | 1.3E-01 | 6 | 1.88 | 1.6E-01 |
| Tcell PHA | 16 | 3.04 | 6.2E-04 | 20 | 4.51 | 1.8E-06 |
| Tcell PHA+DEX | 18 | 2.13 | 1.3E-02 | 7 | 0.97 | 1.0E+00 |

**Table S9:** Summary of DLDA dynamic eQTL mapping and overlap with eGenes, and and immune diseases-associated genes identified from PTWAS

| Cell-type DLDA treatment | eQTLs(FDR<0.1) | eGenes | PTWAS Genes |
| --- | --- | --- | --- |
| Bcell LPS | 139 | 32 | 2 |
| Bcel LPS+DEX | 0 | 0 | 0 |
| Bcell PHA | 114 | 40 | 7 |
| Bcell PHA+DEX | 27 | 2 | 0 |
| Monocyte LPS | 35 | 3 | 0 |
| Monocyte LPS+DEX | 37 | 3 | 1 |
| Monocyte PHA | 0 | 0 | 0 |
| Monocyte PHA+DEX | 0 | 0 | 0 |
| NKcell LPS | 4 | 3 | 0 |
| NKcell LPS+DEX | 2 | 2 | 0 |
| NKcell PHA | 6 | 4 | 1 |
| NKcell PHA+DEX | 3 | 2 | 0 |
| Tcell LPS | 2073 | 244 | 45 |
| Tcell LPS+DEX | 269 | 18 | 5 |
| Tcell PHA | 1329 | 225 | 39 |
| Tcel PHA+DEX | 1298 | 54 | 9 |
| Total unique | 3899 | 588 | 101 |

**Table S10: Results of differential expressed genes (DEGs) using DESeq2 pseudo bulk aggregated data**. Columns 1-7 are: 1) Cell type; 2) Treatment; 3) Ensembl gene ID; 4) Log2 fold change of gene expression; 5) Standard error; 6) P-value; 7) FDR

http://genome.grid.wayne.edu/SCAIP/Supp_Tables/Differential_expression.txt.gz

**Table S11: Results of differential gene expression variability (DGV)**. Columns 1-7 are: 1) Cell type; 2) Treatment; 3) Ensembl gene ID; 4) Log2 fold change of gene variability; 5) Standard error; 6) P-value; 7) FDR

http://genome.grid.wayne.edu/SCAIP/Supp_Tables/Differential_variability.txt.gz

**Table S12: FastQTL eQTL mapping results**. Results of eQTL mapping across the 20 conditions. Columns are: 1 - ENSEMBL gene name; 2 - genetic variant coordinates (GRCh37); major and minor alleles and dbSNP ID; 3 - genetic variant distance from the TSS of the gene; 4 - p-value of the effect of the genetic variant on expression of the gene; 5 - effect size of the genetic variant on expression of the gene; 6 - condition.

http://genome.grid.wayne.edu/SCAIP/Supp_Tables/FastQTL-pseudobulk-4PC.txt.gz

**Table S13: mash eQTL effect estimates**. Multivariate adaptive shrinkage estimates of genetic effects on gene expression. Columns 1-20 are the conditions; rows are the tested gene-SNP pairs.

http://genome.grid.wayne.edu/SCAIP/Supp_Tables/effect_mash_pseudobulk.txt.gz

**Table S14: mash eQTL significance**. Multivariate adaptive shrinkage significance (LFSR) of genetic effects on gene expression. Columns 1-20 are the conditions; rows are the tested gene-SNP pairs.

http://genome.grid.wayne.edu/SCAIP/Supp_Tables/lfsr_mash_pseudobulk.txt.gz

**Table S15:** 61 reGenes overlapping with immune disease genee. Columns are: 1-ENSEMBL gene name; 2-SYMBOL gene name

http://genome.grid.wayne.edu/SCAIP/Supp_Tables/reGene_twas.txt.gz

**Table S16: FastQTL vQTL mapping results**. Results of vQTL mapping across the 20 conditions. Columns are: 1 - ENSEMBL gene name; 2 - genetic variant coordinates (GRCh37), major and minor alleles and dbSNP ID; 3 - genetic variant distance from the TSS of the gene; 4 - p-value of the effect of the genetic variant on gene expression variability of the gene; 5 - effect size of the genetic variant on gene expression variability of the gene; 6 - condition.

http://genome.grid.wayne.edu/SCAIP/Supp_Tables/FastQTL-var-5PC.txt.gz

**Table S17: mash vQTL effect estimates**. Multivariate adaptive shrinkage estimates of genetic effects on gene expression variability. Columns 1-20 are the conditions, rows are the tested gene-SNP pairs.

http://genome.grid.wayne.edu/SCAIP/Supp_Tables/effect_mash_var.txt.gz

**Table S18: mash vQTL significance**. Multivariate adaptive shrinkage significance of genetic effects on gene expression variability. Columns 1-20 are the conditions; rows are the tested gene-SNP pairs.

http://genome.grid.wayne.edu/SCAIP/Supp_Tables/lfsr_mash_var.txt.gz

**Table S19: FastQTL mean-eQTL mapping results**. Results of mean-eQTL mapping across the 20 conditions. Columns are: 1 - ENSEMBL gene name; 2 - genetic variant coordinates (GRCh37), major and minor alleles and dbSNP ID; 3 - genetic variant distance from the TSS of the gene; 4 - p-value of the effect of the genetic variant on mean expression of the gene; 5 - effect size of the genetic variant on mean expression of the gene; 6 - condition.

http://genome.grid.wayne.edu/SCAIP/Supp_Tables/FastQTL-mean-7PC.txt.gz

**Table S20: mash mean-eQTL effect estimates**. Multivariate adaptive shrinkage estimates of genetic effects on gene expression mean. Columns 1-20 are the conditions, rows are the tested gene-SNP pairs.

http://genome.grid.wayne.edu/SCAIP/Supp_Tables/effect_mash_mean.txt.gz

**Table S21: mash mean-eQTL significance**. Multivariate adaptive shrinkage significance of genetic effects on gene expression mean. Columns 1-20 are the conditions, rows are the tested gene-SNP pairs.

http://genome.grid.wayne.edu/SCAIP/Supp_Tables/lfsr_mash_mean.txt.gz

**Table S22: Dynamic eQTL mapping results**. Results of dynamic eQTL mapping across 16 conditions, each file representing the results of dynamic eQTL mapping in one of the four cell types (B cells, Monocytes, NK cells and T cells) treated with LPS, LPS+DEX, PHA or PHA+DEX interacting with treatment response pseudotime. Columns 1-6 are: 1) Ensembl gene ID; 2) genetic variant coordinates (GRCh37); major and minor alleles and dbSNP ID; 3) genetic effect on gene expression interacting with response pseudotime; 4). Standard error of the interaction term; 5) P-value for the interaction term; 6) The stratified FDR q-value

http://genome.grid.wayne.edu/SCAIP/Supp_Tables/dynamic_eQTL.tar.gz

**Table S23: table of number of cells per combination (cell-type+treatment+individual** across 1,536 combinations. Columns 1-5 are: 1) combination ID; 2) number of cells per combination; 3) cell-type; 4) treatments; 5) individual ID

http://genome.grid.wayne.edu/SCAIP/Supp_Tables/NumberOfCell_per_combination.txt.gz

